# The plastid-encoded RNA polymerase structures a logistic chain for light-induced photosynthesis

**DOI:** 10.1101/2024.10.25.620210

**Authors:** François-Xavier Gillet, Gregory Effantin, Gian Luca Freiherr von Scholley, Sabine Brugière, Maud Turquand, Noor Pasha, Daphna Fenel, Alicia Vallet, Yohann Couté, David Cobessi, Robert Blanvillain

**Author notes:** co-corresponding authors: François-Xavier Gillet David Cobessi Robert Blanvillain. Univ. Lyon1, CNRS, INSA-Lyon, Bayer SAS, UMR 5240 MAP, 69100 Villeurbanne. Plant Physiology Laboratory, Univ. of Neuchâtel, Rue Emile Argand, 11; 2000 Neuchâtel, Switzerland (Maud Turquand). Université Paris-Saclay, INRAE, CNRS, Univ. Evry, Institute of Plant Sciences Paris-Saclay, 91405 Orsay, France.

## Abstract

The chloroplast is the semi-autonomous organelle of eukaryotes that performs photosynthesis. In higher plants, chloroplast biogenesis depends on a tight transcriptional coordination of both nuclear- and-plastid photosynthesis-associated genes. The plastid-encoded RNA-polymerase (PEP) is composed of a plastid-encoded catalytic core, similar to multi-subunit RNA polymerases, bound to fifteen nuclear-encoded PEP-associated proteins (PAPs). The binding of all the PAPs to the catalytic core is essential for plastid transcription of photosynthesis-associated genes. Our cryo-electron microscopy structure of the native 21-subunit PEP from *Sinapis alba* reveals the distinctive patterning of PAP interactions, which evolved upon the ancestral cyanobacterial catalytic core acting as a scaffold. Using PAP8 *in planta* as bait for affinity purification and proximity labeling, we provide the protein landscapes surrounding the PEP and other PAP8-interacting complexes at the transition from skotomorphogenesis to photomorphogenesis. The data highlight multiple functional couplings in which plastid transcription is at the beginning of a spatial logistic chain, extending from transcription to the assembly of the photosynthetic apparatus into the thylakoids. In addition, dark-specific interactions between photoreceptors and PAP8 establish a physical link between an integrated light signaling and plastid functions.

## Introduction

Chloroplasts are the photosynthetic plastids of the monophyletic lineage including plants, algae and a common ancestor that engulfed a cyanobacterium more than 1.5 billion years ago (Margulis 1975; Archibald et al. 2015). Major lateral gene transfers from the endosymbiont towards the nuclear genome has changed the cyanobacterium into a semi-autonomous organelle (Martin et al. 1998). Consequently, most terrestrial plants possess a reduced plastome with a common set of about 130 genes (Jarvis and Lopez-Juez, 2013). These plastid genes encode components of nearly all functional complexes involved in transcription, translation, protein import, and the photosynthetic apparatus (Puthiyaveetil et al. 2021), in particular, the four prokaryotic subunits (⍺_2_, β, β’, and β’’) corresponding to the catalytic core of the plastid-encoded RNA-polymerase (PEP) (Pfannschmidt et al. 2015). The plastome reduction was permitted by a concomitant evolution of an efficient import system, translocating thousands of nuclear-encoded proteins through recognition of a chloroplast transit peptide (Agne & Kessler 2007). This massive movement of proteins is a major innovation route for newly acquired characters, such as those that occurred during the terrestrialization of the green lineage. For example, a new control of plastid transcription resulted from the acquisition of a nuclear-encoded phage-type RNA polymerase (NEP) expressing plastid housekeeping genes while PEP could specialize mainly in the expression of photosynthesis-associated plastid genes (*PhAPGs*). This transcriptional specificity of the PEP, as observed in higher plants, is now highly dependent on nuclear-encoded PEP-associated proteins (PAPs) that reshape the catalytic core into a highly-active and biochemically stable PEP-PAPs complex (Steiner et al. 2011; Liebers et al. 2020; Ruedas et al. 2022). Most of the PAPs that are singleton genes (PAP3, 5, 7, 11 and 12), display genetic mutants with an albino syndrome characterized by a lack of chlorophylls, defective PEP transcriptional activity, NEP transcriptional compensation, and defect in the construction of the inner membrane thylakoid network. This “albino block” reveals a functional checkpoint in which all PAPs may stabilize the catalytic core increasing the PEP processivity and enabling enzymatic activities such as those providing an appropriate shield against reactive oxygen species (ROS). The cryo-EM analyses of the PEP 3D-structure from *Sinapis alba* (SaPEP) reveal how the PAPs stabilize the PEP architecture by establishing an interaction network as transcriptional cofactors. A nucleo-chloroplastic module of PAPs that accumulate both in the chloroplast and the nucleus has been characterized. PAP5/pTAC12/HMR, PAP7/pTAC14, PAP8/pTAC6, and PAP12/pTAC7 possess nuclear localization signals (NLS) and can interact with each other in the nucleus (Chambon et al., 2022). In Angiosperm the transition from skotomorphogenesis to photomorphogenesis involves a large transcriptional reprogramming triggered by photoreceptors (Hernández-Verdeja & Strand, 2018). Dark-grown seedlings express PAPs only in epidermal cells (Liebers et al. 2018), where the organelles act as sensor plastids (Charuvi et al., 2012; Chan et al., 2016). This pattern of expression may account for the presence of the PEP/PAP during skotomorphogenesis (Ji et al., 2021). However, exposure to light activates the phytochrome B that translocates into the nucleus and coalesces in large photobodies associated with the launching of photomorphogenesis in mesophyll cells. This phytochrome B-mediated light-signaling pathway is dependent on the dually-localized PAP8 and PAP5 for the formation of late photobodies, which is altered in the *pap5/hmr* and *pap8* mutants (Chen et al., 2010; Liebers et al., 2020). The nucleo-chloroplastic module, as part of a retrograde signal, may therefore coordinate the expression of specific nuclear and plastid genes through the photoreceptor signaling pathway. By coupling TurboID-based proximity labeling (PL) and interactomic experiments (Liu et al., 2018) using PAP8 as bait during the dark-to-light transition, we revealed the dynamics of PEP-interacting proteins and uncovered a multi-functional coupling of transcription, translation, and protein homeostasis in the chloroplast as well as a significant relationship of PAP8 with global photoreception in plants.

## Results

### The PEP overall architecture

Our cryo-EM electron density map displays the 21 subunits of SaPEP identified in our mass spectrometry (MS)-based proteomic analyses (Ruedas et al., 2022; Figures 1, S1; Table S1). Its structure is similar to those recently described (Vergara-Cruces et al., 2024; Wu et al., 2024; Do Prado et al., 2024), but our analysis revealed structural features not yet reported. The catalytic core contains two copies of the α subunit that assemble in an asymmetric dimer α_1_/α_2_ (Rpb3/Rpb11 in RNAPII), and one copy of each β (Rpb2 in RNAPII), β’ and β’’ (Rpb1 in RNAPII). The core displays the “crab’s claw” shape observed in multi-subunits RNAPs and is similar to that of the cyanobacteria RNAP (CyRNAP) except for a substantial difference in β’’ (Table S2). As expected in light-exposed samples, the active PEP complex displays nuclear-encoded PAPs (PAP1 to PAP12) and 3 additional subunits FLN2/PAP13, pTAC18/PAP14 and PRIN2/PAP15 (Figure S2) (Ruedas et al., 2022). PAP12 is located where the ω subunit is in the bRNAPs (Rpb6 in RNAPII) (Minakhin et al., 2001). Its structural similarity with the ω of CyRNAP (Table S2) suggests a prokaryotic origin and a lateral gene transfer into the nuclear genome during evolution. Nevertheless, conserved catalytic residues described in bacterial RNAPs (bRNAPs) (Lane & Darst, 2010) are also observed in SaPEP, illustrating the strict conservation of RNA synthesis mechanisms within the chloroplast.

**Figure 1:**
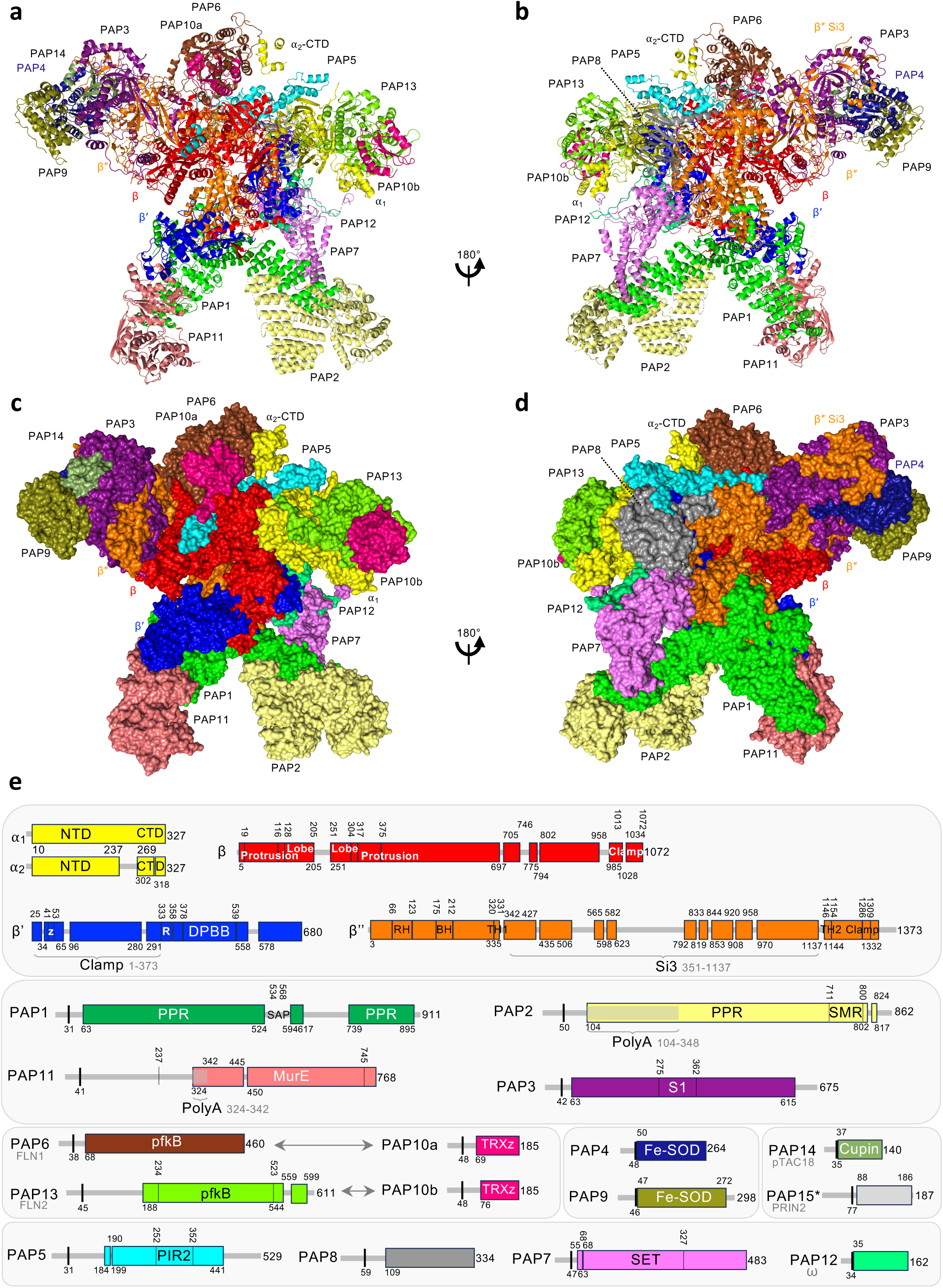
Structure of SaPEP. The complex has a molecular weight of 1.154 MDa and a volume that fits in 253 Å × 180 Å × 204 Å. Each subunit is colored-coded as indicated. (**a-b**) The SaPEP is drawn with the α-helices and β-strands displayed in ribbons and arrows, respectively. (**c-d**) The surface of the subunits is drawn in two orientations. Note that PRIN2 is not drawn in Figure 1a-d due to poor defined electron density connecting subunit β’’ to PRIN2. (**e**) Sequence representation of the subunit’s domains, which are vertically numbered; the observed residues and chloroplast transit peptide (cTP) are horizontally indicated. β displays the classical lobe (Q128-L205/F251-D304) and protrusion (Q19-N116/V317-S375) domains. β’ harbors several catalytic elements (Cramer et al., 2001; Lane & Darst, 2010) well defined in the cryo-EM electron density map: the double-psi-β-barrel domain (Y378-L519), the zipper (Y41-D53) and the rudder (T334-S355). The poorly defined electron density prevented to build the lid (I279-D291) and Swicth2 (V559-D577). β’’ displays the bridge helix (L175-H212) well defined in the cryo-EM map, the rim helices (P66-G89, A93-D123,) that delineate the secondary channel.

The PAPs are grouped in four clusters, all visible from the back of the β’’ subunit; the scarf, the ⍺-cluster, the Si3-cluster, and the tether (Figure 1, S1).

The scarf includes the pentatricopeptides PAP1, PAP2 and the MurE-like protein PAP11 (Figure 2a). PAP1 has twelve-PPR motifs and contains a suspected DNA-binding SAP domain that is not observed in the electron density. PAP1 binds DNA at the same loci as β (Wang et al., 2024) and binds to it as in 8RDY (Vergara-Cruces et al., 2024). PAP1 shares contacts with β’, β’’, PAP11, and the C-terminal SMR (small MutS related) domain (V711 to K800) of PAP2. PAP1 interacts with β’’ in PEP similarly to Rpb5 with Rpb1 in RNAPII, although PAP1 and Rpb5 do not share structural homology. The PAP1/PAP2 heterodimer occupies a position similar to that of the multifunctional Rpb4/Rpb7 in RNAPII (Sharma & Kumari, 2013) (Figure S3). Distal from PAP2, PAP11 is pinched between PAP1 and β’ on the upper part of the clamp, PAP1 and PAP11 being the only PAPs interacting with the clamp. PAP1 also interacts with PAP7 linking the scarf and the tether.

**Figure 2:**
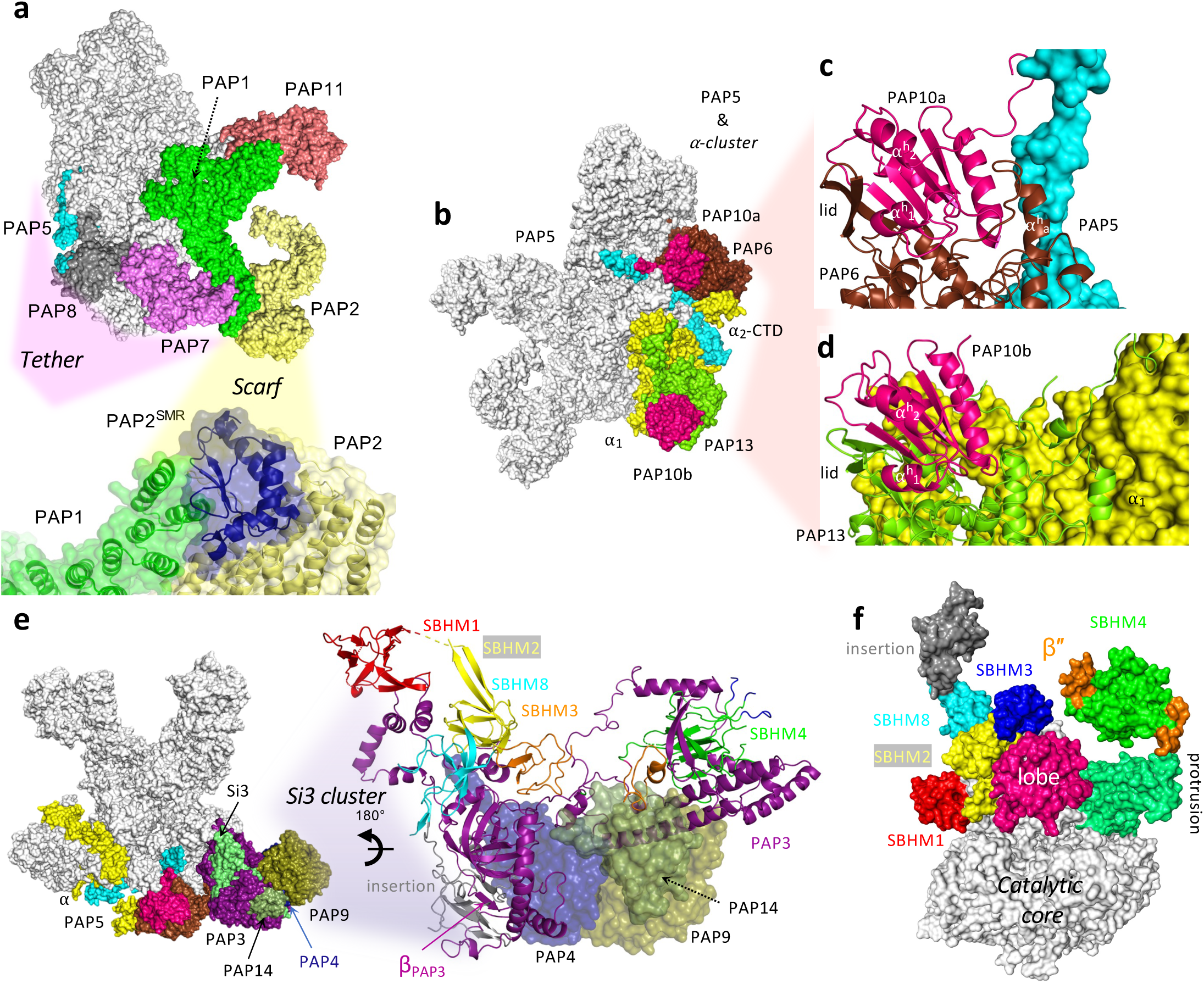
PEP clusters. **(a)** Overview of the scarf cluster (PAP1, PAP2 and PAP11) with PAP5, PAP7 and PAP8 from the tether cluster (see Figure 3). The enlarged view depicts the PAP1-PAP2 interaction with the involvement of the SMR domain of PAP2 in blue. **(b)** Overview of the ⍺-cluster. **(c, d)** Zooms on the PAP6/PAP10a and PAP13/PAP10b heterodimers. PAP6 superposes onto the pfkB with low rmsd (1.52 Å with the pfkB from *Xylella fastidiosa*, PDB entry: 3LKI). Note that two strong peaks of electron density (>12 rmsd) observed in the cryo-EM map were modeled with Ca^2+^ and K^+^ ions by comparison with pfkB from *Vibrio cholerae* (PDB entry: 5YGG, Paul et al., 2018). PAP13 binds a Ca^2+^ ion in a position similar to PAP6, yet it also relies on stabilization by the P246 carbonyl group (P316 in PAP6), influenced by the altered orientation of V107-P136 which includes α_a_. Color legend same as in Figure 1. **(e)** The Si3 cluster, where Si3 is bound to PAP3, PAP4, PAP9 and pTAC18. PAP3 and Si3 are drawn in cartoon with the α-helices displayed as ribbons and the β-strands as arrows. PAP3 is purple colored. Additional domain L970-E1061 is gray colored and folds as an extra-domain of five β-strands in which the β-strand βpap3 of PAP3 inserts. The surface of PAP4, PAP9 and pTAC18 are shown and color-coded as in Figure 1. Si3 is inserted in the trigger-loop between the α-helices V320-P331 and L1146-E1154, and interacts with the lobe and protrusion shown in (**f**). Only five sandwich-barrel-hybrid motifs (SBHM) of Si3 were identified in the cryo-EM map of SaPEP due to its flexibility, preventing to build SBMH5, 6 and 7. The tail, SBHM1 (A347-E427) (red), is along the rim helices. The fin, SBHM2 (K435-L479 and K1105-S1137) (yellow) and SBHM8 (N945-F952 and I1066-A1104) (cyan), is in the front of the rim helix as in SeRNAP. SBHM3 (G482-S506 and Q920-S943) (orange) and SBHM4 (N565-Q582, G598-F618, N792-Y815, V835-D844 and E853-I881) (green) are the body with the non-observed SBMH5 and SBMH7 (Qayyum et al., 2024). The fin and tail are therefore in vicinity of the secondary channel.

The ⍺-cluster includes the thioredoxin-z (TRXz/PAP10) and the B-type phosphofructokinases (pfkB/PAP6 and PAP13) (Gilkerson et al., 2012). The heterodimers (PAP6/PAP10a and PAP13/PAP10b) interact separately with the ⍺-dimer (Figure 2b). PAP10a stabilizes the α_2_-C Terminal Domain (CTD) and sits at the top of the predicted fructose binding site of PAP6 (Figure 2c) gripped by P113-P129 containing the helix α^h^_a_. The “pfkB large lid domain” A192-P213 of PAP6 covers the crevice between the α^h^_1_ and α^h^_2_ helices of TRXz whose catalytic site is oxidized and buried. The second TRXz (PAP10b) forms a heterodimer with PAP13 that interacts only with the α_1_/α_2_ dimer (Figure 2d). PAP13 shares 43.6 % sequence identity with PAP6 and superimposes with a rmsd of 1.23 Å. This superimposition brings PAP10a and the α_2_-CTD in similar positions than PAP10b and α_1_-CTD. As in the PAP6/PAP10a heterodimer, the disulfide bridge Cys108-Cys111 of the WCGPC motif is buried, preventing the formation of intermolecular bridges. Given that PAP6, PAP13, and PAP10a interact with specific regions of the PEP, these pfkBs cannot substitute for one another in stabilizing the PEP architecture.

In the Si3-cluster, PAP3, PAP4, PAP9 and PAP14 interact together and cover the long stretch of β″ called “Sequence insertion 3” (Si3) (Figure 2e, f). Si3 is only found in gram-negative bRNAPs, CyRNAP and PEP (Shen et al., 2023; Qayyum et al., 2024), where it plays a pausing role during transcription in *E. coli* (Kang et al., 2018). In SaPEP, Si3 from T347 to S1137, is the largest part of β’’ containing eight sandwich-barrel-hybrid motifs (SBHMs) and an additional domain in which a β-strand of PAP3 inserts. PAP3 is an S1-protein with non-specific RNA-binding properties (Yu et al. 2018). Within the cluster, PAP4 and PAP9 form an iron superoxide dismutase heterodimer that protects the PEP against ROS (Myouga et al., 2008); while PAP14/pTAC18 is a cupin-like protein of unknown function that displays an eight-stranded β-barrel (Dunwell et al., 2004).

Among the PAPs, PAP3, PAP5 and PAP8 have no structural homologs in the PDB (Figure S4). PAP3 and PAP5 are divided in domains separated by long unfolded stretches interacting with the catalytic core and a large subset of PAPs. PAP5, in particular, interacts with PAP3 and the heterodimer PAP6/PAP10a (Figure 2a, b) linking the different clusters. Outside the chloroplast, PAP5 interacts with phytochrome A (PhyA) through two regions called PIR1 and PIR2 (Galvao et al., 2012). PIR2 (R252-D352) is in the central domain and folds into five α-helices (Figure S4) mainly interacting by one face with the four-stranded β-sheet of α_1_, the other face being exposed to the stroma. PIR1 (M1-W115), however, was not observed in the cryo-EM map due to flexibility.

The tether cluster (Figure 3) includes PAP5, PAP7, PAP8, and PAP12 as a strong partner of PAP7 with a buried surface of 1793.4 Å^2^. These four PAPs possess NLS, and dually-localize in the chloroplast and nucleus (Chambon et al., 2022). PAP7 superimposes onto the histidine-methyltransferase SETD3 (PDB entry: 6ICV; Guo et al., 2019) and histone-lysine N-methyltransferase SETD6 (PDB entry 3QXY; Chang et al., 2011). PAP7 interacts with the α_1_-CTD, PAP12, and its SET domain contacts β’’. Endogenous S-adenosylmethionine bound to PAP7 suggests its activity as a *bona fide* methyltransferase (Figure 3f). PAP7/PAP8 form the tether which connects the upper to the lower claw of the catalytic core by interacting with PAP1 on one side and PAP5 on the other side (Figures 2a, 3). PAP8 is located at the front of the secondary channel in a similar position to that of Rpb8 in RNAPII (Figure S3). While Rpb8 interacts non-specifically with ssDNA (Kang et al., 2006), PAP8 displays a non-specific RNA-binding activity *in vitro* (Chambon et al., 2022). PAP8 interacts with the rim helices, β’, PAP5, and PAP7. The 3-stranded β-sheet of the PAP8 C-terminal part interacts with the β-strand V187-Y189 of PAP5 forming an antiparallel 4-stranded β-sheet. This 4-stranded β-sheet interacts with the 5-stranded β-sheet (L602-T654) of β’ in a 9-mixed stranded β-sheet. Moreover, the three last turns of the PAP8 C-terminal α-helix interacts with the three first turns of the kinked α-helix of PAP5 (Figure 3e). Thus, taken together, the PAP5-PAP8 buried surface is 1415.4 Å^2^. Combined to our previous NMR studies showing interactions between the PAP8 globular domain and PAP5, this heterodimer may also occur outside of the chloroplast *e.g.* in a PAP nuclear subcomplex as previously observed (Liebers et al., 2020; Chambon et al., 2022). Within the globular domain of PAP8, D175 and D177 form hydrogen bonds with L238 and R242 of PAP7 and the loop D211-R214 interacts with the SET domain of PAP7 with an overall buried surface of 411.8 Å^2^. It is therefore possible that outside the PEP the NLS-bearing proteins of the tether cluster form a sub-complex involving other proteins such as PhyA thru PIR1 and PIR2 domains of PAP5 and possibly the targets of the methyltransferase PAP7. The first 49 residues of PAP8 are mainly Gly, Ala, Asp and Glu, that are not observed in the electron density but accessible in the stroma. The PAP8 flexibility described from 2D-NMR studies (Liebers et al., 2020) combined to the position of PAP8 in the PEP were keystones for our AP/PL studies.

**Figure 3:**
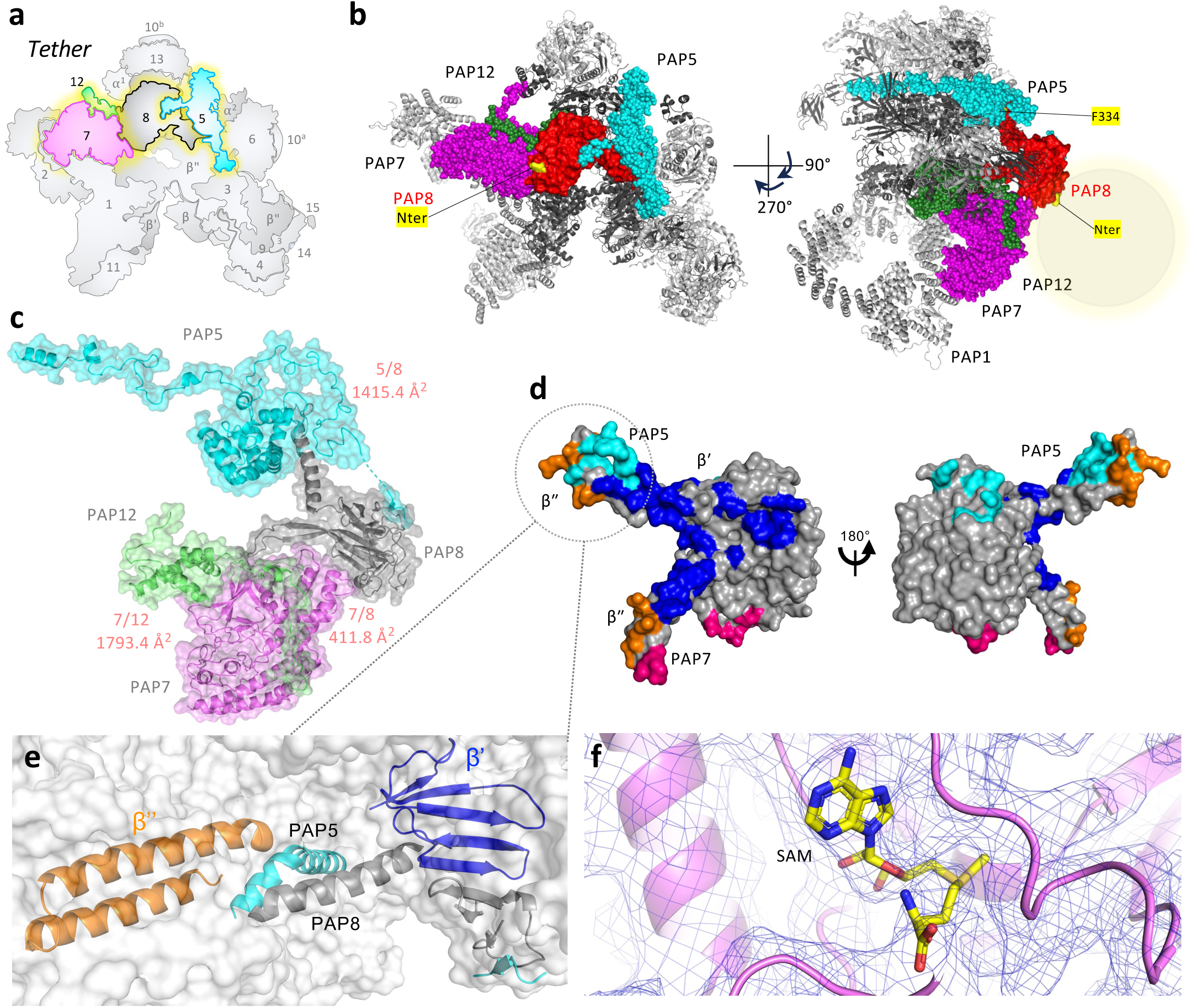
The tether cluster. (**a**). Topology of the tether cluster comprising PAP7, PAP8, extended to a larger Nucleo-chloroplastic module with PAP5 and PAP12. (**b**) Two views of the module within the PEP: PAP5, 7, 8 and 12 are drawn in spheres and colored as in Figure 1 except PAP8 in red. The catalytic core and the other PAPs are colored in dark grey and light grey, respectively. The first and last visible residues of PAP8 in the cryo-EM electron density map are highlighted in yellow (Nter and F334). The gray disc at the N-terminal shows the open space for the TurboID fusion. The steric hindrance surrounding PAP8-F334 prevented C-terminal functional fusions. **(c)** The isolated tether cluster as observed in the SaPEP complex with the buried surface calculated using PISA from CCP4. (**d**) Two views of PAP8 surfaces, colored according to the protein it interacts with: as a bifid structure PAP8 in gray displays 2 zones of interactions with PAP5 in cyan and two others with PAP7 in magenta. (**e**) Interactions between β’, β’’, PAP5 and PAP8. The rim helices are orange colored. The β-sheet of β’, PAP8 and the β-strand of PAP5 involved in the nine-stranded β-sheet are colored in blue, gray and cyan respectively. (**f**) View of the SAM bound to PAP7 superimposed onto the cryo-EM electron density map. The electron density map is blue colored.

### Set-up of PAP8 tagged versions to study partnership dynamics

Our structural analysis of the PEP raised questions regarding the roles of the PAPs at the transition from dark (D) to light (L). This transition triggers a significant remodeling of the prokaryotic PEP catalytic core, which supports the reprogramming of plastid gene expression (PGE). To explore the dynamics of interactions between PEP and plastidial proteins during the initial D-to-L transition, PAP8 was used as a bait for TurboID-based proximity labeling (PL; Branon et al., 2018) and affinity purification (AP) coupled to MS (Figures 4, S5). PAP8 was selected for its ability to localize in both chloroplasts and nuclei (Liebers et al., 2020), following an unknown pathway that could potentially be mapped in PL.

**Figure 4:**
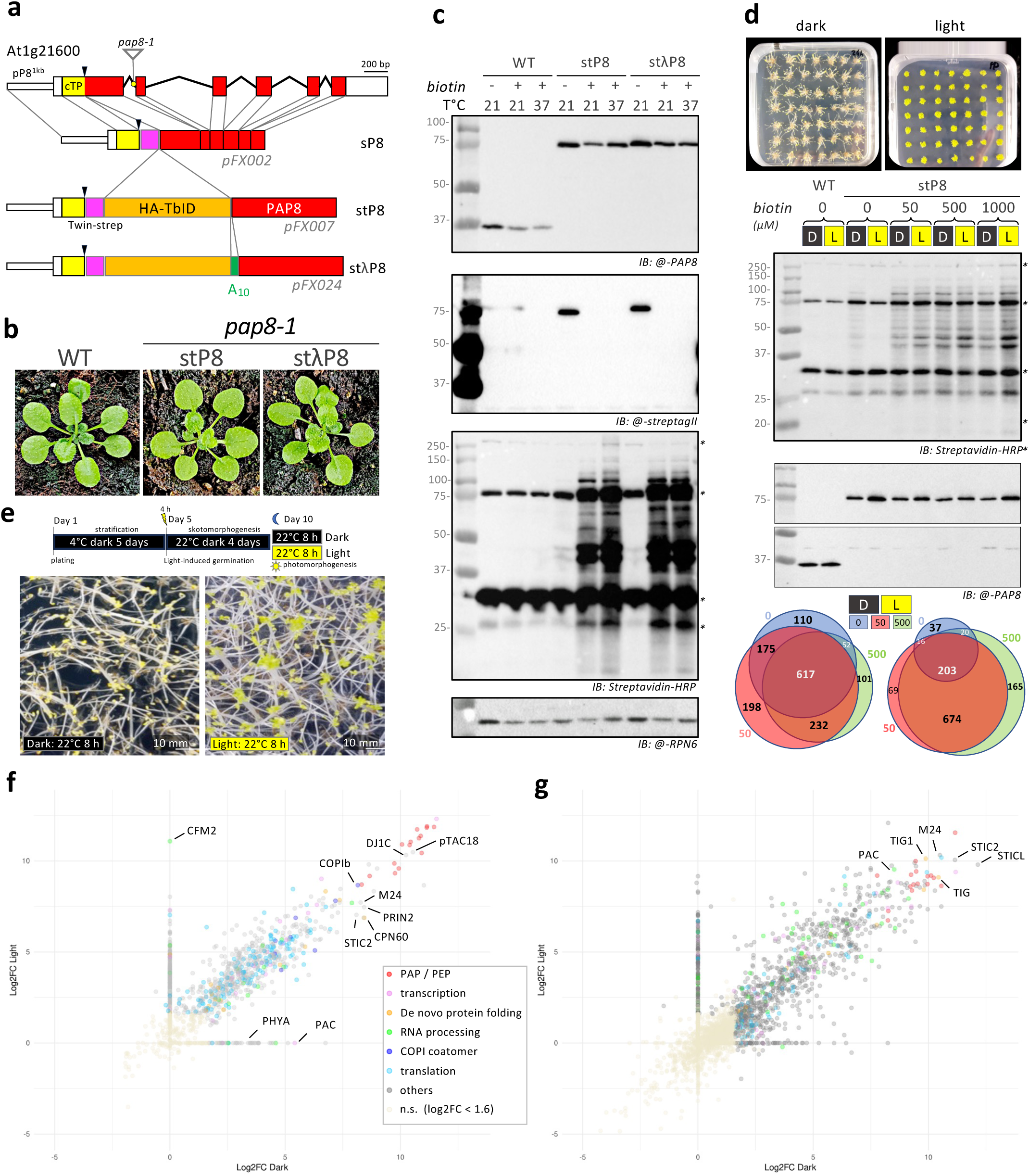
TurboID-Strep-tagged PAP8 restores the mutant phenotype of *pap8-1* and exhibits biotinylation activity at the transition from skotomorphogenesis to photomorphogenesis in *Arabidopsis*. **(a)** Genetic background of the experimental setup. pP8^1kb^ (PAP8 5’ region of 1180 bp), cTP^PAP8^(PAP8 predicted chloroplast transit peptide), twinstrep, hemagglutinin fused to TurboID (HA-TbID)^t^, the poly-ALA (A_10_)^λ^ used as linker, and the conceptually processed PAP8 DNA fragments were assembled into sP8, stP8 and stλP8 constructions used for transient and stable transformation of *N. benthamiana* and *A. thaliana pap8-1*/+ heterozygotes. **(b)** Phenotypes of wild type (WT), stP8 and stλP8 (T3 generation doubly-homozygous for the transgene and the *pap8-1* allele) at the rosette stage; a difference from WT was not observed during this phase or the reproductive phase. **(c)** Expression and activity of the TbID constructs were analyzed by immunoblots (IB) with anti-PAP8 antibodies, @-streptagII (2xStrep detection), streptavidin-HRP (biotinylated protein detection) and anti-RPN6 antibodies as loading control. Biotin was added at 21°C or 37°C and compared to no biotin control at 21°C. Asterisks mark the positions of naturally biotinylated proteins. **(d)** Seedlings *in vitro* production and biotin ligase activity at the transition from dark (D) to light (L) with a biotin-dose response; IB as in (**c**); and Venn diagram of proteins detected in MS for the different concentrations of biotin in dark and light samples. (e) *In vitro* production of seedlings at the transition from skoto-morphogenesis (dark grown hypocotyls with a length of 10 to 15 mm) to photomorphogenesis (samples collected after 8 hours of white light exposure). (**f, g)** Global Dark-to-Light comparison of stP8 affinity purification **(f)** and proximity labeling **(g)**. Dots represent the Log2 Fold changes in Dark (x-axis) and Light (y-axis) for each detected protein, color code as indicated.

To insure proper physiological context, we utilized the *pap8-1* mutant in which PAP8 translational fusions could complement the albino phenotype. Knowing that PAP8 cTP can be separated from its processed amino acid stretch by a GFP fusion without altering its function (Chambon et al., 2022), a similar molecular approach was developed (cTP8-2xStrep-HA-TurboID-PAP8) under the control of 1.2 kb 5’ upstream region of the PAP8 CDS referred as promoter “pP8” (Liebers et al., 2018). Three fusions were cloned without HA-TurboID (sP8) or with HA-TurboID but with or without a 10-alanine linker λ, stP8 and stλP8 respectively (Figure 4a, S5). sP8 was used for a strep-based AP of PAP8-containing complexes *in planta*, while stP8 and stλP8 were used in parallel for PL and AP experiments carried out in the *pap8-1* homozygous background in which the greening process was restored by the corresponding transgene of interest (Figure 4b). stP8 and stλP8 were immunodetected at their calculated molecular weights (75/76 kDa) without any detectable cleavage products showing the stability of the recombinant protein. The absence of PAP8 at 37 kDa confirms that the recombinant proteins are produced in the *pap8-1* homozygote background (Figure 4c). The amount of the chimeric proteins is comparable or slightly superior to that of the wildtype PAP8.

### Characterization of the AtPEP

We studied the PEP-enriched fraction of the sP8/*pap8-1 Arabidopsis* line using MS-based quantitative proteomics. The ranking of the identified proteins using the intensity-based absolute quantification iBAQ metrics (Schwanhäusser et al., 2011), showed that the analyzed fraction was strongly enriched in PEP components. The four core subunits (⍺, β, β′, β″) and the twelve PAPs (PAP1-12) were among the 18 most abundant proteins and represented 65% of the total amounts of proteins within the fraction (Figure S6, DataSet1_AtPEP_MS.xlsx). Among the abundant proteins, we identified CSP41b (Qi et al., 2012) and PRIN2 (Kremnev & Strand, 2014; Díaz et al., 2018) required for proper greening of the seedling, as well as FLN2 and pTAC18, also identified in the SaPEP 3D-structure. Our MS-based proteomic analysis also detected significant levels of the light-harvesting complex subunit LHCB1.1, ribosome, and ribosome-associated proteins. Other components identified in the MS analysis belong to a set of proteins found in all samples (stP8 and stλP8) treated below such as AtRABE1B, CPN60b2, DJ1C and pTAC13. The 3D envelope of AtPEP purified from sP8 was calculated at 31 Å resolution (Figure S6b), and is similar to that of SaPEP (EMDB entry: EMD-14571) (Ruedas et al., 2022).

### Detailed analysis of stP8 and stλP8 interactomes and proxisomes

The study of the PEP protein dynamics in early development requires large quantities of seeds. Therefore, optimization of the PL approach was first conducted with the application of exogenous biotin, testing temperature and dosage (Figure 4d). At 37°C, protein biotinylation was slightly increased compared to 21°C. This is consistent with the behavior expected for enzymes from intestinal bacteria, which catalyze reactions at 37°C. However, this was not chosen to avoid the high-temperature stress response of the plant. Exogenous biotin significantly increases the number of biotinylated proteins and is particularly important in light when the seedling becomes metabolically very active. TurboID is therefore functional and optimal with 50-µM supply of biotin. Quadruplicate experiments were then performed on wild type, stP8, and stλP8 for both AP and PL under dark (4-day old dark-grown seedlings) and light conditions (4-day-old dark-grown seedlings + 8-h of white light treatment) (Figures 4e-g, S7). Comparison of stP8 and stλP8 with wild type generated 8 datasets, (AP/PL) × (stP8/stλP8) × (D/L) (DataSet2_stP8stlP8APPL.xlsx), presented as volcano plots in Figure S7. Among the 4337 TAIR proteins reliably identified and quantified, 2086 were significantly enriched with PAP8 compared to negative control in AP or PL experiments. Among them, proteins from the catalytic core (red), PAPs (orange), and accessions common to all sets (blue) were distinguished, ranked according to their log2(Fold Changes (FC)) values and grouped in functional categories (Dataset2b_allSig.xlsx). A striking graphical difference lays on the appearance of a top cluster composed of the PEP subunits in AP while embedded with many more proteins of similar enrichment (Figures 4f, g, S7) or abundance (Figure 5) in PL. This attests for a true difference between AP and PL approaches and shows that a stable PEP can be purified in AP with the labeled PAP8 bait. The repartition of these 2086 proteins (Figures 5a, S8) allows for the identification of 107 common proteins present in all sets including all the PEP subunits observed in the SaPEP structure (DataSet2c_ComRank.xlsx). Global subcellular localization analysis indicates that the highest-ranked proteins are predominantly plastid-localized. However, proteins from other compartments are increasingly represented further down the list, revealing non-chloroplastic partners. Gene ontology analyses (Figure S9) reveal that the preponderant groups are associated with transcription and PGE. Yet other biosynthetic pathways are well-represented indicating the possibility of functional couplings.

**Figure 5:**
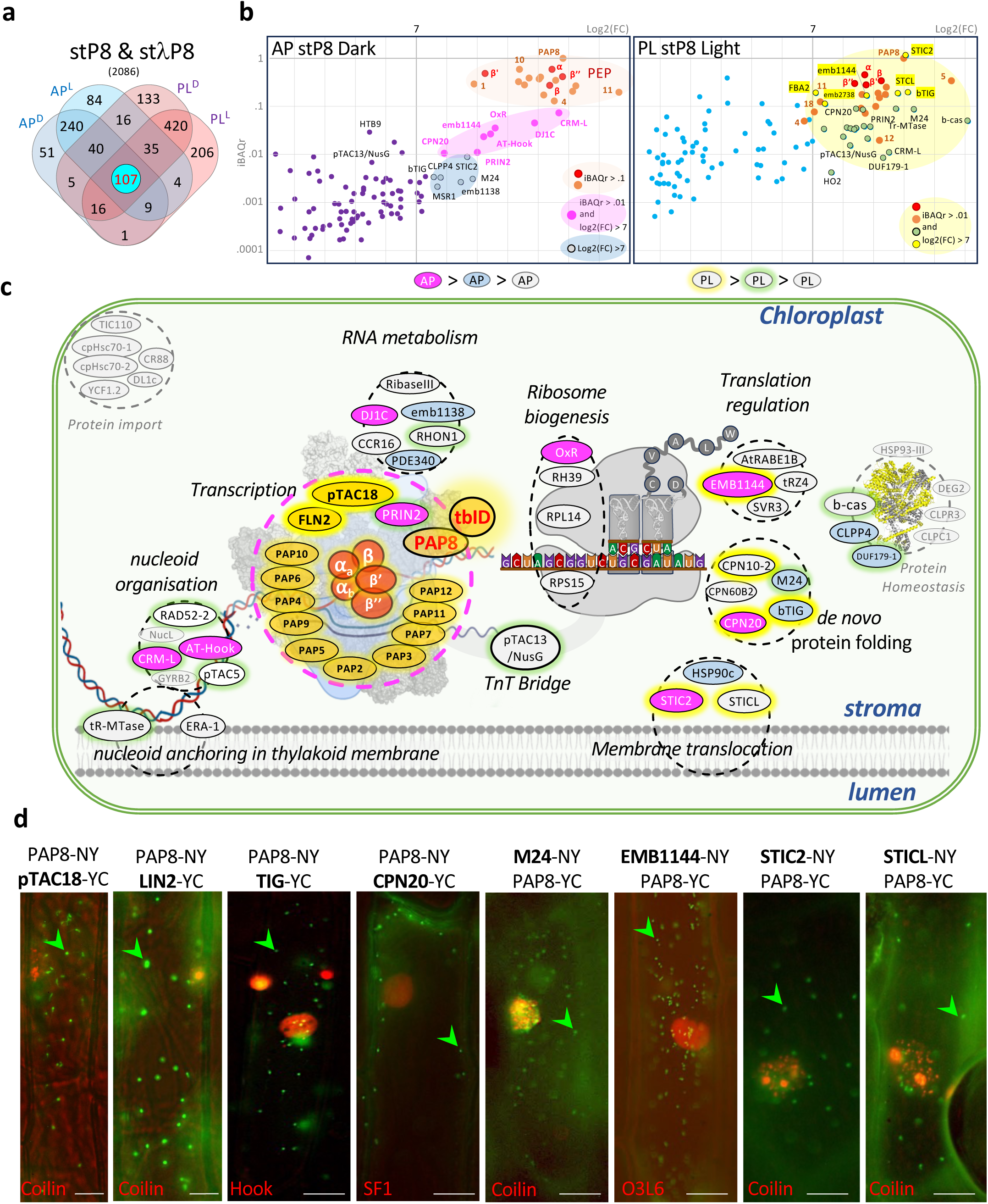
Analysis of the proteins common to all sets (AP/PL, D/L, stP8/stλP8) reveals a structurally organized logistic chain for plastid gene expression from anchoring to membrane translocation including the prokaryotic type TnT coupling. **(a)** The Venn diagram shows the distribution of the 2086 individual candidates within the stP8 sets (1755) or stλP8 sets (1787) or both (2086). **(b)** Plots of iBAQr=f(Log2FC), iBAQr is relative to the highest iBAQ from the sample, for two representative datasets AP stP8 Dark and PL stP8 Light (see Figure S10 for full set). **(c)** Schematic representation of the logistic chain associated with PEP-dependent PGE. Plastid DNA anchored to the thylakoid membrane going thru nucleoid organization then entering the PEP complex (PDB entry: 9EPC). TnT transcription and translation coupling represented by pTAC13/NusG acting as a bridge between the functional complexes. The ribosome schematically represented as the small and large subunits displaying A and P sites filled with tRNAs. The elongating aa chain is represented with translational regulators in the way to the *de novo* protein folding mainly involved in successful membrane translocation or degradation by the CLP-mediated protein homeostasis module (PDB entry: 7EKO (Wang et al., 2021)) with yellow parts corresponding to AP/PL detected proteins. **(d)**. BiFC test for several PAP8 partners within the common set of proteins. NY and YC correspond to the N-terminal and C-terminal parts of YFP respectively fused to the tested protein. Coilin, Hook, SF1 and O3L6 fused to DsRed are transfection controls (Blanvillain et al., 2009); green arrowheads point to plastids; bars = 20 µm.

According to a similar ability to restore PAP8 functions in *pap8-1,* stP8 and stλP8 datasets are very similar for the most enriched proteins. To minimize spurious discoveries, we disregarded the proteins found enriched with a lower FC. As expected though, the “linker” could account for longer range biotinylation, hence increasing the protein pools in PL experiments (+3% in D to +31 % in L) but not in AP where an opposite trend is observed (-13% in D to -50% in L) suggesting that a long linker might destabilize complexes, no further attention given. Analysis of the FCs computed for the proteins found enriched in AP and PL experiments shows that both experiments yield different results; proteins with strongest FC in AP are not necessarily the most differential ones in PL as illustrated with PAP11 that sits on the clamp far away from PAP8. The reverse situation is observed, for example with PAP5. Stoichiometric analysis using iBAQ relative to PAP8 was conducted on all datasets (Figure S10; DataSet2d_Stoichiometry.xls) for which representative AP/stP8/D and PL/stP8/L are given (Figure 5b). In the categories with the highest abundances, proteins can be classified into groups that align with a logistic chain of plastid gene expression, highlighting a strong functional coupling between membrane-anchoring, transcription, translation, and translocation (Figure 5c). Protein import and homeostasis, especially the CLP-chaperon proteases, were also represented in the common proteins. Some of these proteins (pTAC18, TIG, CPN20, M24, EMB1144, STIC2, STICL, and LIN2) were tested in bimolecular fluorescent complementation experiments using the split YFP system with PAP8 (Figure 5d). All of the observed interactions mark the plastids within the cell, corroborating the *in vivo* proximity of PAP8 with the tested proteins. Two proteins from the common set are predicted to be present in the nucleus AT-Hook (AT1G48610) and HTB9 (AT3G45980). Although AT-Hook is also predicted to localize in plastids (SUBA5/chloroP) and potentially present in the nucleoid, the detection of the histone 2B (HTB9) is intriguing. This specific histone indeed, is involved through ubiquitination in the rapid chromatin-based modulation of gene expression at the transition from dark to light (Bourbousse et al., 2012). PAP8 or the PAP nuclear sub-complex may therefore play a role in this process.

### Modulation of PAP8 partnership in Dark versus Light conditions

The comparison of the dark and light samples provides insight in the dynamics of the PEP protein partners at the transition from skotomorphogenesis to photomorphogenesis. The light-enriched protein pool contains many proteins from the photosynthetic apparatus (DataSet2e_Light_Components_Rank.xlsx). Proteins were sorted in three categories according to their mode of detection (AP, PL, or both) (Figure S11) and highlighted in the different PS complexes (Figures 6, S12). Strikingly AP-detected proteins belong mostly to the PSII and PL-detected proteins belong to the NDH complex. We conclude that the PEP complex is anchored on the thylakoid by interacting at least with PSII subunits (Figures 6a, S12). Such physical interaction is consistent with previous observations showing that PAP8 antibody recognizes a specific molecular complex in the thylakoid fraction of the chloroplast (Chambon et al., 2022). The strong differences observed between AP and PL for these partners may result from a PEP positional effect, masking reproducibly some surfaces while others are exposed knowing that stP8 is close to the secondary channel likely facing an open space of the stroma. This would easily mark with biotin unlinked proteins such as the NDH subunits (NdhM, K, S). One side of the ATP synthase (ATPF, ATPA) interacts with the PEP, while the more central ATPB and D are prone to biotinylation. This suggests that some interactions between the different photosynthetic complexes and the PEP are more frequent than others during the initial steps of the building of the photosynthetic apparatus. This occurs when the grana stacks are not yet formed and the membranes are highly dynamic.

**Figure 6:**
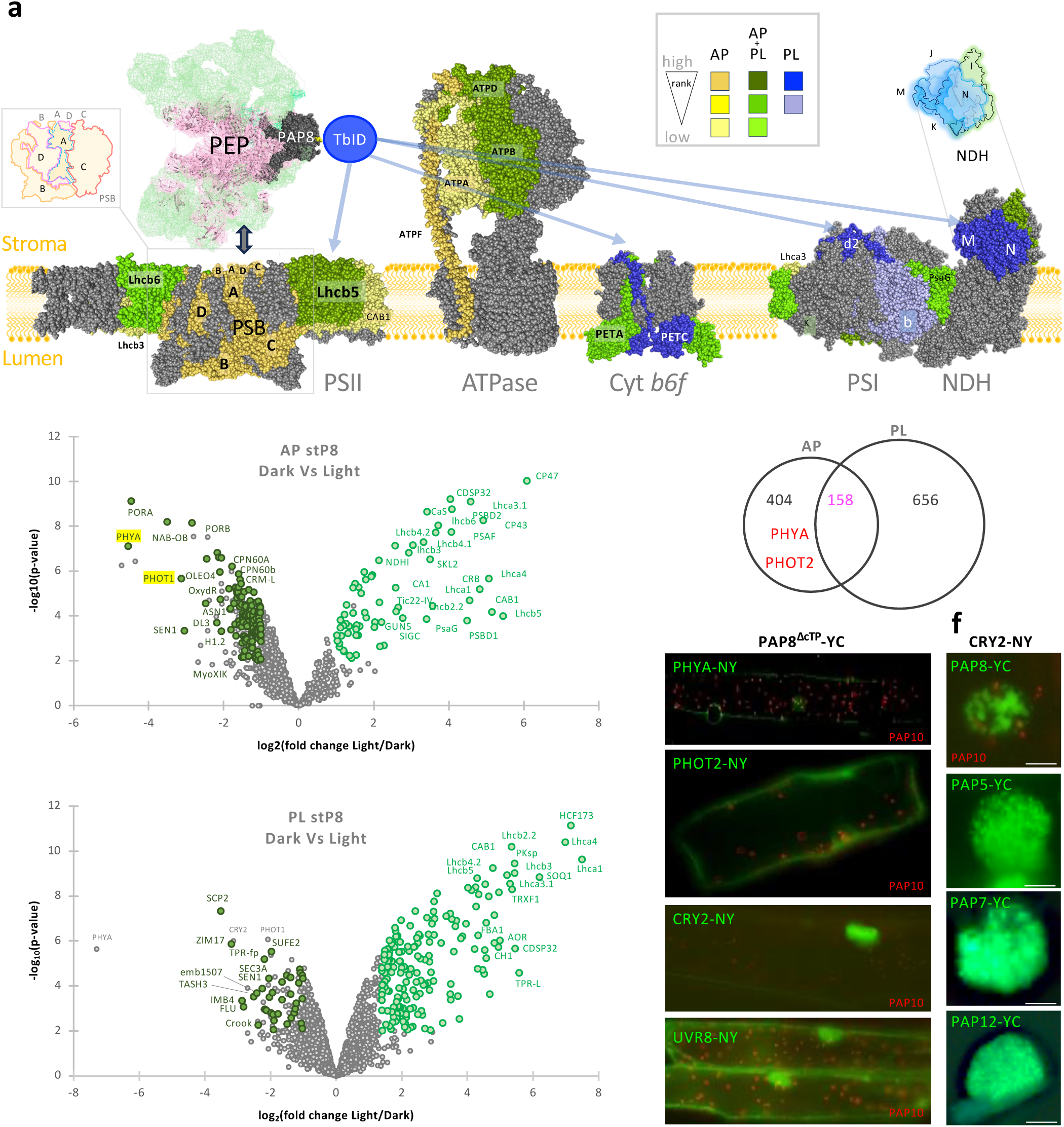
Light versus dark comparison of the data reveals an intimate relationship of the PEP with PSII after light exposure, while PAP8 interacts with photoreceptors, outside of the chloroplast, during skotomorphogenesis. **(a)** PSII (PDB entry: 7OUI, (Graça et al., 2021)), Cyt b6f (PDB entry: 6RQF (Malone et al., 2029)), PSI/NDH (PDB entry: 7WG5 (Su et al., 2022)), and ATPase (PDB entry: 6FKF (Hahn et al., 2018)), were color coded according to the rank analysis of light enriched components of the photosynthetic apparatus (Figure S11 table S6). Box: color codes: yellow blue and green represent respectively AP, PL, or, AP+PL-enriched subunits, shades represent the ranking of the protein in the dataset. **(b, c)** Volcano plots of the “light versus dark” enriched proteomes obtained in AP (b) or PL (c). **(d)** Venn diagram of the repartition of the Dark-enriched proteins between AP and PL samples. **(e)** BiFC test of the interactions between PAP8 and the photoreceptors PHYA PHOT2 CRY2 and UVR8. NY and YC correspond to the N- and C-terminal parts of YFP respectively fused to the photoreceptor as indicated or the ΔcTP version of PAP8. PAP10 fused to DsRed is used as a transfection control. **(f)** In the CRY2 column the ΔcTP versions of PAP8, 5, 7, and 12 were tested with CRY2-NY. Pictures were taken as a closeup on the nucleus photobodies; bars = 10 µm.

Sigma factors play crucial roles in the regulation of PGE by directing the PEP to specific promoters. In particular, SIG2 and SIG6/SIGF are essential for proper chloroplast biogenesis as their corresponding mutants are pale green (Woodson et al., 2013). Accordingly, SIG2 and SIG6 are the two predominant σ-factors found to be particularly enriched in AP_MS experiments, in dark and light conditions. Their weak detection in PL-MS experiments likely reveals the lack of accessibility to their residues.

### Binding of σ-factor, NusG, DNA and 70S ribosome to the PEP

Our AP/PL studies revealed NusG/pTAC13, SIG2 and ribosomal proteins as potential partners of the PEP. We thus, modeled the PGE initiation step by superimposition of SaPEP onto SyRNAP bound to the σ-factor SIGA and DNA (PDB entry: 8GZH; Shen et al., 2023), and onto SIG2 from *A. thaliana* (AF-O22056-F1). The modeling shows SIG2 interacting with PAP11, and the DNA with SIG2 in contact with the PAP2 N-terminal part (Figure S13). The Si3-σ arch structure cannot be observed since the Si3 head is not defined in the cryo-EM map. Stabilization of chloroplastic sigma factors with DNA during initiation stage may therefore involve interactions with PAP2 and PAP11. Modeling reveals also SIG2 in the vicinity of the C-terminal part of PAP5 for which the residues 442-529 were not built. In the initiation stage, the PAP5 C-terminal part may stabilize the sigma factor. The comparison between SaPEP and SeRNAP (Qayyum et al., 2024) reveals that SaPEP may bind both DNA and NusG/pTAC13 (AlphaFold model AF-Q94AA3-F1; Jumper et al., 2021) without any large conformational change. In prokaryotes NusG links the ribosome with the transcription elongation complex providing the structural framework of the transcription-translation (TnT) coupling (Washburn et al., 2020). Since TnT coupling involving the PEP is highly considered in chloroplast (Zoschke & Bock, 2018), we also modeled a complex between SaPEP and the 70S chlororibosome from *S. oleracea* (PDB entry: 5MMM; Bieri et al., 2017) based on structure of the *E. coli* expressome (PDB entry: 6ZTL; Webster et al., 2020). Model analyses showed clashes mainly between the 30S subunit, PAP2, PAP10a and PAP11. Conformational changes must therefore occur to allow interactions similar to those observed in *E. coli*.

### PAP8 associates with photoreceptors in the dark

When comparing AP-MS in dark and light conditions, the photoreceptors phytochrome A and phototropin appear in the top list of PAP8-associated proteins. CRY2 and UVR8 were also detected but not significantly enriched compared to the wild-type control (Figures 6b-c, S14). Nevertheless, bimolecular fluorescent complementation studies using the Δ-cTP version of PAP8 confirmed these interactions *in planta* (Figure 6e). Fluorescent patterns were restricted to the cytoplasm with PHOT2. A faint signal was observed in the cytosol and around the nucleus with PHYA, whereas strong signals were detected in the nucleus with CRY2 and UVR8. These results are in agreement with the pattern of each photoreceptor fused to GFP (Figure S15). PAP8 may therefore act as an integrator of a photoreception hub including all major photoreceptors. With CRY2 a signal was also observed using PAP5, 7 and 12 according to a speckling pattern resembling to a phase separation of small and numerous photobodies (Figure 6f). The dually localized PAPs are able to interact with CRY2, not merely as individual proteins but likely as a larger PAP nuclear sub-complex, whose 3D structure may resemble the tether cluster given in Figure 3c.

## Discussion

Our previous studies, together with results presented here, revealed a 3D envelop of SaPEP similar to that of AtPEP purified with a tween-strep affinity (Figure S6). This opens the road for structural analyses of *Arabidopsis* plastid transcription with the tools of genetics. The cryo-EM structure of SaPEP highlights both differences and analogies with the multi-subunit RNAPs, underscoring the role of the PAPs in PEP stability, transcription and the overall PGE. SaPEP, like other multi-subunit RNAPs, share a similar catalytic core. Its α, β, β’ and β’’ subunits closely resemble those found in the CyRNAPs. Interestingly, the CyRNAPs ω subunit is replaced by the nuclear-encoded PAP12, with structural similarities. PAP12 may correspond to an ancestral cyanobacterial ω gene transferred into the nucleus and extensively modified, but other evolutionary scenarios, such as lateral gene transfer or neofunctionalization, are also possible. Comparison with RNAPII revealed that PAP8 occupies a position analogous to Rpb8; while PAP1 and PAP2 are positioned similarly to Rpb4/Rpb7. This suggests that these subunits may have similar roles despite their different folds (Figure S3). The PAPs do not bind to essential catalytic elements such as the lid, the ruder, the zipper or any other catalytic residues conserved across prokaryotes. Thus, the catalytic mechanism of the PEP and CyRNAPs are likely similar. The PEP is therefore a unique RNAP built on the CyRNAP scaffolding without the ω subunit, *i.e.* PAP12. In the context of chloroplast biogenesis, the PAPs bring enzymatic activities and stability essential to the transcription in a compartmentalized organelle, the mandatory presence of PAP being well highlighted by the *pap* mutant albino phenotypes. PAP12 substitutes for the ω subunit to stabilize the assembly. PAP6/PAP10a and PAP13/PAP10b stabilize the α-CTD of α_2_ and α_1_ respectively where transcriptional cofactors bind in bacteria. The flexibility of α subunits is also observed in bRNAPs. The α_1_/α_2_ dimer serves as a crucial scaffold for the assembly of the catalytic core in bRNAPs. Additionally, it plays a significant role in transcription activation by interacting through its CTDs with transcription factors and an AT-rich DNA sequence located upstream of the - 35 element in strong *E. coli* promoters. (Sutherland and Murakami, 2018). PAP3 sandwiches the additional domain of Si3 unique to the plant β’’ subunit, shielding it. PAP1 binds to DNA (Vergara-Cruces et al., 2024; Wang et al., 2024) and PAP2 binds RNA (Wu et al., 2024). PAP4 and PAP9 are SODs that protect the PEP against the ROS (Myouga et al. 2008). PAP8 occupies a similar position to that of Rpb8 in RNAPII and may be involved in RNA binding (Chambon et al., 2022). PAP7 bound to S-adenosylmethionine likely exhibits methyltransferase activity. It is the sole PAP to interact with PAP12. PAP11 and PAP2 may interact with sigma factors and the PAP2 N-terminal may also interact with DNA during transcription initiation. The TADs of PAP15 may interact with transcriptional cofactors. The PAP clusters interact with, and wrap around, the catalytic core to stabilize the PEP architecture. Structural studies of the PEP at various stages of transcription will elucidate the precise roles of PAPs.

### Dynamics of the PEP interacting proteins at the transition from dark to light

Our AP/PL-based study highlighted protein dynamics surrounding the PEP at the developmental transition from dark to light and revealed a common set of proteins that are invariable PAP8 interacting and fleeting partners. However, according to the transcriptional regulation of PAP8, it can be inferred that complexes from dark and light samples correspond to distinct cell identities within the cotyledon. Specifically, dark datasets predominantly display protein partners from epidermal cells containing sensory plastids (Mackenzie & Mullineaux, 2022) whereas light datasets predominantly include proteins from chlorophyllous palisade tissue. (Liebers et al., 2018). The epidermal cells may therefore contain an active PEP complex conducting an epidermoplast-specific transcription. Alternatively, additional mechanisms of pausing or repression such as those observed in the coupling of genomes prevents the PEP from producing the photosynthetic apparatus in a light-specific and/or cell-type specific manner. Such pausing mechanism involving PAP8/pTAC6 and termination factors mTERF proteins have already been identified (Ding et al., 2019; Méteignier et al., 2020). mTERF5/MDA1 in particular (global rank 1632) was exclusively found in stλP8 PL Dark, which however indicates that these regulations are likely not preponderant during the initial phases of chloroplast development.

The trafficking of PAP8 between the different compartments, chloroplast and nucleus in particular, akin to other dually-localized proteins (Whirly, PAP5/HMR, NCP, RCB), remains a topic of debate (Krause & Krupinska, 2009; Chen et al., 2010; Kleine & Leister, 2016; Yang et al.,2019). stP8 and stλP8 may conceptually leave biotin-mark along its trafficking route allowing for the identification of partners encountered along the journey. Strikingly, gene ontology analysis reveals the presence of several subunits of the coatomer especially those belonging to the COPI complex in the top list of several datasets, mainly APD and APL (Figure S7, dataset2b: subunits: ε_1_; ⍺_2_; β_1_; and ⍺_1_). This raises the question of a possible vesicular transport for PAP8 (Jürgens 2004; Ahn et al., 2015 for reviews). Whether this vesicular transport is associated with the retrograde movement of PAP8 towards the nucleus *via* the endoplasmic reticulum, part of a lytic pathway, associated to the trimeric assembly of the triad (Gomez-Navarro & Miller, 2016) or disconnected, remains to be determined. Should PAP8 circulate in COPI-coated vesicles, it is plausible that PL would be significantly hindered by the vesicular coating as suggested by our datasets.

### Logistic chain

Most proteins found across all datasets belong to distinct functional complexes but clearly participate in PGE. Indeed, PAP8 used as bait may interact, directly or not, with the isolated candidates. A strong anchoring of the light-activated PEP complex on the thylakoid, as previously described (Powikrowska et al., 2014), should help absorb the physical forces of transcription, allowing for the production of the transcript in a control space for rapid building of the photosynthetic apparatus. Specifically, proteins such as pTAC5 interacting with heat shock proteins (Zhong et al., 2013), or CRM-L and RAD52-2 involved in chromosomal maintenance and repair; or ERA-1 and tR-MTase found in the thylakoid and in the RNA-binding proteome (Bach-Pages et al., 2020) may provide such anchoring of the nucleoid to the thylakoid membrane. This functional anchoring has been recently proposed (Palomar et al., 2024). By comparison with prokaryotes, NusG/pTAC13 may ensure proper positioning of the PEP with the small subunit of the ribosome to efficiently load the transcripts into the ribosome. This TnT functional coupling is extended to other PEP partners in RNA metabolism and ribosome biogenesis such as emb1138 (RH3, Asakura, et al., 2012) and RH39 (Nishimura et al., 2010) RNA helicases of the DEAD-box superfamily. These enzymes hydrolyze ATP to modify RNA structures such as splicing group II introns (emb1138/RH3), or generating hidden breaks for ribosomal RNAs maturation (RH39). TnT of the PEP extends to translational and post-translational regulation with other proteins involved in aromatic amino acid biosynthesis (EMB1144), in the proper folding of nascent polypeptides such as the trigger factor (Rohr et al., 2019) and associated chaperonin cofactors (CPN20, CPN10). Furthermore, this extends to membrane translocation involving STIC2 and STICL, associated with protein homeostasis and the inner side of the translocon (Bédard et al., 2017). After light exposure and activation of the PEP in the palisade, the anchoring of the PEP to the photosynthetic complexes occurs at the site of the *de novo* protein translocations (Figure 6a).

This study marks a significant step toward understanding the evolutive innovations brought by the PAPs to optimize the building of the chloroplast upon light exposure. Although numerous other proteins were detected in the study, the discovery of interactions with multiple photoreceptors surpassed expectations. PAP8, with the other dually-localized PAPs, may interact together with all photoreceptors in a framework of an integrated response to the full spectrum of light from ultra-violet to far-red (Kami et al. 2010). This could coordinate nuclear and plastid gene expression in processes such as shade avoidance, photoprotection, and circadian rhythms (*e.g*. Andreeva et al. 2021).

## Acknowledgments

The authors acknowledge the European Synchrotron Radiation Facility (ESRF, Grenoble) for provision of beam time on CM01. The IBS/ISBG electron microscope facility is supported by the Auvergne-Rhône-Alpes Region, the Fondation Recherche Médicale (FRM), the fonds FEDER and the GIS-Infrastructures en Biologie Santé et Agronomie (IBISA). We acknowledge the platforms of the Grenoble Instruct-ERIC centre (ISBG; UMS 3518 CNRS-CEA-UGA-EMBL) within the Grenoble Partnership for Structural Biology (PSB). Platform access was supported by FRISBI (ANR-10-INBS-0005) and GRAL, a project of the University Grenoble Alpes graduate school (Ecoles Universitaires de Recherche) CBH-EUR-GS (ANR-17-EURE-0003). IBS acknowledges integration into the Interdisciplinary Research Institute of Grenoble (IRIG, CEA). The project received further support by institutional grants to the Laboratoire de Physiologie Cellulaire et Végétale by Labex Grenoble Alliance of Integrated Structural Biology (GRAL) and ANR-17-EURE-0003.

The authors thank M. Webster for early sharing of atomic coordinates of the SaPEP; R. Ruedas for help with PRIN2 crystallization; M-C. Ducarre for help with cloning. We thank C. Carles for critical reading.

## Author contributions

FXG, RB and DC designed the research. RB and DC collected fundings. FXG, GE, GLFvS, MT, NP, DF, AV, SB, YC, RB, and DC performed the research. FXG constructed the genetic tools for AP/PL, collected and analyzed data. MT and NP cloned and tested PAP8 partners. SB and YC contributed MS-based proteomic data. DF, GLFvS and GE contributed EM data. RB and DC wrote the manuscript with contributions from FXG, YC. All authors approved the manuscript.

## Limitation of the structural studies

Some regions of the PEP structure were not resolved in the cryo-EM electron density map (Table S1). The overall chain of β″ and PAP11 could not be built due to limitations in the quality of the cryo-EM electron density map. The polyalanine model was employed for the N-terminal part of PAP2, highlighting the flexibility of this subunit. Additionally, both N-terminal and C-terminal segments of several PAPs were not observed (Table S1). It is possible that some of these missing segments could become visible upon binding to DNA or protein partners. The structure of PEP described here is an open conformation, which may undergo changes during transcription stages coupled with translation and throughout plant development, given its isolation from very young cotyledons. Further genetic, biochemical and structural investigations are essential to fully characterize the proposed model for transcription initiation, ribosome binding, and the roles of the PAPs.

## Supplemental Figures

**Figure S1:**
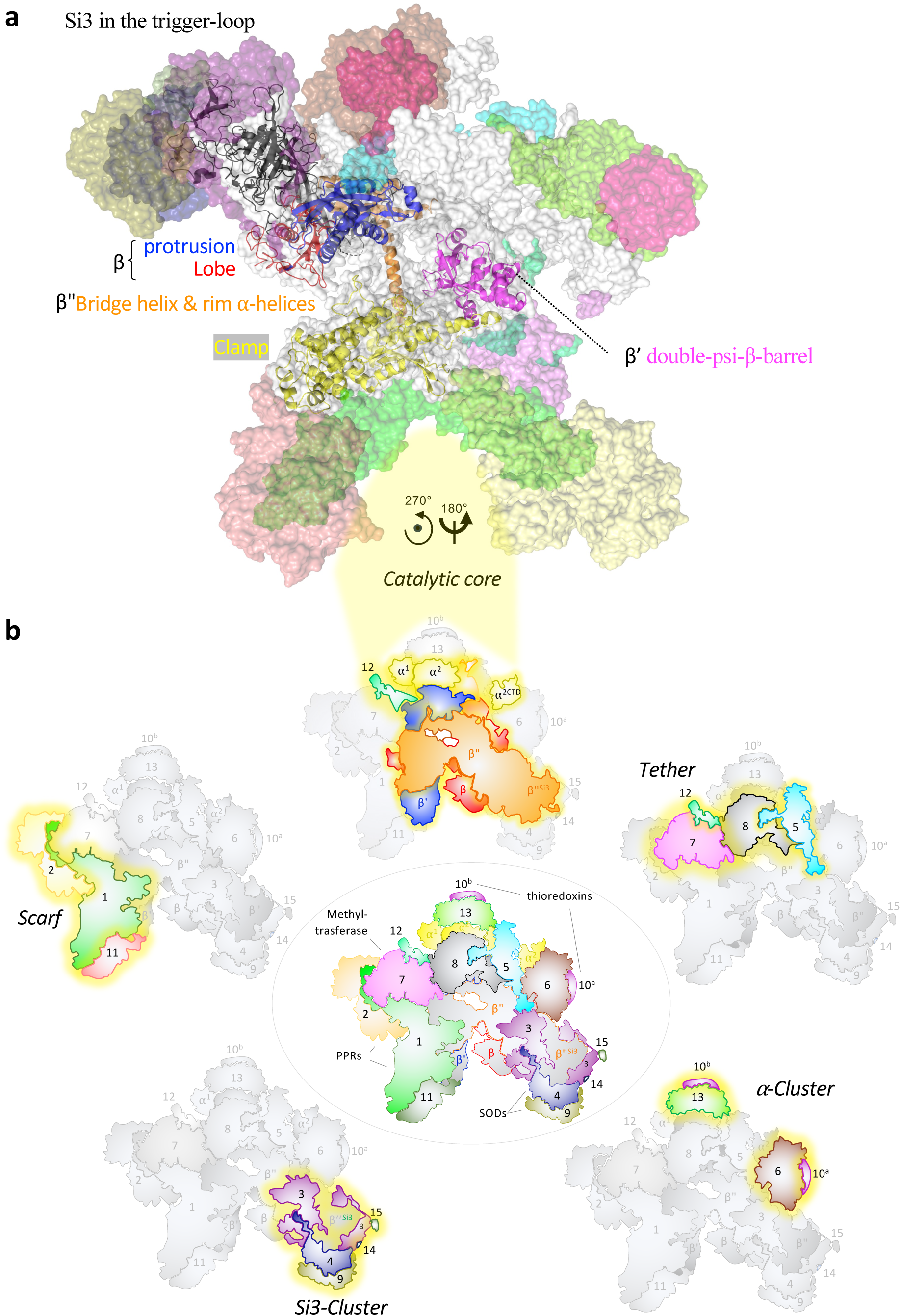
The catalytic core and the PAP clusters. **(a)** View of the lobe (red) and protrusion (blue) of β, the double-psi-β-barrel domain (magenta) of β’, the bridge helix (orange) and rim α-helices (orange) of β’’, and the clamp (yellow). Si3 in the trigger-loop is colored in black. The surface of catalytic subunits is white colored and the PAPs color-coded as in Figure 1. **(b)** Topological representations of the PEP subunits highlighting their relative positions within the complex and organized in four clusters and the core; each number correspond to the serial PAP color-coded as in Figure 1. Note that PAP12 conceptually belongs to the core as the ω homolog and the tether cluster as a PAP7 interactor that is dually localized. Center view: some enzymatic activities and domains of PAPs are given. Surrounding views: subunits are separated in four different clusters around the catalytic core.

**Figure S2:**
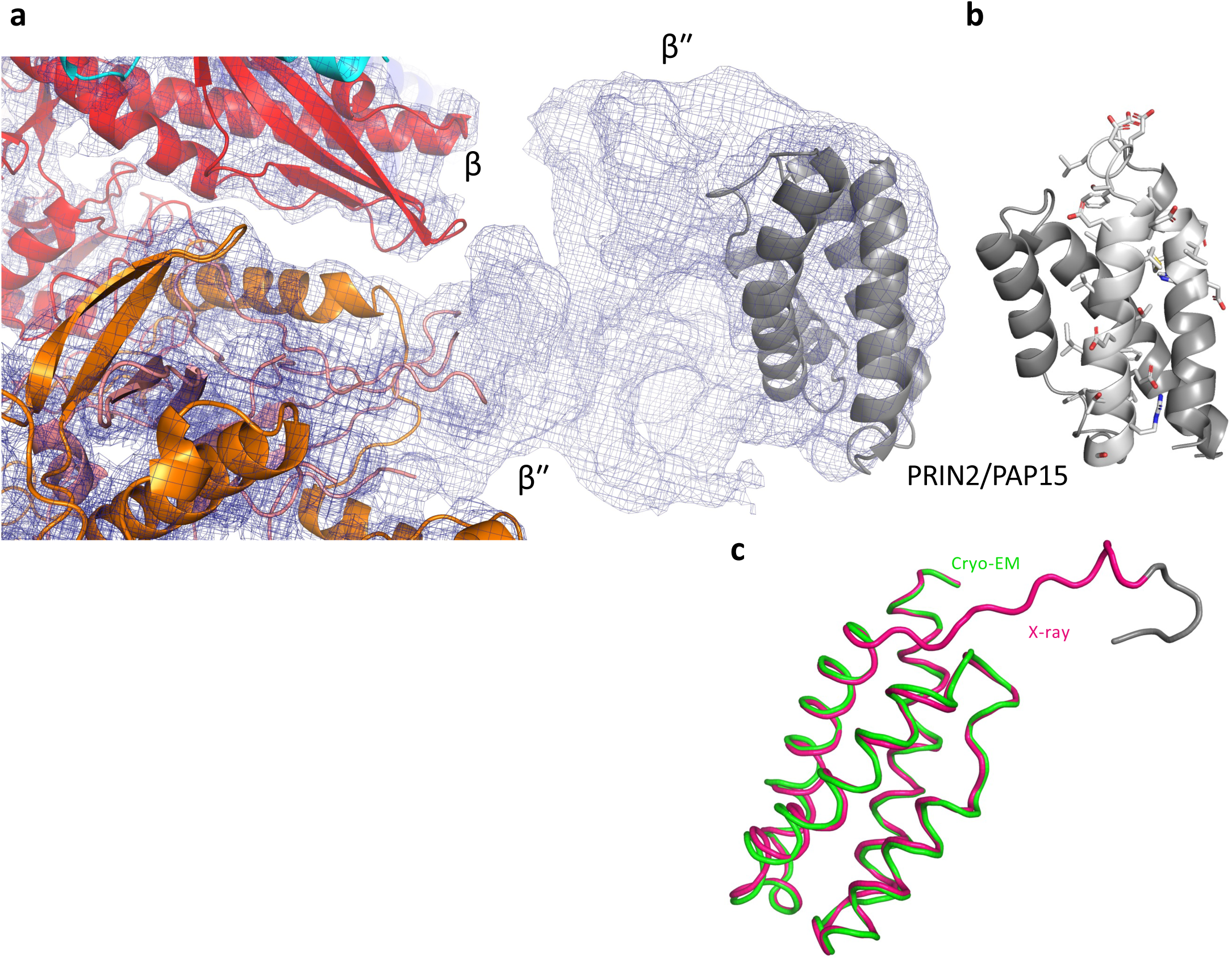
**(a)** View of the missing model of β’’ in the cryo-EM electron density map. β is red colored, β’’ is colored in orange, and PAP5 in cyan. β’’ interacts with PRIN2 (PAP15), a redox protein that binds non-specifically to DNA (Diaz et al., 2018). PAP15 displays three predicted nine amino-acid transactivation domains fully accessible in the PEP. Its crystal structure was solved at 1.72 Å resolution (Table S3). PAP15 folds as a bundle of four α-helices and was found in MS/MS analyses of the SaPEP (Ruedas et al., 2022). PRIN2 is displayed in gray in the cryo-electron density map. **(b)** The structure of PRIN2 in the PEP, after rotation, is displayed with the amino-acid side chains of the TADs shown in sticks. **(c)** View of the superimposition of PRIN2 as in the PEP (green) onto PRIN2 solved using X-ray (pink); they superimpose with an rmsd calculated on C⍺ atoms of 0.666 Å. The longest N-terminal part of the mature PRIN2 solved using X-ray diffraction contains also four residues of the thrombin cleavage site and His(-1)-Met(0) colored in gray.

**Figure S3:**
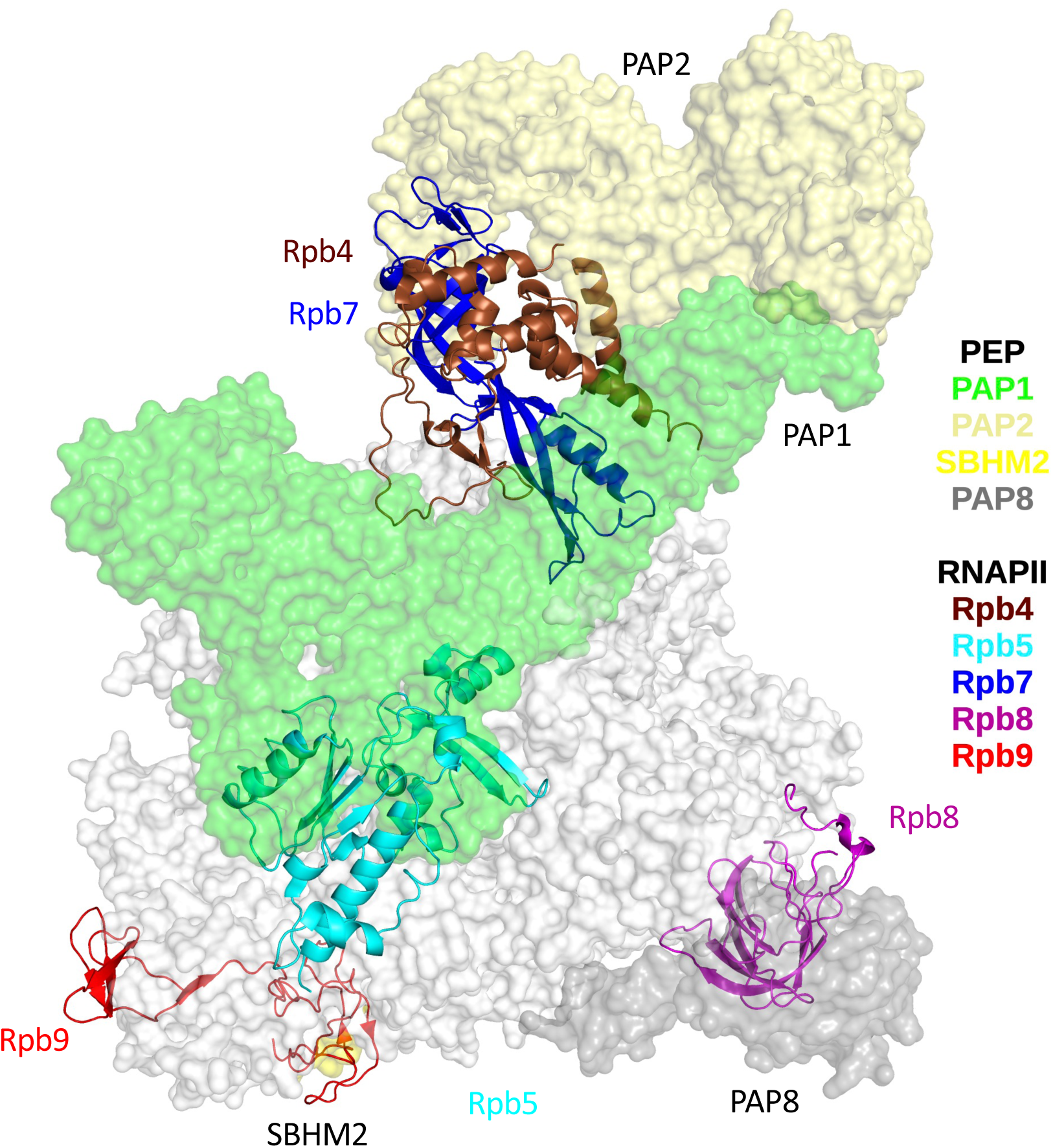
Surface representation of SaPEP superimposed on Rpb4 (maroon), Rpb5 (yellow), Rpb7 (blue), Rpb8 (magenta) and Rpb9 (red) from RNAPII. The surface of PAP1 is in green, PAP2 in lemon, PAP8 in gray, and SBHM2 in yellow. As in SeRNAP, the fin of Si3 locates to a similar position than Rpb9, which maintains the transcription fidelity in RNAPII (Nesser et al., 2006).

**Figure S4:**
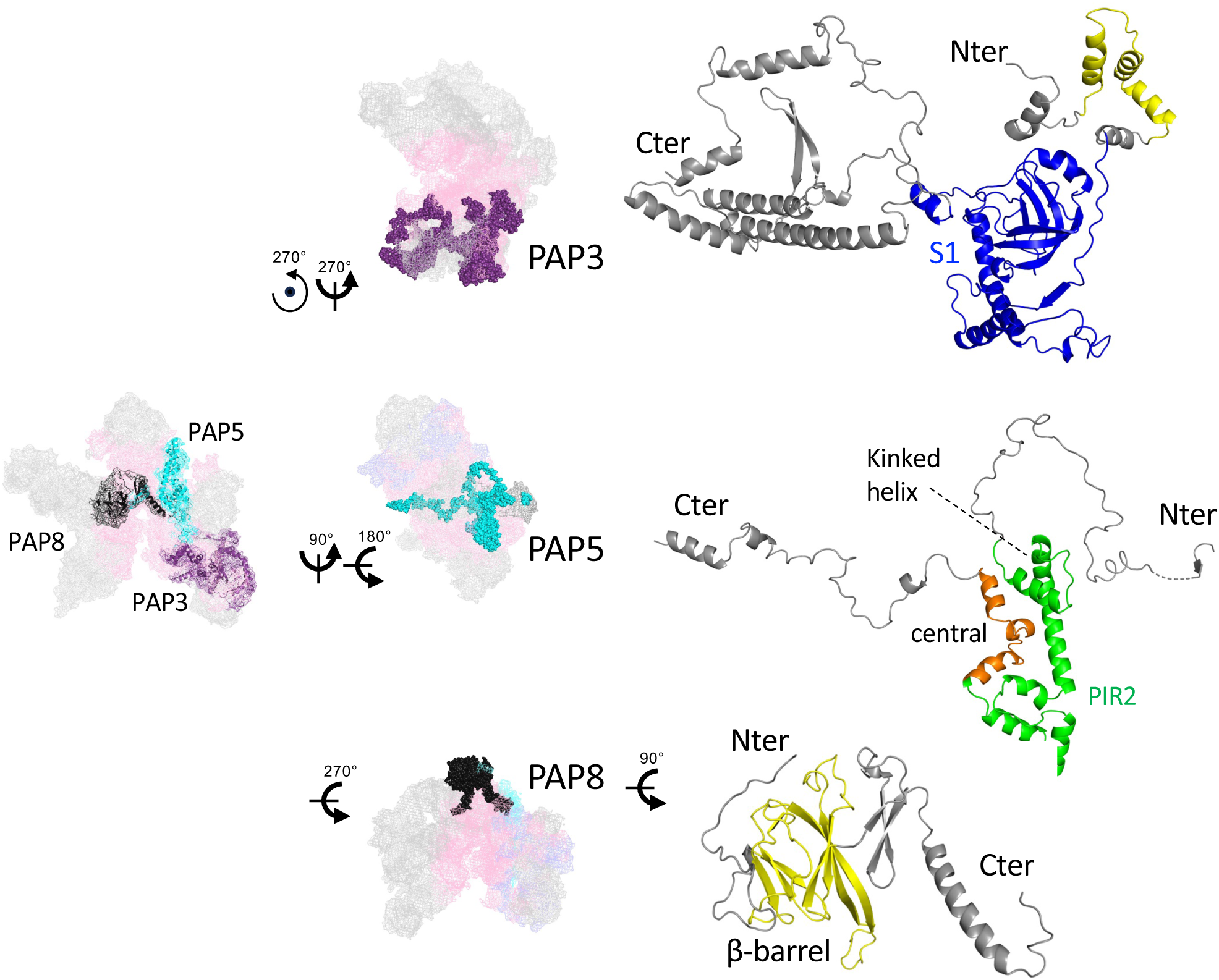
Structures of PAP3, PAP5 and PAP8, alone or in PEP context. The catalytic core is displayed in light pink mesh, other PAPs in gray. Color code of PAP3, 5 and 8 as in Figure 1. PAP3 has three domains: the N-terminal domain where L88 to I133 shares structural similarities with mitochondrial pyruvate carboxylase; a globular central domain (D148 to G379) harbors a domain S1 akin to those in the exosome complex, 30S ribosomal protein S1, or RNAPII’s Rpb7; and the C-terminal domain (R380 to R614) lacks structural homology with known proteins. The PAP3 globular domain with its domain S1 is blue colored. The N-terminal of PAP3 interacts with the tail and fin of Si3; the two first α-helices wrapping SMBH2 as a horseshoe. The central domain interacts with SMBH3, SMBH8, the domain S1 seating between SMBH8 and the extra domain of Si3. The C-terminal part forms an antiparallel six-stranded β-sheet with SMBH4. PAP5 has an unfolded N-terminal part (N184-D254), a C-terminal part (G385-T441) with one α-helix (R429-T441), and a central domain (V255-G384) that contains seven α-helices, one of them (L278-E306) kinked in the front of the rim α-helices of β’’. This domain does not share similarity when compared with the structures of the PDB. The PAP5 central domain is colored in orange with PIR2 colored in green. The PAP8 β-barrel domain is yellow.

**Figure S5:**
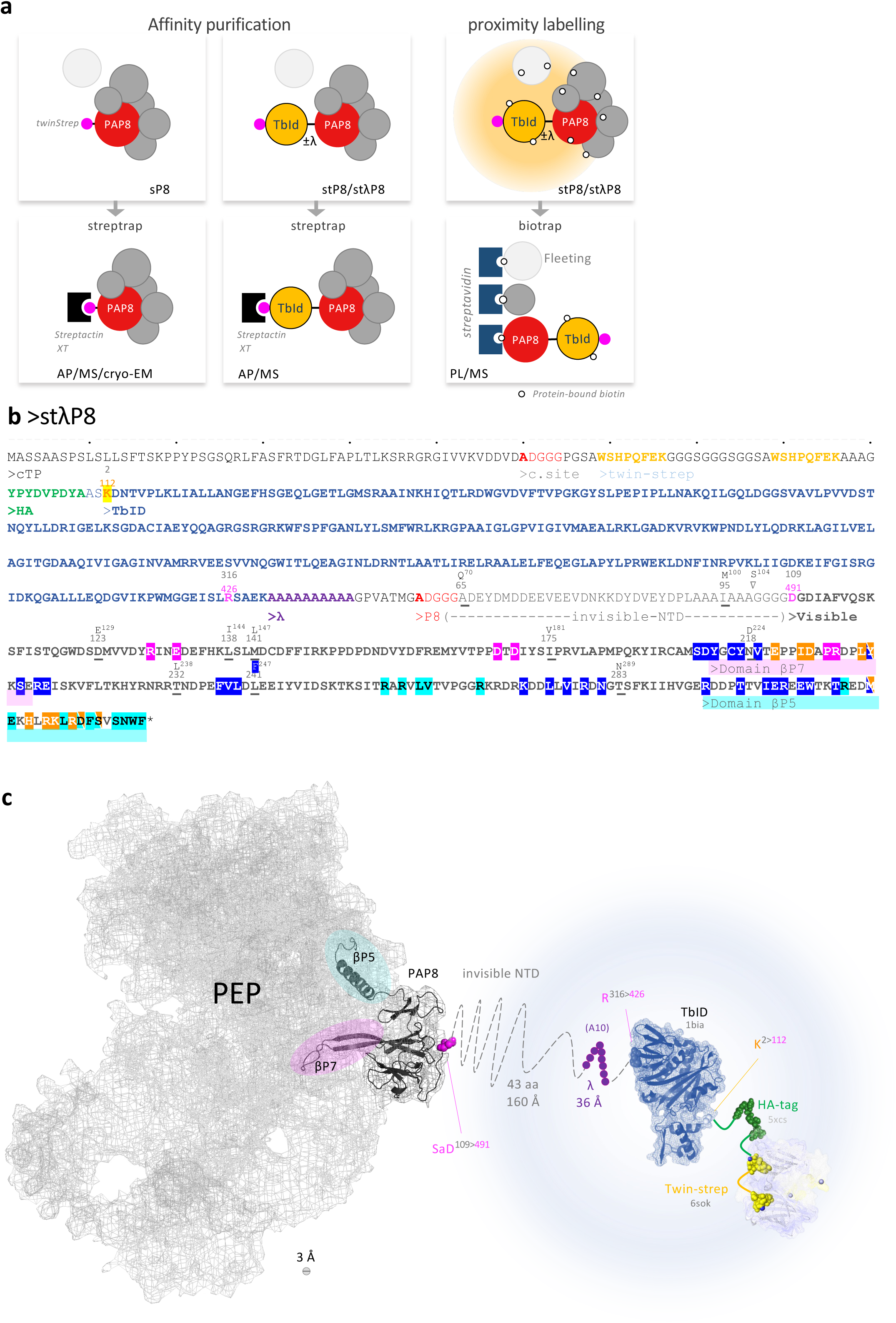
Experimental setup using PAP8 translational fusions. **(a)** twinstrep in magenta; TbID, hemagglutinin fused to TurboID; λ, poly-ALA (A_10_) used as linker; PAP8, processed PAP8 DNA fragments were assembled into sP8, stP8 and stλP8 as in figure 4. sP8 in homozygous *pap8-1* mutant was used for affinity purification (AP) followed with Mass Spectrometry (MS) and negative stained electron microscopy. stP8 and stλP8 in *pap8-1* were used both in AP and PL respectively; both followed with MS. **(b)** Amino acid sequence of stλP8 (cloned from *Arabidopsis thaliana*); cTP, chloroplast transit peptide; c. site, cleavage site; twin strep tag; HA, hemagglutinin tag; TbID, TurboID; P8, PAP8 CDS; βP5 and βP7 domains correspond respectively to those interacting with β′, β″, PAP5 or β′, β″, PAP7; highlighted amino acids in the color code corresponds to the contact points with the corresponding proteins. Underlined residues are those different from SaPAP8 given on top with its position in the protein (gray numbers) or in the fusion (magenta numbers). Note that differences in the cTP are not given. **(c)** Cartoon representation of stλP8 within the PEP complex; the dashed line is the 43 invisible amino acids of PAP8 corresponding to a polypeptide chain of 160 Å. TbID HA and twin strep were taken from PDB accessions 1BIA, 5XCS, and 6SOK, respectively.

**Figure S6:**
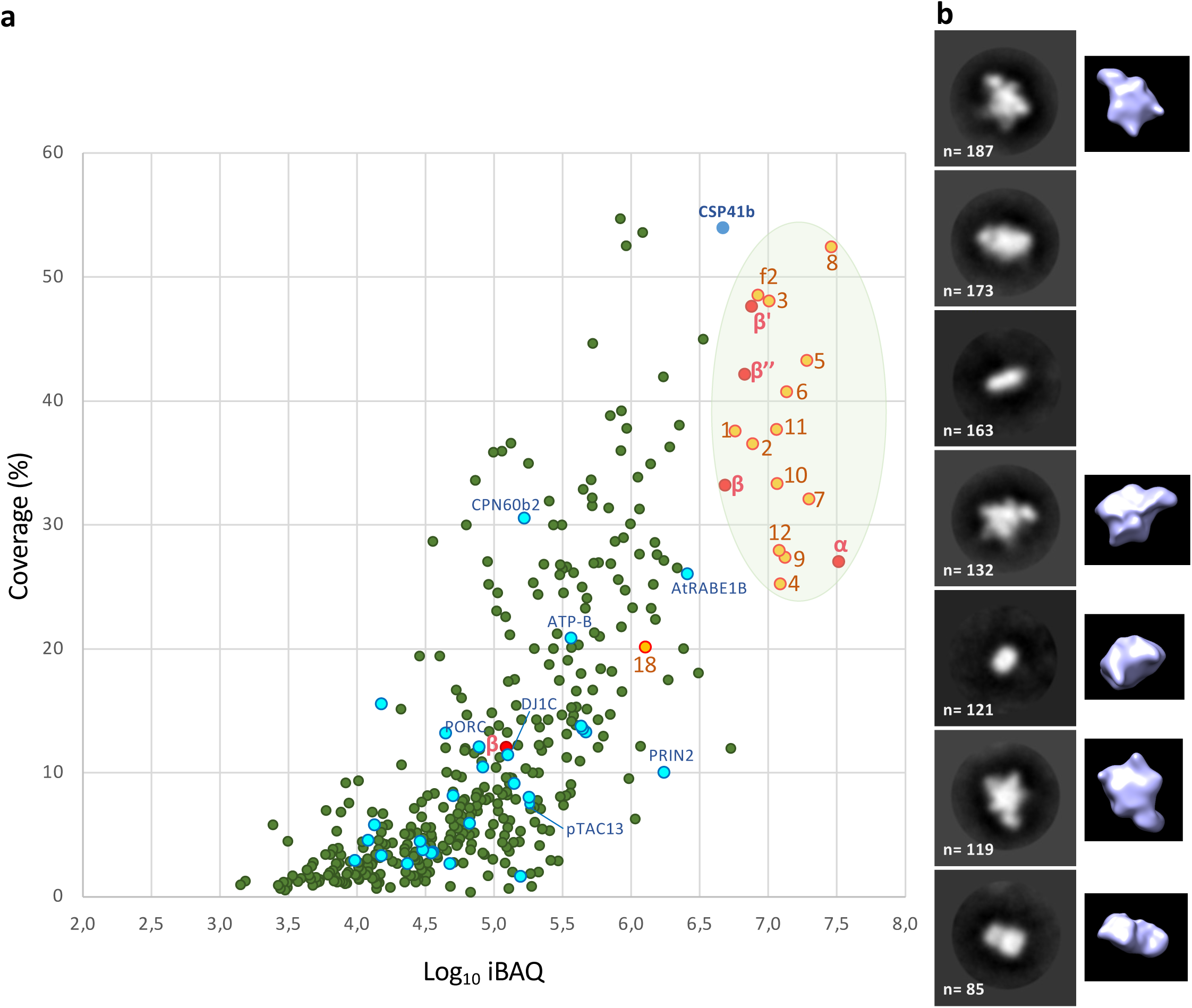
AtPEP proteomic and structure studies (dataset 1). (**a**) Scatter plots of the Mass Spectrometry-based quantitative proteomics analysis presented as Log_10_iBAQ=f(coverage). Components plotted in red: core subunits α, β, β’ and β’’; orange: PAPs (1-12) as their corresponding numbers, pTac18 as 18; FLN2 as f2; blue: proteins of the common set (Figure 5) including PRIN2; except CSP41b found in only a subset of the skoto-to photomorphogenesis analysis. All the expected components of the PEP fall in the shaded yellow area. They correspond to the major protein mass contribution to the purified sample. (**b**) Classification of 980 particles from negative-stained electron microscopy of AtPEP purified from sP8. The AtPEP envelope was calculated at 31-Å resolution.

**Figure S7:**
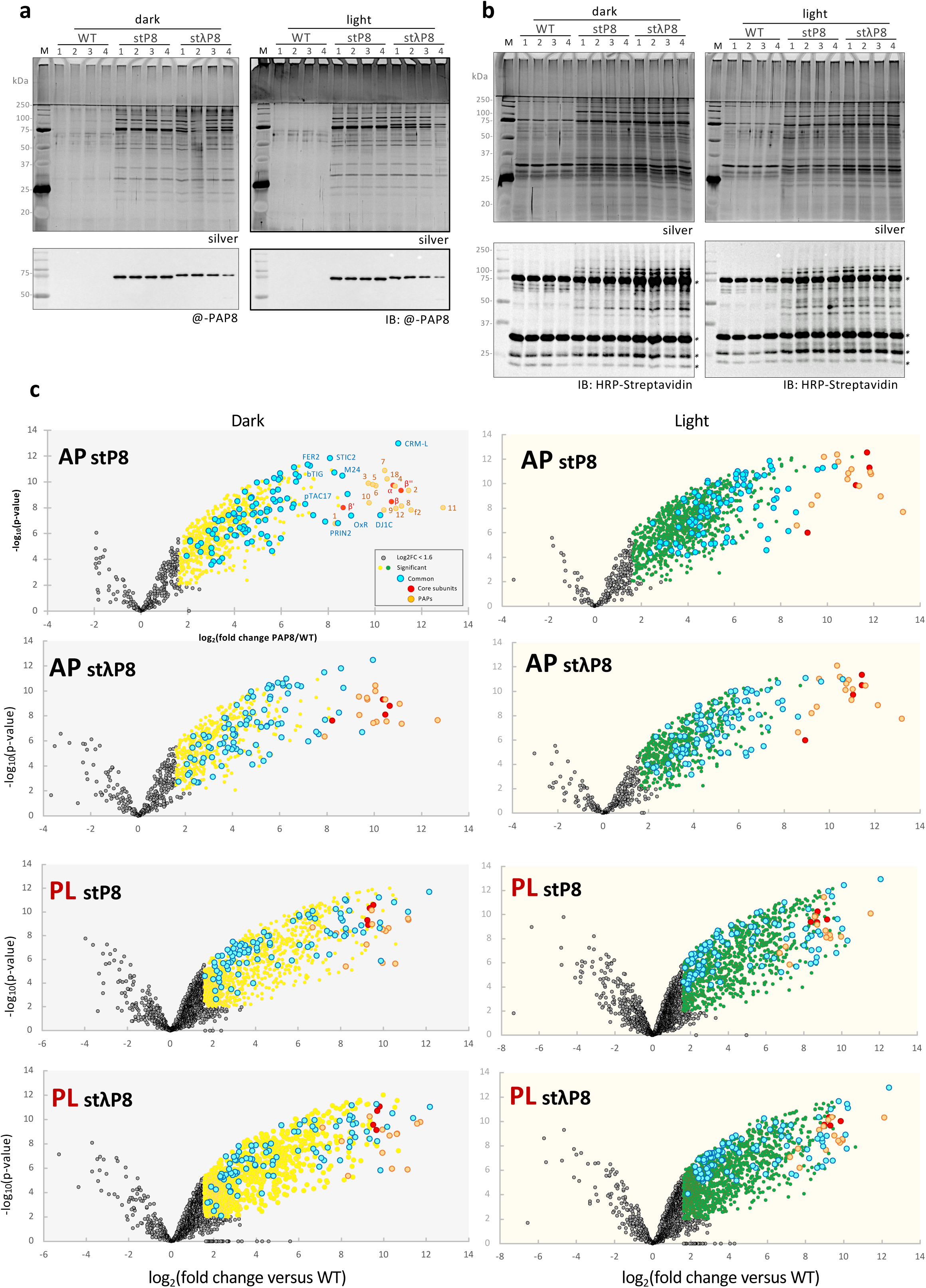
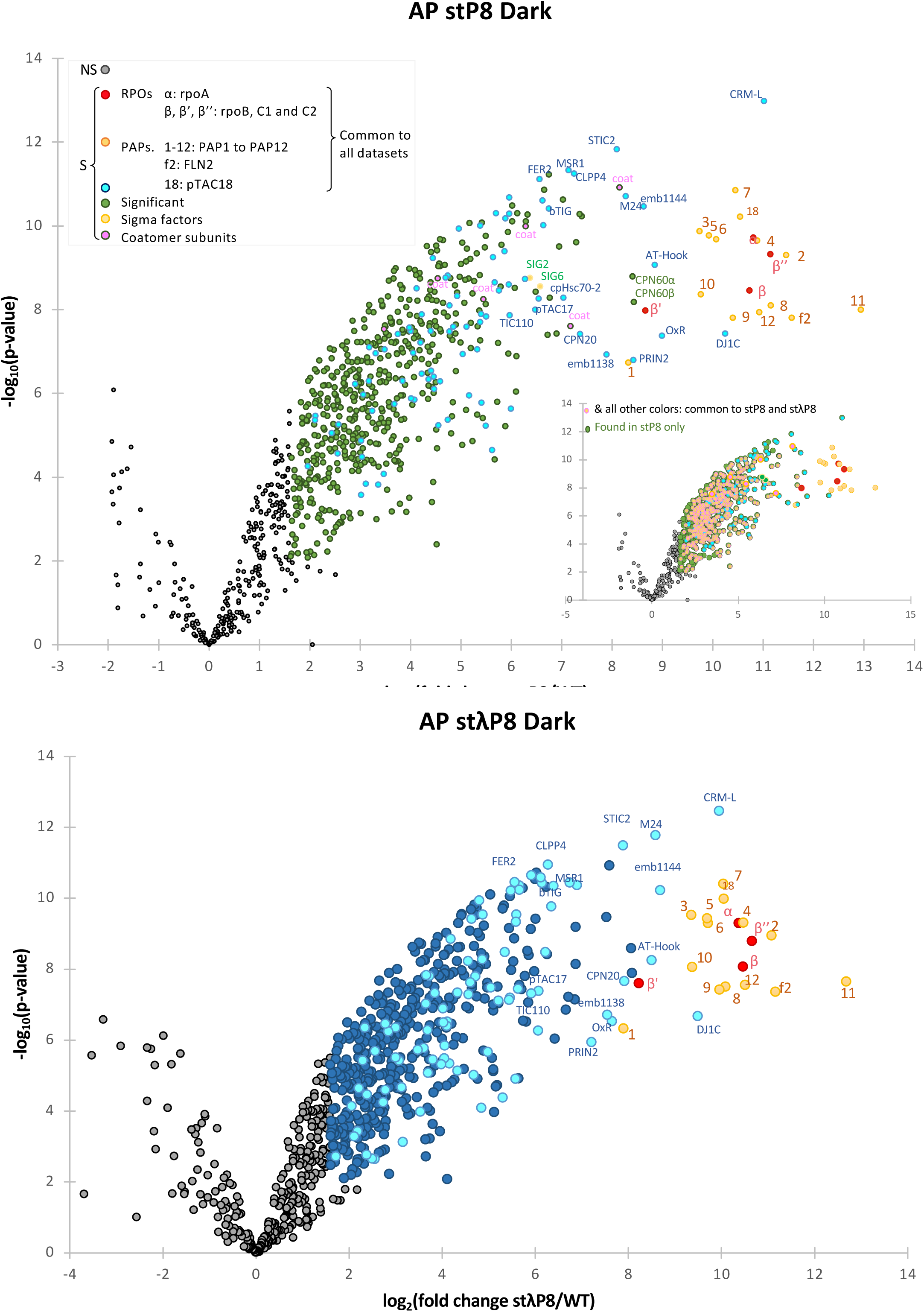
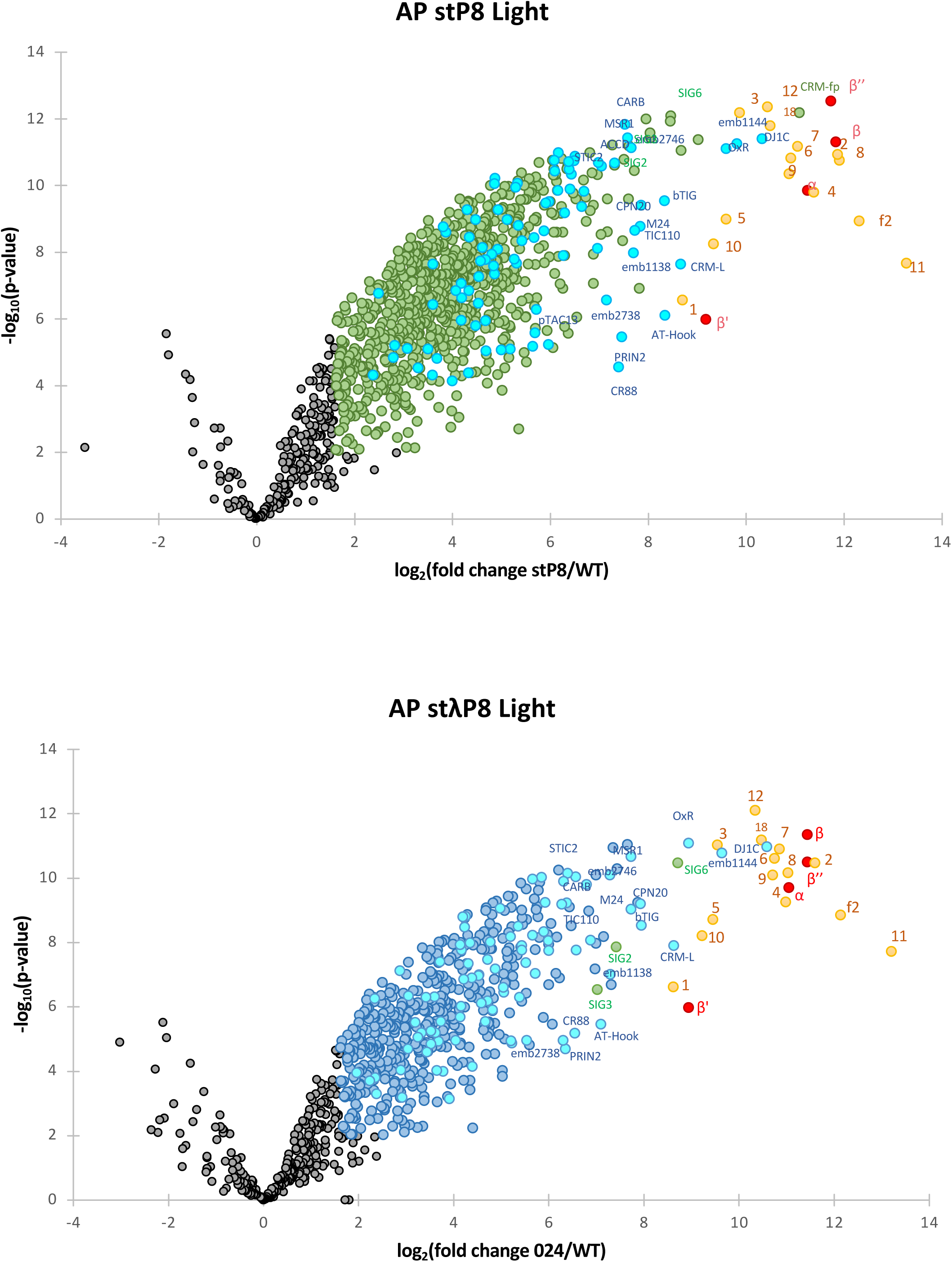
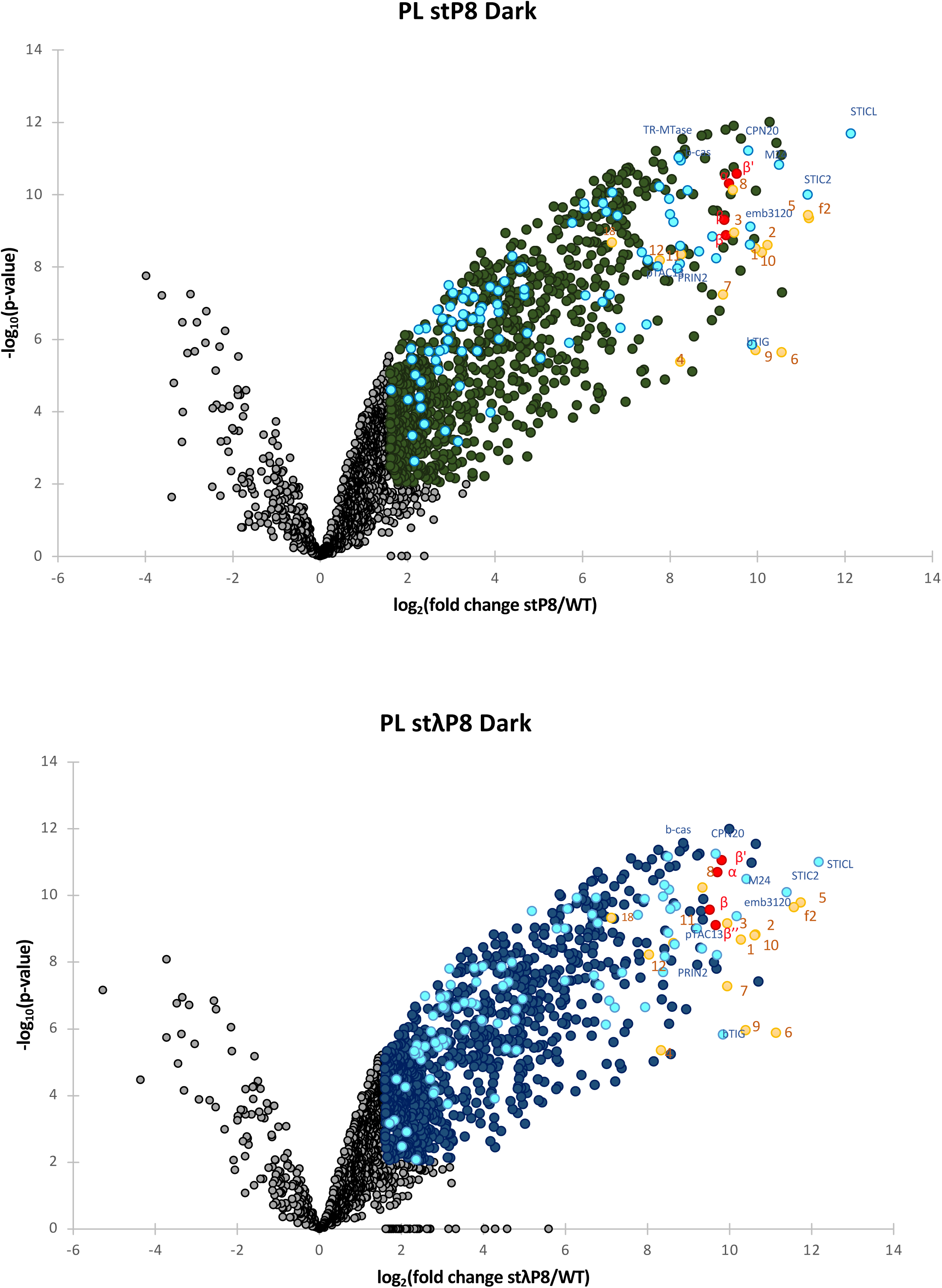
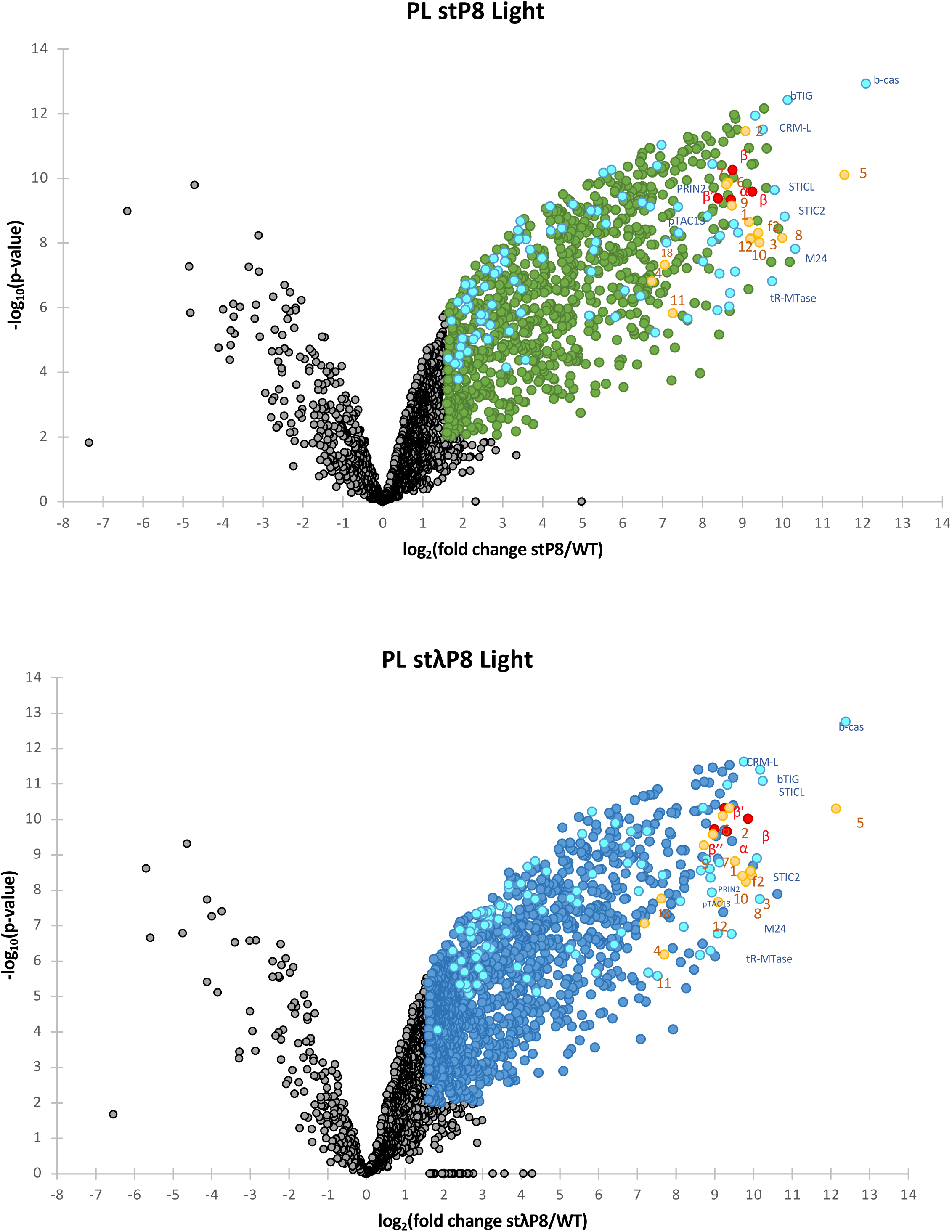
Sample preparation for MS and their respective volcano plots. **(a, b)** Quadruplicates of WT, stP8, and stλP8 corresponding to 500 µL of dry seeds each were grown on separate plates in the dark or dark plus 8 h of light. Seedlings were harvested and protein samples were prepared for AP (a) or PL (b). Silver-stained membranes (Silver) IB immunoblots (anti-PAP8 or Streptavidin-HRP). **(c)** Volcano plots of the corresponding proteomes. Significant protein numbers are given in parenthesis. Yellow and green dots (with the other colors described below) for dark or light conditions respectively, represent differentially detected proteins using a log_2_(Fold Change) cut-offs of 1.6 for WT versus stP8 or stλP8, and a p-value cut-off of 0.01. Red dots, PEP core subunits (⍺, β, β′, β″); orange dots, PAPs (PAP1 to PAP12 (1-12), FLN2 (f2) and pTAC18 (18) and 89 blue dots represent collectively the 107 proteins found in all datasets (common proteins).

**Figure S8:**
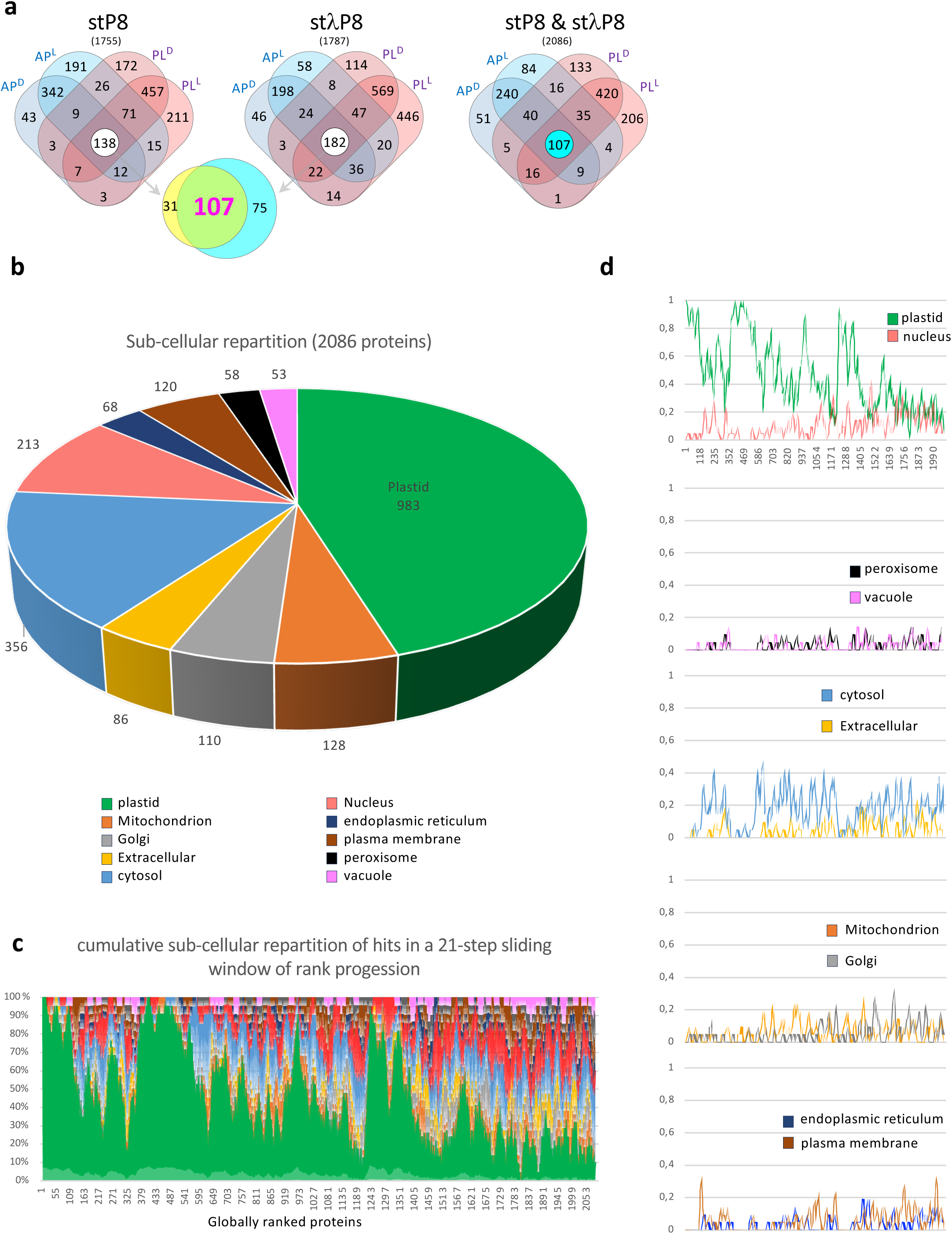
**(a)** Venn diagrams and **(b)** localization prediction using SUBA with **(c)** cumulative sub-cellular repartition of hits in a 21-step sliding window in the global rank progression. **(d)** Curve in **(c)** breaks down for 2 compartments at a time.

**Figure S9:**
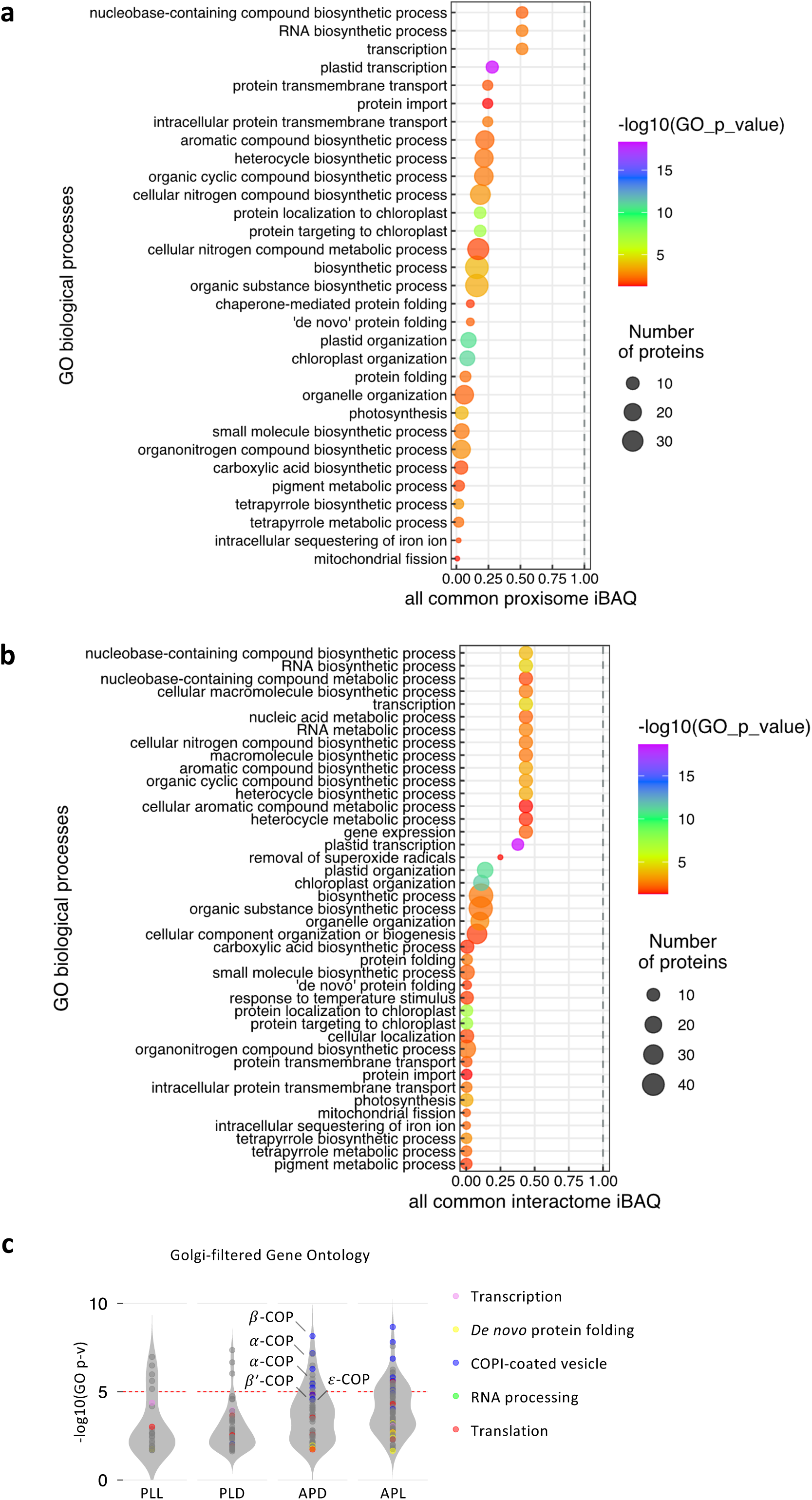
Gene Ontology (GO) analysis presented as lists of enriched GO in PL **(a)** or AP **(b)** extracted from the 107 common proteins with Panther (https://pantherdb.org) and Revigo (Supek et al., 2011) according to the protocol in Bonnot et al., (2019). **(c)** GO cell compartment (GOCC) annotation extracted from TAIR (https://v2-arabidopsis-org) of all significant accessions (2086) filtered for Golgi (110 acc.), sorted by cellular functions color-coded as indicated, and represented as a violin plot.

**Figure S10:**
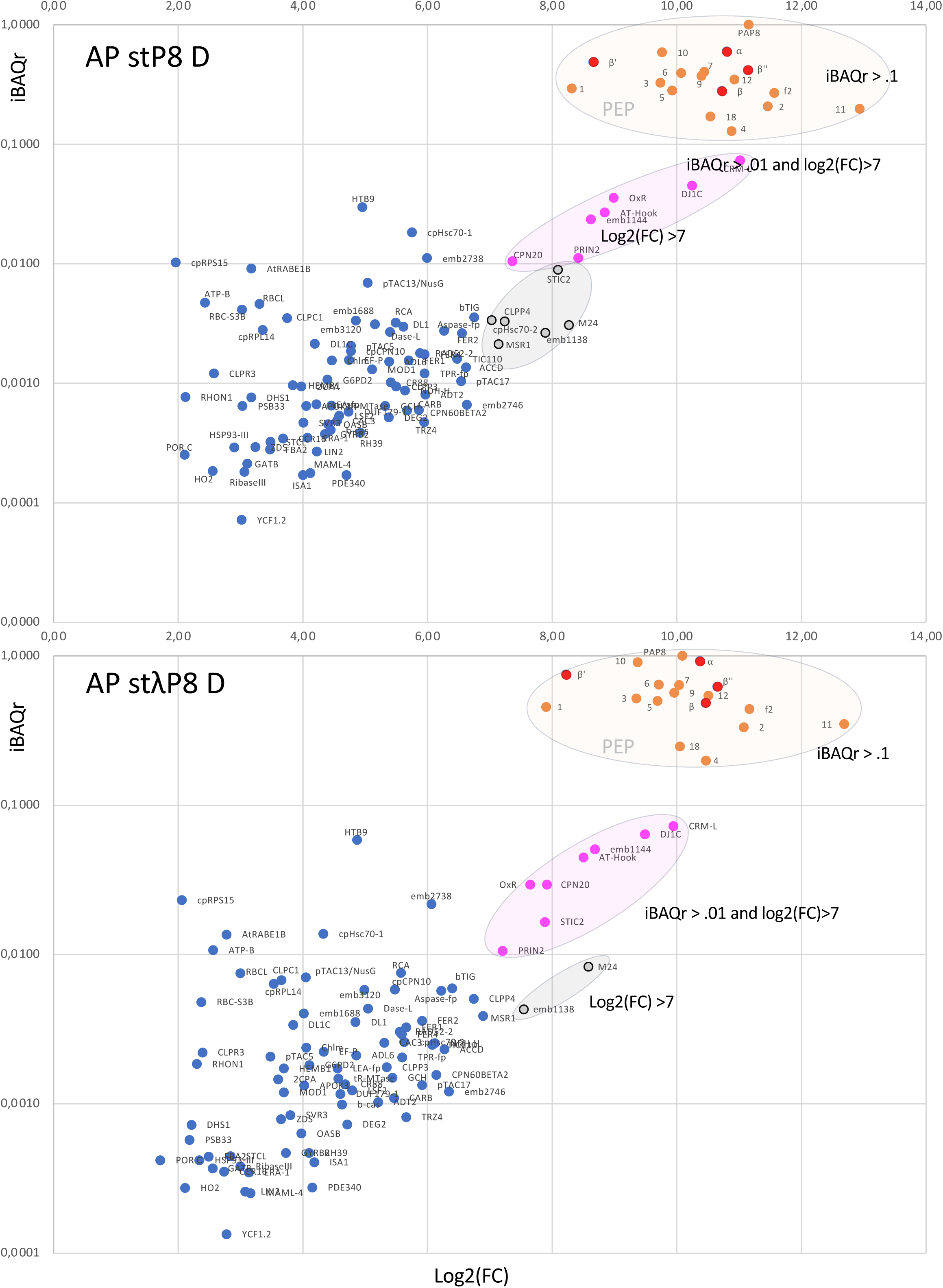
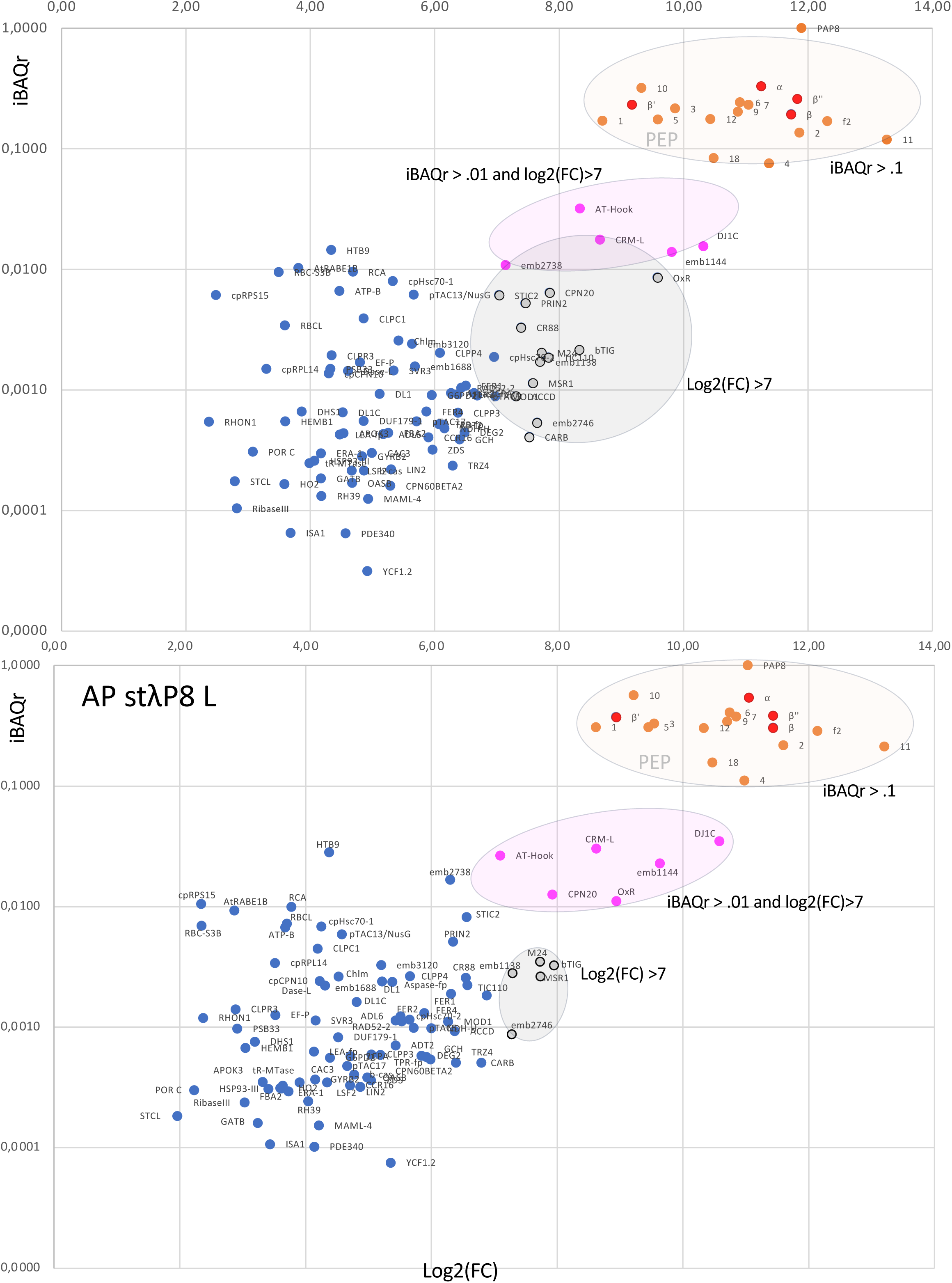
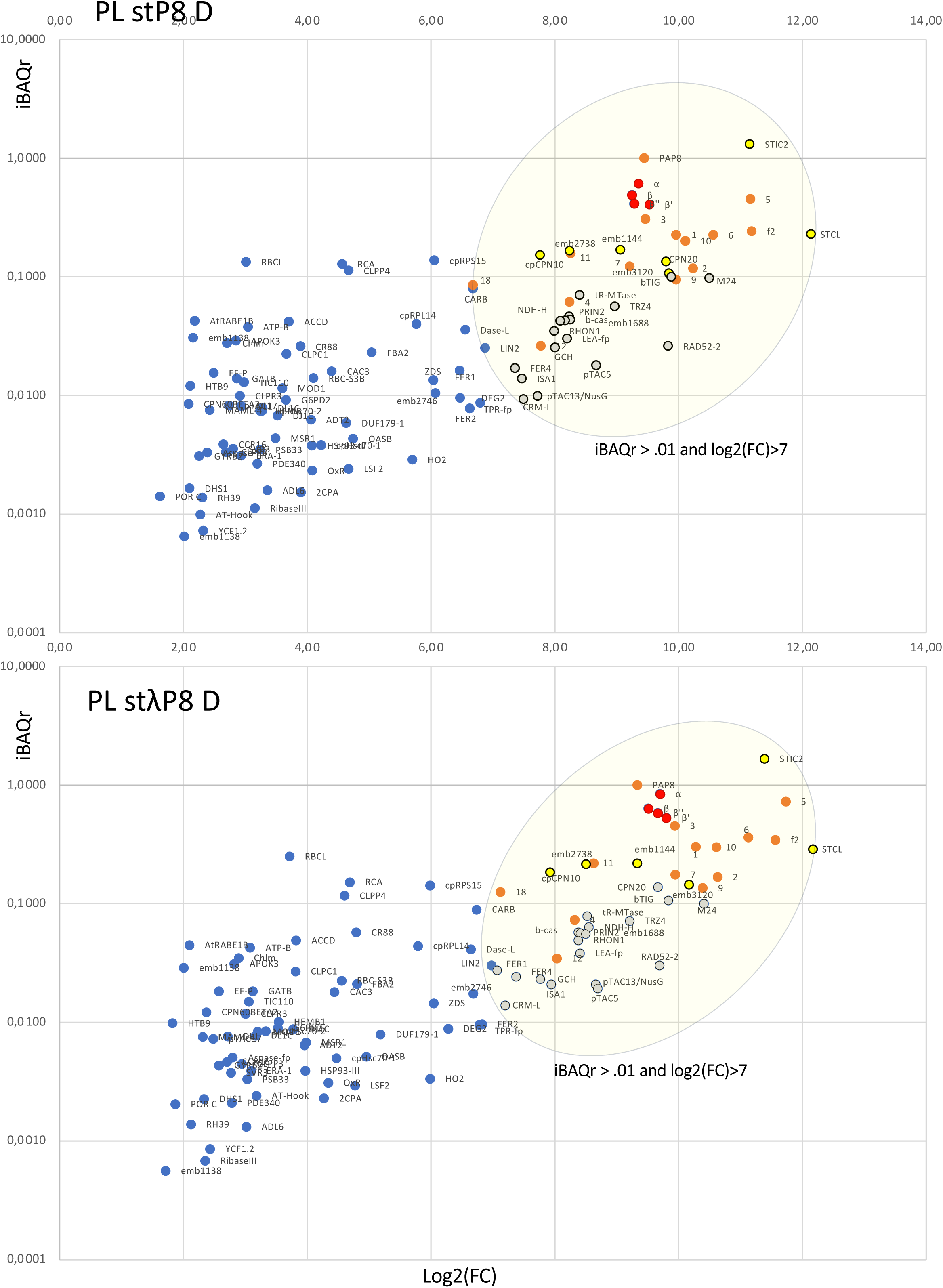
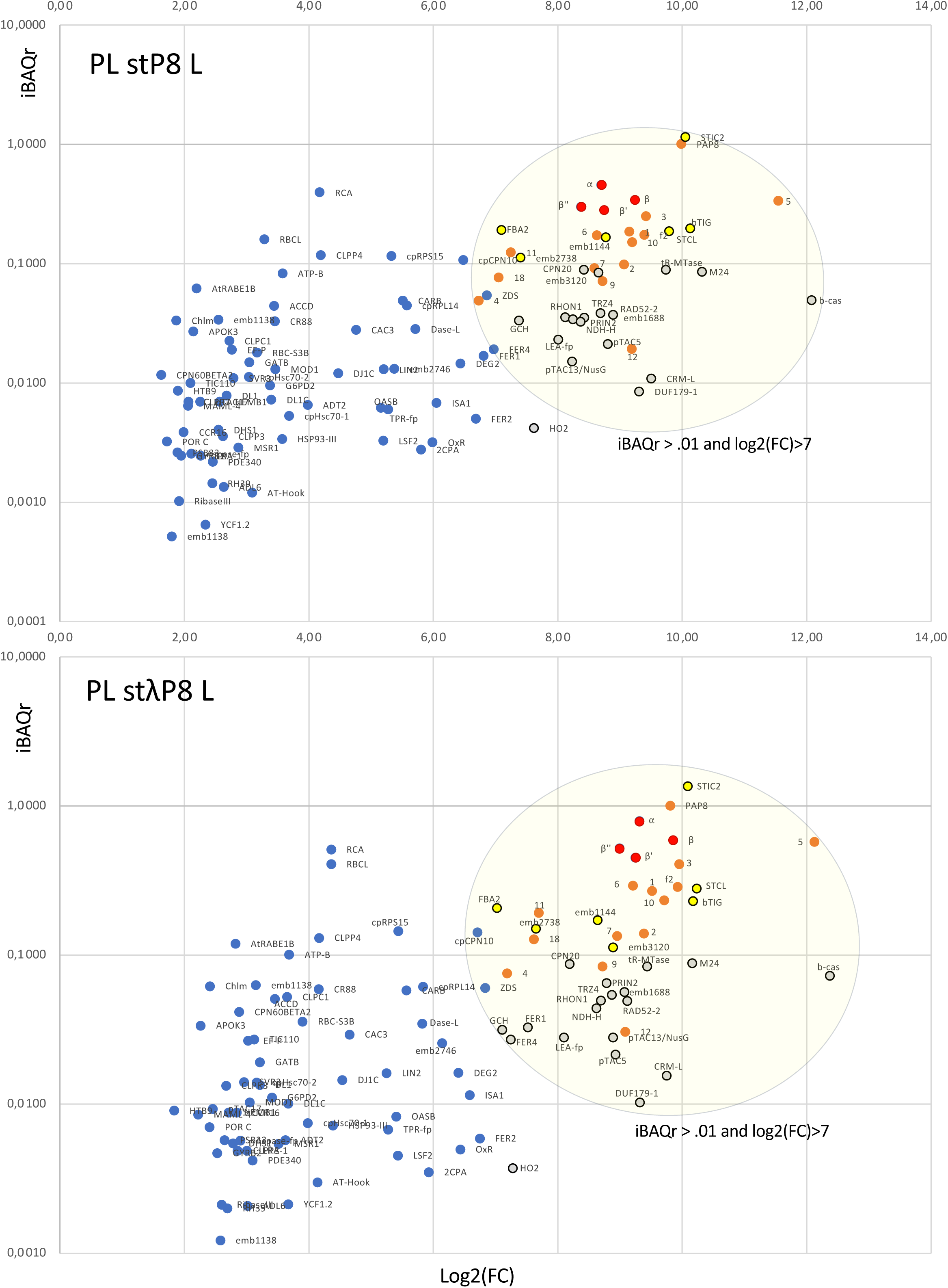
Stoichiometry analysis of common proteins for the eight datasets (AP/PL)x(stP8/stλP8)x(D/L)

**Figure S11:**
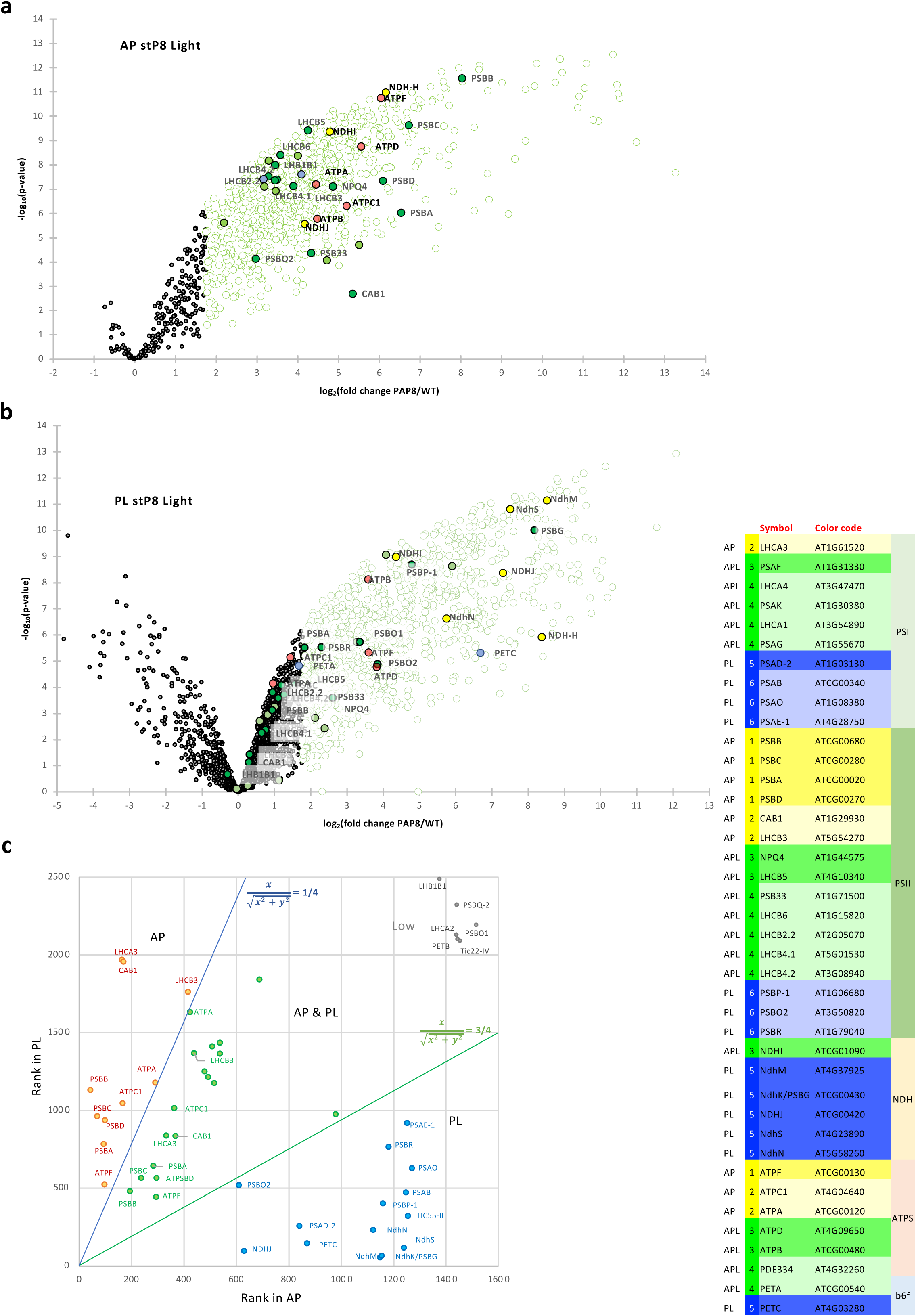
Light versus dark analysis. **(a, b)** Volcano plots of AP stP8 Light **(a)** and PLstP8 Light **(b)**. **(c)** Candidate proteins plotted according to their rank in AP (y-axis) versus PL (x-axis). 4 area have been set for classification.

**Figure S12:**
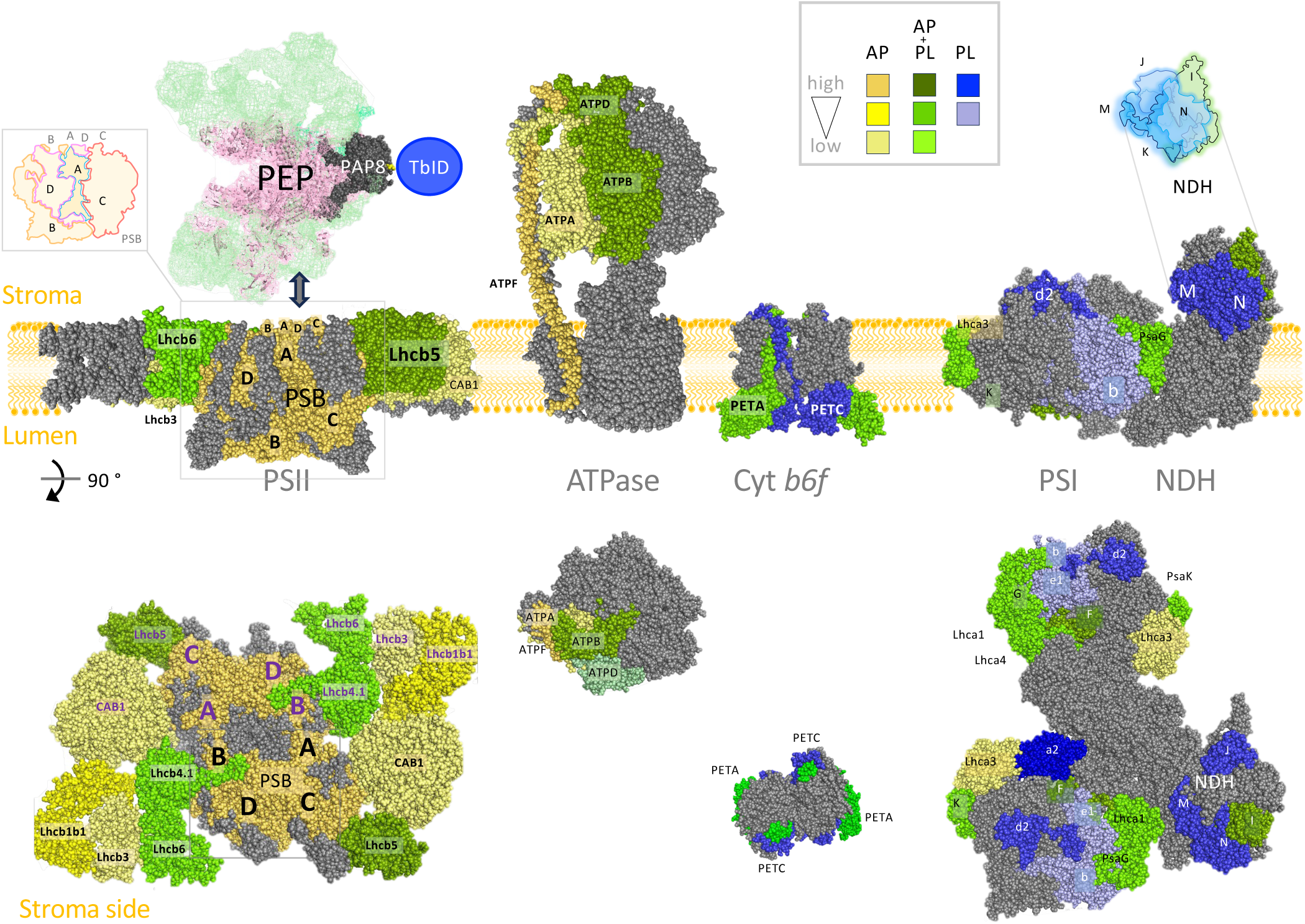
Front view from the stroma side of the photosynthetic complexes shown in Figure 6.

**Figure S13:**
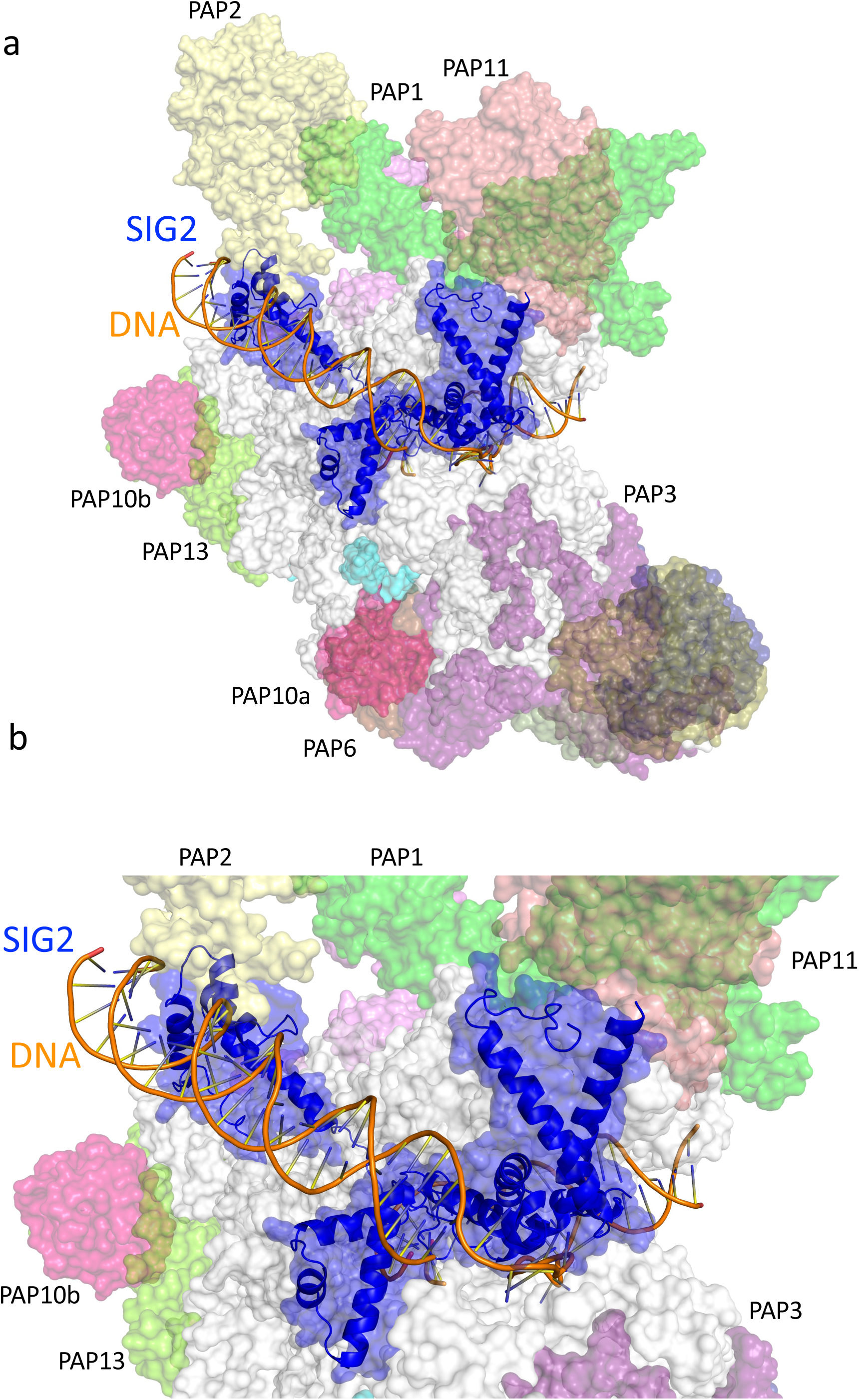
Modeling of the initiation complex SIG2/DNA with SaPEP. The PEP overall surface is displayed in transparency. PAP2 (lemon), PAP11 (pale magenta), SIG2 (blue) and DNA are drawn in cartoon. (**a**) Overall view. (**b**) Zoom on the interactions involving PAP2 and PAP11. The SaPEP catalytic core is white colored. The PAPs are color-coded as in Figure 1.

**Figure S14:**
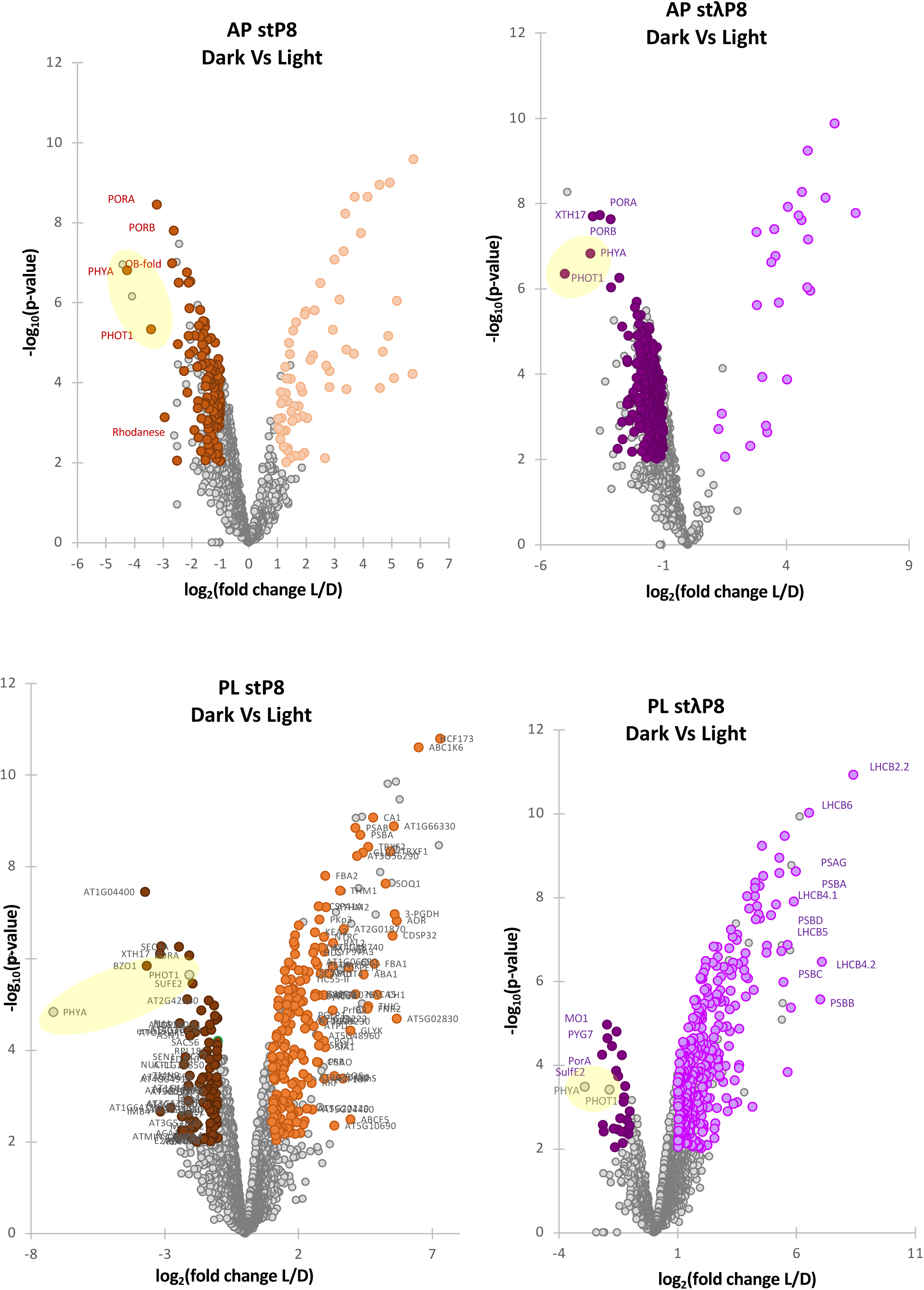
Volcano plots from the Dark-enriched versus light-enriched proteins in the different datasets (AP/PL)x (stP8/stλP8). The yellow areas highlight the photoreceptors.

**Figure S15:**
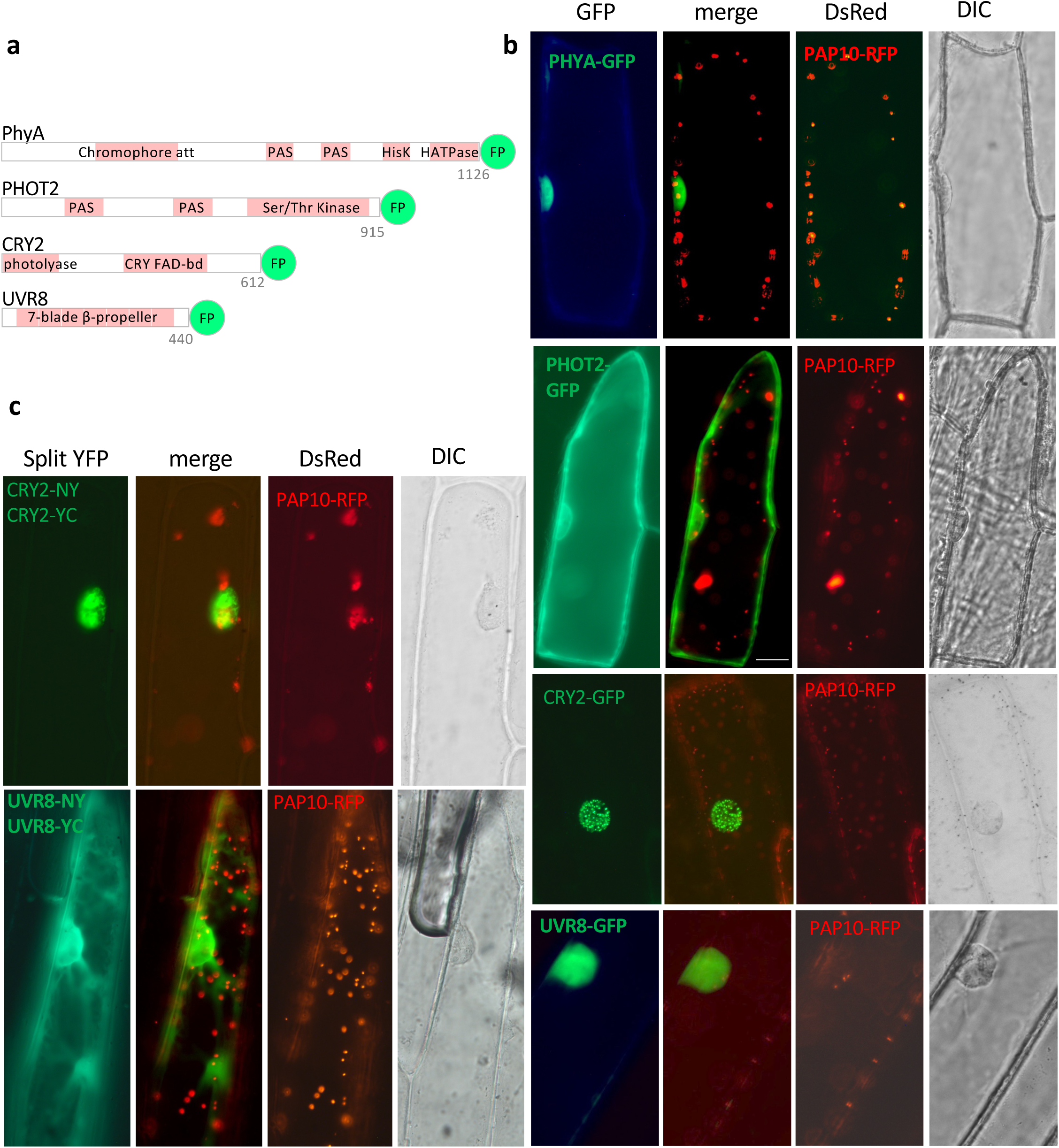
Photoreceptor-GFP translational fusions tested in transient assay. **(a)** Fusion proteins presented in domains as annotated in the TAIR (light pink); numbers in gray gives the length of the photoreceptor in amino acids. **(b)** GFP fusions tested in onion cells. **(c)** BiFC test of CRY2 and UVR8 as indicated. **(b, c)** The DIC image gives the position of the nucleus (ca. 20 µm in diameter); PAP10-RFP marking the plastids used as an internal control.

**Figure S16:**
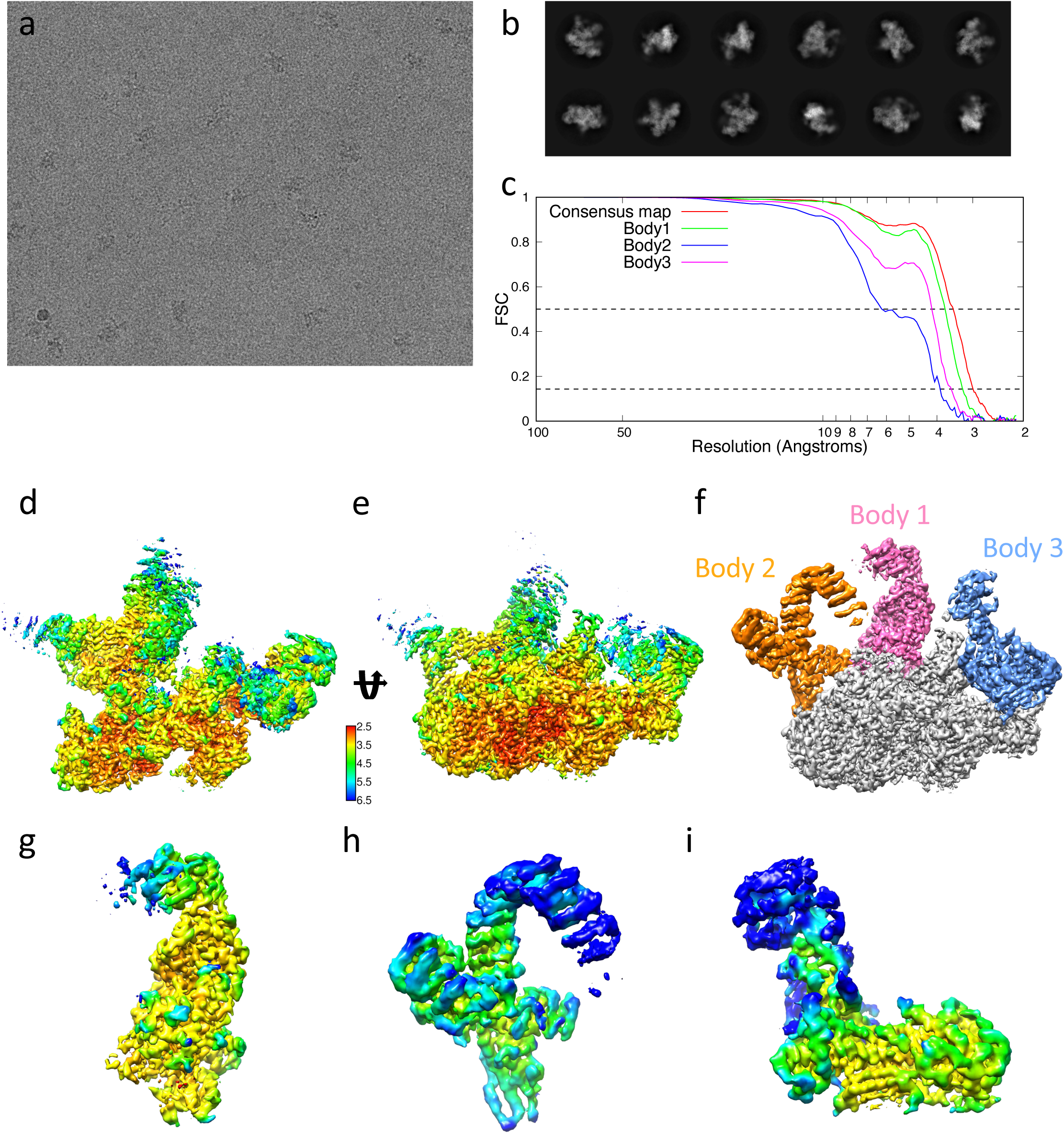
**(a)** Cryo-EM field of view of SaPEP. **(b)** Representative 2D class averages of SaPEP. (**c**) Fourier Shell Correlation (FSC) curves for the consensus 3D map of SaPEP as well as for the different local 3D maps (bodies) of SaPEP. The two dotted horizontal lines represent FSC=0.143 and 0.5. FSC=0.143 is used as a cutoff to determine the resolutions for the different reconstructions. (**c, d**) Local resolution of the consensus 3D map. Two views rotated by 45° are represented. The color code (in Å) is indicated. (**f**) Same view as (**e**) for the composite 3D map of SaPEP. The 3 different local maps (bodies) shown in panels g to i are highlighted in a different color. (**g, h, i**) 3D maps of body 1, 2 and 3 colored by local resolution.

**Figure S17:**
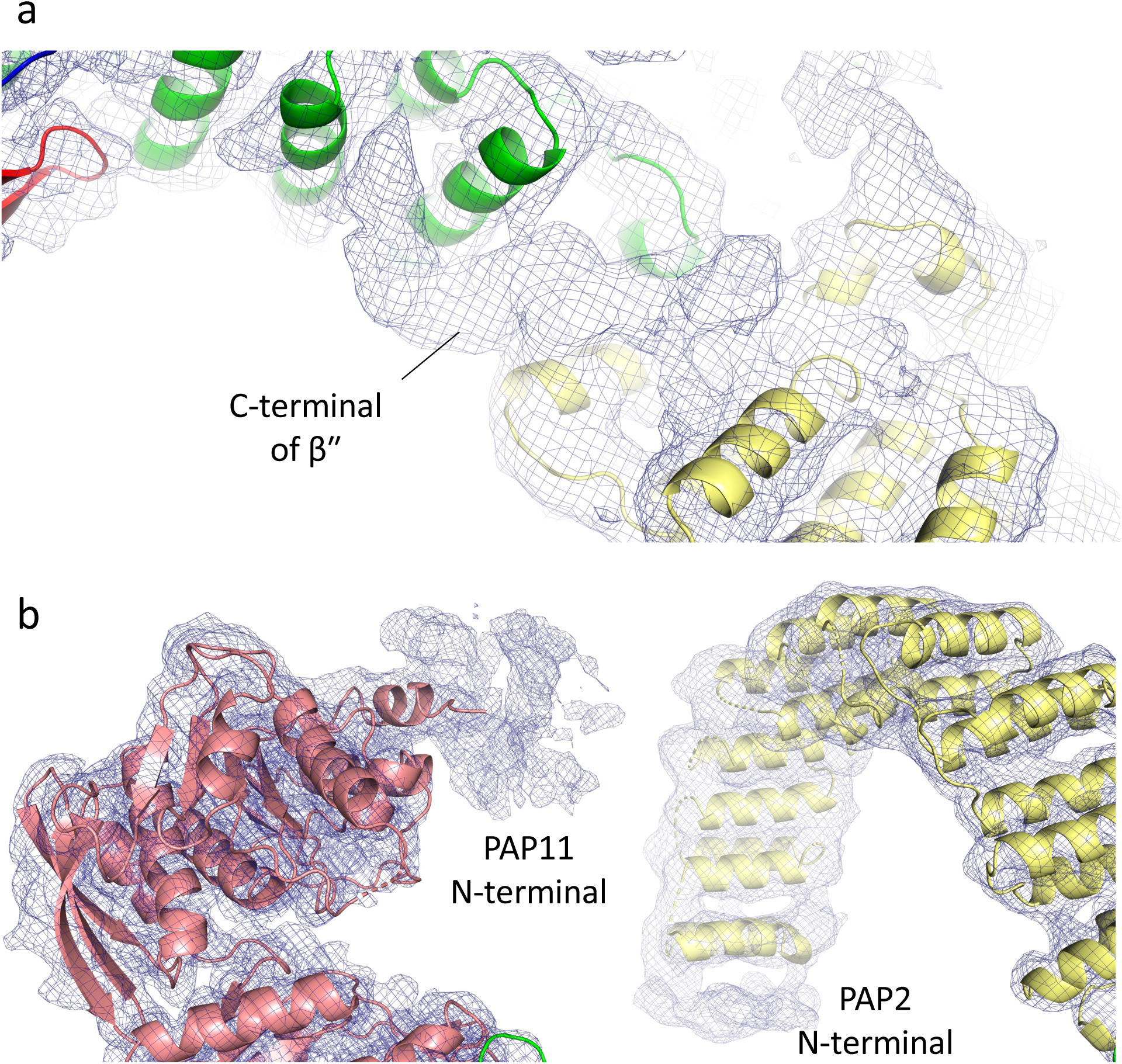
(**a**) Electron density corresponding to the β’’ C-terminal part not built between PAP1 and PAP2. (**b**) Electron density of the N-terminal part of PAP2 (51-103) and PAP11 (42-323).

**Tables S1:**
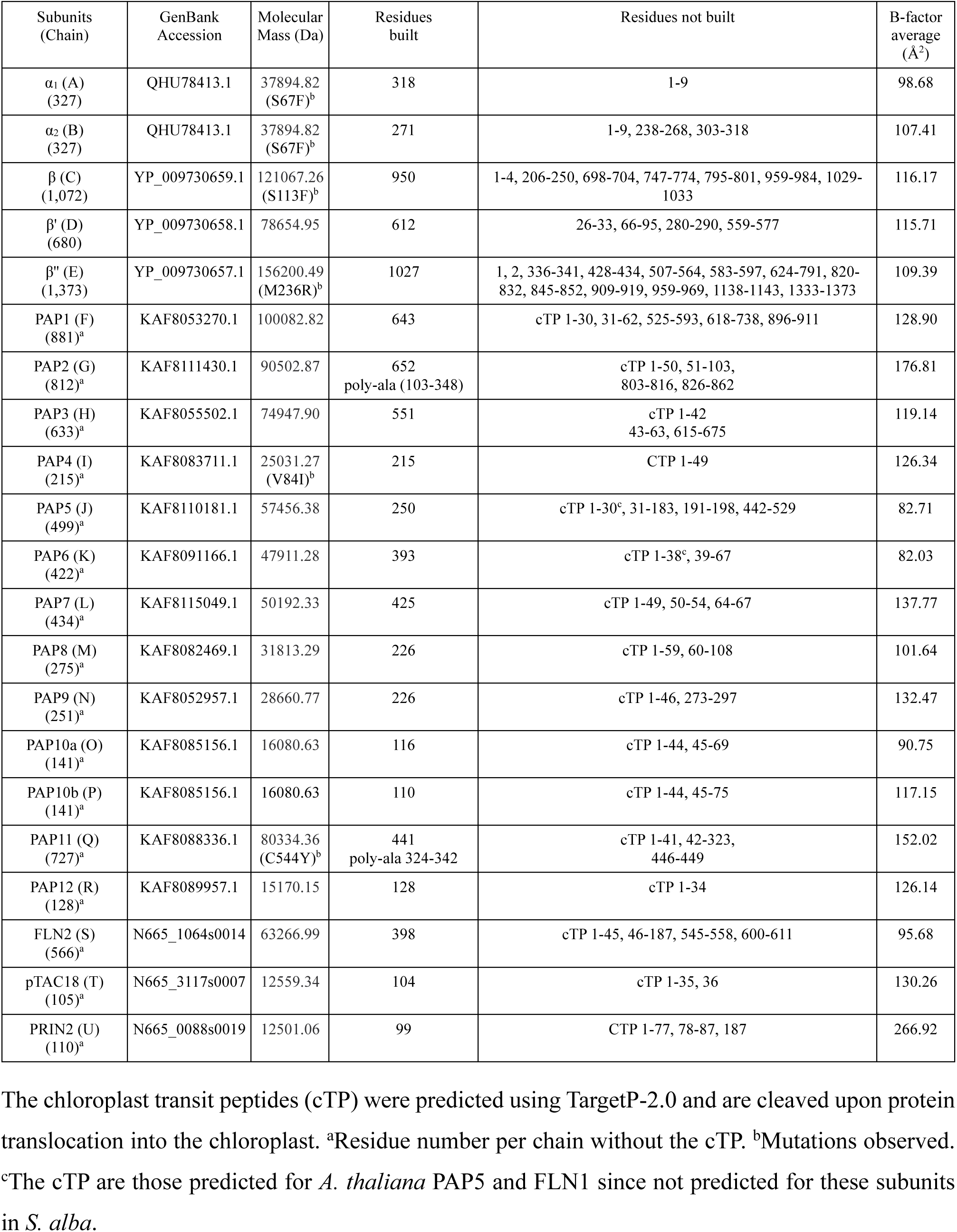
subunits of SaPEP: gene accession, molecular mass of each subunit. The residues built and not built in the model are indicated as well as the B-factor average of the chains.

**Table S2:** rms deviations (Å) between the catalytic subunits of SaPEP and those of Synechoccus elongatus and Synechocystis sp RNAPs (SeRNAP and SyRNAP).

| Subunits | SeRNAP (PDB entry: 8SYI) | SyRNAP (PDB entry: 8GZG) |
| --- | --- | --- |
| $\alpha_1$ (327)<sup>a</sup> | 1.806 (213)<sup>b</sup> | 1.513 (210)<sup>b</sup> |
| $\alpha_2$ (271)<sup>a</sup> | 1.912 (204)<sup>b</sup> | 1.740 (181)<sup>b</sup> |
| $\beta$ (950)<sup>a</sup> | 2.061 (858)<sup>b</sup> | 1.788 (860)<sup>b</sup> |
| $\beta'$ (612)<sup>a</sup> | 2.346 (513)<sup>b</sup> | 1.803 (510)<sup>b</sup> |
| $\beta''$ (1027)<sup>a</sup> | 2.490 (693)<sup>b</sup> | 2.354 (672)<sup>b</sup> |
| PAP12 ($\omega$) (128)<sup>a</sup> | 1.008 (54)<sup>b</sup> | 1.431 (54)<sup>b</sup> |

<sup>a</sup>Residue number per chain. <sup>b</sup>Residues aligned between the subunits

**Table S3:** data collection statistic of PRIN2.

|  |  |
| --- | --- |
| Resolution range (Å) | 37.27-1.74<br>(1.79-1.74) |
| Space group | P4<sub>1</sub> |
| Unit cell parameters (Å, °) | a, b= 37.27 c= 101.00 |
| Molecules in au | 1 |
| Number of total reflections | 153,775 (9,272) |
| Unique reflections | 14,084 (1,041) |
| Average multiplicity | 10.91 (8.91) |
| Data completeness (%) | 99.8 (99.0) |
| R<sub>merge</sub> (%) | 7.8 (73.1) |
| <I/$\sigma$(I)> | 20.55 (3.23) |
| CC(1/2) (%) | 99.9 (92.7) |

$R_{\text{merge}} = \frac{\sum \sum |I_i - I_m|}{\sum \sum I_i}$ , where $I_i$ is the intensity of the measured reflection and $I_m$ is the mean intensity of this reflection. Values indicated in parentheses correspond to the statistics in the highest resolution shell.

**Table S4:** refinement statistic of PRIN2.

|  |  |
| --- | --- |
| Resolution (Å) | 37.27-1.74<br>(1.88-1.74) |
| R<sub>cryst</sub> (σ<sub>F</sub>=0) (%) | 16.94 (20.03) |
| R<sub>free</sub> (σ<sub>F</sub>=0) (%) | 21.15 (23.16) |
| Number of atoms | 1009 |
| Water molecules | 77 |
| B average (Å<sup>2</sup>) | 51.82 |
| Rmsd bonds (Å) | 0.005 |
| Rmsd angle (°) | 0.769 |
| Ramachandran favored (%) | 99.12 |
| Ramachandran disallowed (%) | 0.0 |

Values indicated in parentheses correspond to the statistics in the highest resolution shell.
$R_{\text{cryst}} = \sum ||F_{\text{obs}}| - |F_{\text{calc}}|| / \sum |F_{\text{obs}}|$ . $R_{\text{free}}$ (Brünger, 1992) is the same as $R_{\text{cryst}}$ but calculated for 5% data omitted from the refinement.

**Table S5:**
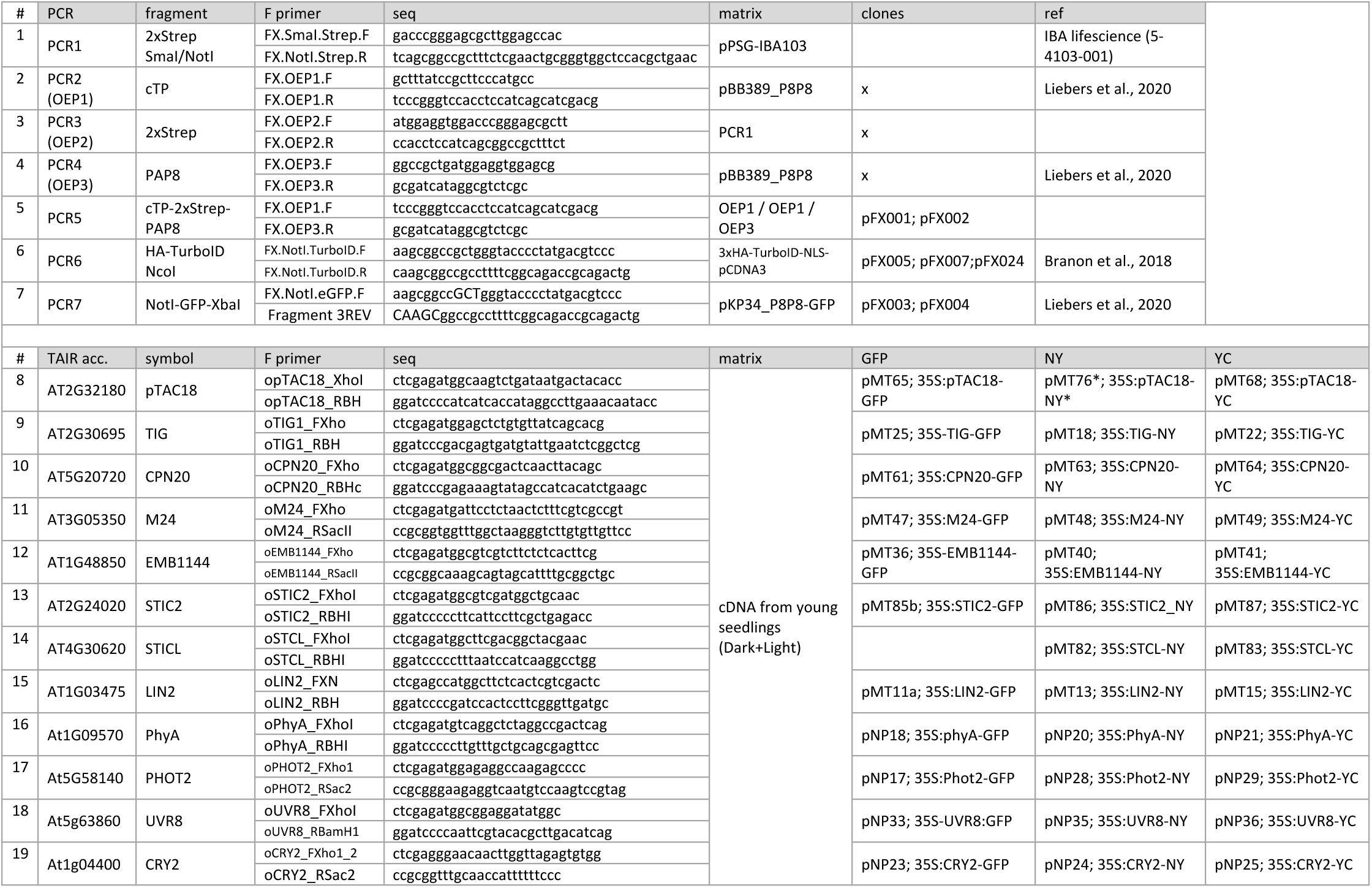
PCR-based cloning overview and corresponding primers.

**Table S6:** proteins from the photosynthetic complexes found enriched in Light versus dark.

| Locus Identifier | Gene Model Name | Symbol | cat | rank AP07L | rank AP24L | rank PL07L | rank PL24L |
| --- | --- | --- | --- | --- | --- | --- | --- |
| AT1G61520 | AT1G61520.1 | LHCA3 | 3 | 145 | 182 | 2037 | 1899 |
| AT1G31330 | AT1G31330.1 | PSAF | 3 | 276 | 200 | 600 | 531 |
| AT3G47470 | AT3G47470.1 | LHCA4 | 3 | 427 | 419 | 1609 | 1654 |
| AT1G30380 | AT1G30380.1 | PSAK | 3 | 549 | 408 | 1154 | 1347 |
| AT3G54890 | AT3G54890.1 | LHCA1 | 3 | 582 | 492 | 1244 | 1626 |
| AT1G55670 | AT1G55670.1 | PSAG | 3 | 611 | 463 | 1415 | 1314 |
| AT1G03130 | AT1G03130.1 | PSAD-2 | 3 | 784 | 897 | 254 | 260 |
| ATCG00340 | ATCG00340.1 | PSAB | 3 | 1093 | 1402 | 478 | 464 |
| AT4G28750 | AT4G28750.1 | PSAE-1 | 3 | 1092 | 1411 | 782 | 1052 |
| AT1G08380 | AT1G08380.1 | PSAO | 3 | 1090 | 1450 | 740 | 518 |
| AT3G61470 | AT3G61470.1 | LHCA2 | 3 | 1091 | 1787 | 1871 | 1274 |
| ATCG00680 | ATCG00680.1 | PSBB | 4 | 31 | 57 | 1294 | 969 |
| ATCG00280 | ATCG00280.1 | PSBC | 4 | 66 | 74 | 1124 | 802 |
| ATCG00020 | ATCG00020.1 | PSBA | 4 | 70 | 119 | 856 | 710 |
| ATCG00270 | ATCG00270.1 | PSBD | 4 | 99 | 99 | 1087 | 786 |
| AT1G29930 | AT1G29930.1 | CAB1 | 4 | 162 | 177 | 1843 | 2069 |
| AT1G44575 | AT1G44575.1 | NPQ4 | 4 | 238 | 330 | 710 | 578 |
| AT4G10340 | AT4G10340.1 | LHCB5 | 4 | 371 | 356 | 966 | 1061 |
| AT1G71500 | AT1G71500.1 | PSB33 | 4 | 354 | 381 | 844 | 826 |
| AT5G54270 | AT5G54270.1 | LHCB3 | 4 | 447 | 384 | 1822 | 1699 |
| AT1G15820 | AT1G15820.1 | LHCB6 | 4 | 515 | 364 | 1476 | 1257 |
| AT2G05070 | AT2G05070.1 | LHCB2.2 | 4 | 538 | 448 | 1281 | 1145 |
| AT5G01530 | AT5G01530.1 | LHCB4.1 | 4 | 552 | 466 | 1551 | 1270 |
| AT3G08940 | AT3G08940.2 | LHCB4.2 | 4 | 589 | 443 | 1173 | 1179 |
| AT3G50820 | AT3G50820.1 | PSBO2 | 4 | 654 | 566 | 511 | 529 |
| AT1G06680 | AT1G06680.1 | PSBP-1 | 4 | 1085 | 1234 | 366 | 437 |
| AT1G79040 | AT1G79040.1 | PSBR | 4 | 1086 | 1276 | 752 | 779 |
| AT2G34430 | AT2G34430.1 | LHB1B1 | 4 | 551 | 2196 | 2148 | 2004 |
| AT4G05180 | AT4G05180.1 | PSBQ-2 | 4 | 1087 | 1794 | 2582 | 1065 |
| AT5G66570 | AT5G66570.1 | PSBO1 | 4 | 1088 | 1939 | 584 | 2594 |
| ATCG01090 | ATCG01090.1 | NDHI | 5 | 254 | 335 | 442 | 445 |
| ATCG00420 | ATCG00420.1 | NDHJ | 5 | 392 | 867 | 109 | 85 |
| AT5G58260 | AT5G58260.1 | NdhN | 5 | 1084 | 1163 | 273 | 188 |
| AT4G37925 | AT4G37925.1 | NdhM | 5 | 1083 | 1215 | 52 | 59 |
| ATCG00430 | ATCG00430.1 | NdhK/PSBG | 5 | 1089 | 1222 | 68 | 64 |
| AT4G23890 | AT4G23890.1 | NdhS | 5 | 1082 | 1398 | 97 | 136 |
| ATCG00130 | ATCG00130.1 | ATPF | 6 | 107 | 89 | 547 | 500 |
| AT4G04640 | AT4G04640.1 | ATPC1 | 6 | 185 | 148 | 1018 | 1071 |
| AT4G09650 | AT4G09650.1 | ATPD | 6 | 141 | 247 | 516 | 443 |
| ATCG00120 | ATCG00120.1 | ATPA | 6 | 327 | 255 | 1279 | 1075 |
| ATCG00480 | ATCG00480.1 | PB | 6 | 321 | 270 | 549 | 581 |
| AT4G32260 | AT4G32260.1 | PDE334 | 6 | 1081 | 295 | 2521 | 1161 |
| AT4G03280 | AT4G03280.1 | PETC | 7 | 402 | 1340 | 156 | 137 |
| ATCG00540 | ATCG00540.1 | PETA | 7 | 616 | 1343 | 893 | 1059 |
| ATCG00720 | ATCG00720.1 | PETB | 7 | 1080 | 1805 | 2590 | 469 |
| AT4G02510 | AT4G02510.1 | TOC159 | 9 | 425 | 241 | 795 | 880 |
| AT2G24820 | AT2G24820.1 | TIC55-II | 9 | 1079 | 1433 | 330 | 314 |
| AT4G33350 | AT4G33350.1 | Tic22-IV | 9 | 726 | 2178 | 1417 | 1597 |

## Material and Methods

### Purification of the chloroplasts from *Sinapis alba*

Seven-day-old cotyledons of *S. alba* seedlings were collected (∼ 400 g in total). 100 g fresh material was blended with 200 mL homogenization buffer (50 mM HEPES-KOH pH 8.0, 0.3 M sorbitol, 5 mM MgCl2, 2 mM EDTA, 0.3 mM DTT) with short pulses (3 × 3 s). The homogenized leaf tissue was preliminary filtered using a 50 µm nylon mesh. Double extraction was performed to optimize the chloroplast concentration. The mixture was centrifuged at 6,084*g* for 6 min at 4°C. The pellet was resuspended in the homogenization buffer and pottered before loading into a three-step Percoll gradient (20%-40%-80%). Intact chloroplasts were collected from the 40% interphase and centrifuged at 15,000*g* for 15 min at 4°C. Finally, the chloroplasts were lysed by resuspending the pellet into the lysis buffer (50 mM Tris HCl pH 7.6, 10 mM NaF, 25% (*w/v*) glycerol, 1% (*w/v*) Triton X-100, 4 mM EDTA, 1 mM DTT) at equal volume, frozen in liquid nitrogen and stored at -80°C before use.

### Purification of the PEP from *Sinapis alba*

Chloroplast membranes were broken into the lysis buffer using a potter and centrifuged at 15,000 *g* for 1 h at 4°C. The supernatant was collected. The PEP purification was performed using Fast Protein Liquid Chromatography in a three-step process. The supernatant was loaded onto a heparin column (2 mL resin). The column was washed with 50 mM TrisHCl pH 7.6, 80 mM (NH4)2SO4, 5 mM MgCl2, 10% (*w/v*) glycerol, 0.1% (*v/v*) Triton X-100, 1 mM DTT. Proteins were eluted in one step with 5 column bed volume of 50 mM TrisHCl pH 7.6, 1.2 M (NH4)2SO4, 5 mM MgCl2, 10% (*w/v*) glycerol, 0.01% (*v/v*) Triton X-100, 1 mM DTT. The protein solution was diluted in 50 mM TrisHCl pH 7.6, 5 mM MgCl2, 10% (*w/v*) glycerol, 1 mM DTT, and loaded onto the 5/50 MonoQ™ anion exchange column. After washing, the proteins were eluted using a linear 0 – 1 M NaCl gradient in 50 mM TrisHCl pH 7.6, 5 mM MgCl2, 10% (*w/v*) glycerol, 1 mM DTT. The fractions containing the PEP revealed by Western blotting were pooled and concentrated on a 100-kDa cutoff membrane at 2,500*g*, 4°C before loading a Superose™ 6 10/300 GL column. The proteins were eluted in 50 mM TrisHCl pH 7.6, 5 mM MgCl2, 500 mM NaCl, 8% (*w/v*) glycerol, 1 mM DTT. Selected fractions containing PEP after Western Blot analysis were pooled, concentrated, before frozen in liquid nitrogen and stored at -80°C.

### Purification of the PEP from *Arabidopsis thaliana*

Plant material was broken in a mortar, pottered in lysis buffer (100 mM TrisHCl pH 8, 150 mM NaCl, 2 mM KCl, 5 mM MgCl2, 10% (*w/v*) glycerol, DNase, benzonase, anti-protease, 0.2% (*v/v*) DDM, 1% (*v/v*) Triton X-100, 1mM DTT) and centrifuged at 15,000*g* after 1 hour incubation at 4°C. The supernatant was collected and loaded on Strep-Tactin®XT column. The column was washed with 100 mM TrisHCl pH 8, 5 mM MgCl2, 2 mM KCl, 150 mM NaCl, 10% (*w/v*) glycerol, 0.01% (*v/v*) Triton X-100, 0.05% (*v/v*) DDM. The proteins were eluted with 100 mM TrisHCl pH 8, 5 mM MgCl2, 2 mM KCl, 150 mM NaCl, 50 mM biotin, 10% (*w/v*) glycerol, 0.01% (*v/v*) Triton X-100, 0.05% (*v/v*) DDM and concentrated using a centrifugal filter unit with a 100 kDa cut-off at 2,500*g*, 4°C. The concentrated solution was loaded onto the Superose™6 10/300 GL column in 50 mM TrisHCl pH 8, 5 mM MgCl2, 2 mM KCl, 150 mM NaCl, 8% (*w/v*) glycerol, 1 mM DTT. Selected fractions by western blot analysis were pooled, concentrated, frozen in liquid nitrogen for further uses.

### Negative stain EM of AtPEP

4 µL of purified PEP 0.01-0.1 mg/mL (concentration estimated from chromatogram) were absorbed to the clean side of a carbon film on mica, stained with sodium silico tungstate (SST) at 2% in distilled water (pH 7-7.5) and transferred to a 400-mesh copper grid. The images were taken with defocus values between 1.2 and 2.5 μm on a Tecnai 12 LaB6 electron microscope at 120 kV accelerating voltage using CCD Camera Gatan Orius 1000. The image processing was done in RELION 3.1.2. CTF estimation was done with CTFFind-4.1. An initial set of particles (box size of 256 pixels, sampling of 2.2 Å/pixel) was obtained by manual picking, after 2D classification. The best looking 2D class averages were used as references for an autopicking. A set of particles (box size of 256 pixels, sampling of 2.2 Å/pixel, mask diameter 400 Å) was obtained by autopicking with a gaussian blob. After 2D classification the best looking 2D class averages were clearly identifiable. The 3D envelope modeling was calculated at 31 Å resolution.

### Cryo-EM data collection of SaPEP

3.5 µL of purified SaPEP were applied to 1.2/1.3 holey carbon grids (Quantifoil MicroTools GmbH, Germany) coated with graphene oxide using a protocol adapted from Patel et al., 2021 and similar. Grids were plunged frozen in liquid ethane with a Vitrobot Mark IV (Thermo Fisher Scientific) (4 s blot time, blot force 0). The sample was observed at the beamline CM01 of the ESRF (Grenoble, France) (Kandiah et al., 2019) with a Titan Krios G3 (Thermo Fischer Scientific) at 300 kV equipped with an energy filter (Bioquantum LS/967, Gatan Inc, USA) (slit width of 20 eV). Movies were recorded automatically with a K3 direct detector (Gatan Inc., USA) with EPU (Thermo Fischer Scientific). Movies were recorded in two sessions on the same grid (3022+11493 movies) for a total exposure time of 4 s with 40 frames per movie and a total dose of ∼51 e^−^/Å^2^. The magnification was 81,000x (1.06 Å/pixel at the camera level). The defocus of the images was varied between −1.0 and −2.4 μm.

### Cryo-EM image processing

The movies were first drift-corrected with motioncor2 (Zheng *et al.,* 2017). CTF estimation was done with GCTF (Zhang et al., 2016). The remaining image processing was performed with RELION 4 (Kimanius e*t al*., 2021). From the first dataset of 3022 micrographs, 855 were selected manually based on the presence of particles (Figure S16). An initial set of particles was obtained by auto-picking with Laplacian of Gaussian. After two rounds of 2D classification, the particles from the best looking 2D class averages were selected and used to train Topaz (Bepler *et al*. 2019) on the 3022 micrographs. The picked particles were again classified by 2D classification to remove false positives and a first 3D reconstruction was obtained by using as a reference a previously determined 3D reconstruction from negative stain data (EMD-14571). Following 3D classification (C1 symmetry, no mask, 3 classes), a 3D reconstruction at 4.1 Å resolution was obtained from 23419 particles for the first dataset of 3022 micrographs. In order to improve the resolution, a second larger dataset (11493 micrographs) was added and processed in an identical way leading to an extra 82377 particles. The particles from both datasets were merged and processed in steps in the order particle polishing, 3D refinement, refinement of CTF parameters, 3D refinement. This sequence of steps was repeated two more times until no further improvements in resolution was observed. The final consensus map (EMD-50711) reached the resolution of 3.0 Å resolution. As the 3D map displays local variability in various places, a multibody refinement was run with 4 bodies, the larger one (body 4 – EMD-50722) includes the best resolved part of the consensus map and did not improve significantly while the other 3 bodies are centered on the flexible regions. This approach substantially improved the interpretability of the variable regions. These 3 new 3D maps (body 1: EMD-50718, body 2: EMD-50719 and body 3: EMD-50720) were used to build missing regions of the atomic models but were still displaying variability. In a final attempt to better resolved the missing regions, the various bodies were isolated by signal subtraction of the rest of the particle and 3D classified in steps until no further improvements in resolvability was observed. Body 1 (EMD-50718), body 2 (EMD-50719) and body 3 (EMD-50720) were reconstructed to 3.3 Å (from 78 218 particles), 3.9 Å (from 22 387 particles) and 3.6 Å (from 38 402 particles) respectively. Using the final 3D maps from bodies 1 to 4, a composite map (EMD-19877) was generated in chimera (Pettersen, E. F. *et al*. 2004). All the resolutions were determined by Fourier Shell Correlation (FSC) at 0.143 between two independent half 3D maps. The local resolution was calculated with *blocres* (Cardone *et al*., 2013) and found to be between 2.5 and 6.5 Å. The final 3D map was sharpened with DeepEMhancer for display purpose (Sanchez-Garcia *et al*., 2021).

### Model building and refinements

The model of SaPEP was built and refined using COOT (Emsley & Cowtan, 2004) and Phenix (Liebschner et al., 2019) respectively in the global map calculated at 3 Å resolution. The provided coordinates were placed by rigid-body and the model refined using energy minimization and simulated annealing. Cycles of rebuilding and refinements were then performed by combining with models from AlphaFold (Jumper et al., 2021) and *E. coli* RNAP structures. Three local refinements were performed in corresponding local maps before building of PAP2, PAP11, and some parts of β’, β’’ and PAP3. Each model was then refined in the corresponding local cryo-EM. The four models obtained were then merge after analysis with COOT, and the structure of PRIN2 solved by X-ray crystallography was added. The composite model was then refined in the composite electron density map and rebuilt before a last cycle of refinements. The atomic coordinates and cryo-EM map of the final model were deposited in the PDB (Entry: 9EPC).

Cryo-EM electron density map analyses revealed five main blobs of electron density not filled with models corresponding to the two first α-helices of PAP2, the N-terminal of PAP11, the C-terminal of β’’, the α-helix in α2-CTD and the SMBH5, 6 and 7 of β’’ (Table S1 and Figures S2, S17). The other missing parts of the model are also given in Table S1. The model statistics are given in Table 2. The superimpositions were calculated using Superpose (Krissinel & Henrick, 2004) from CCP4 (CCP4, 1994) and the structural comparisons performed using DALI (Holm, 2020).

### Overexpression and crystallization of PRIN2

PRIN2 synthetic gene was cloned in pET28a between the NdeI and HindIII restriction site with a 6His-Tag and a thrombine cleavage site. PRIN2 was overexpressed in *E. coli* DE3 strain in LB with 50 mg/mL kanamycin. Cells were grown overnight in 50 mL of LB with antibiotics at 37°C. 1 L of LB (with antibiotics) was then inoculated with the first culture to reach an initial OD600 of 0.1. Growth was continued at 37°C. When the OD600 reached 0.6, the temperature was decreased to 16°C and 0.5 mM of isopropyl β-D-1-thiogalactopyranoside was added. After an overnight induction, bacteria were harvested at 5,500g, for 25 min, at 4°C. The cell pellet was resuspended in 15 mL of lysis buffer (50 mM Tris HCl, pH 8.0, 0.5 M NaCl, 20 mM imidazol, 20 mM β-mercaptoethanol) containing a Complete Protease inhibitor Cocktail tablet (Roche). The lysate was centrifuged at 15,000g, for 40 min, at 4°C. The purification was performed at room temperature. The supernatant was applied onto a NiNTA column in 50 mM Tris HCl, pH 8.0, 0.5 M NaCl, 20 mM imidazol, 20 mM β-mercaptoethanol. Column was washed with 50 mM Tris HCl, pH 8.0, 0.5 M NaCl, 20 mM imidazol, 20 mM β-mercaptoethanol and proteins were eluted in four steps in a buffer containing 50 mM Tris HCl, pH 8.0, 0.1 M NaCl, 20 mM β-mercaptoethanol with increasing imidazol concentrations (50, 100, 200 and 300 mM). Then the eluate was dyalised in 50 mM TrisHCL, 500 mM NaCl, 20 mM β-mercaptoethanol and concentrated with an Amicon Ultra 4 mL centrifugal filter and a 10 kDa membrane cut-off before loading on a HiLoad 16/60 Superdex 75 and then eluted with 10 mM Tris HCl, pH 8.0, 50 mM NaCl, 5 mM DTT. The fractions containing the pure protein were pooled and concentrated for further experiments or frozen at -20% with 50 % (v/v) glycerol.

^15^N,^13^C-PRIN2 was expressed in minimum media M9 supplemented with ^15^NH4Cl, ^13^C-glucose and antibiotics. Briefly, 5 mL of LB were inoculated with *E. coli* DE3 stock glycerol overexpressing PRIN2. After 10h of growing, 1 mL were added to 100 mL of minimum media supplemented as described above. After 1 night growing, when OD600 was close to 2, the overnight culture was centrifuged to inoculate 1 L of minimum media M9 supplemented with ^15^NH4Cl and ^13^C-glucose and antibiotics. Cell growth, overexepression and purification followed the procedure described above for PRIN2.

### Crystallization, data collection and structure resolution

PRIN2 at 5 and 10 mg/mL in 10 mM Tris HCl, pH 8.0, 50 mM NaCl was subjected to crystallization using the sitting-drop vapor-diffusion technique and the high throughput crystallization facility at the EMBL, Grenoble, at 4°C. Crystallization hits were optimized using Limbro plates, at 20°C. Crystals of PRIN2 were obtained in several crystallization conditions. Finally, crystals were grown in 6% PEG 8K, 0.2 M magnesium acetate, 0.1 M sodium cacodylate pH 6.5, 25% glycerol, at 4°C, and a protein concentration of 15 mg/mL. Diffraction data were collected on BL13 (Juanhuix et al., 2014) at the ALBA synchrotron, Cerdanyola del Vallès, Spain, 100 K, using a PILATUS 6M detector and one crystal. Diffraction data (Table S3) were processed using XDS and the datasets from two crystals were scaled using XDS (Kabsch, 2010).

### 2D-NMR, phasing and structure refinement

For assignment of PRIN2, 500 μM of ^15^N,^13^C-PRIN2 in a 90:10 H2O:D2O 10 mM Tris, pH 8.0, 250 mM NaCl was used and heteronuclear 3D Best-TROSY-HNCO, Best-TROSY-HNCACO, Best-TROSY-HNCA, Best-TROSY-HNCOCA, Best-TROSY-HNCACB (Solyom *et al*, 2013), Best-TROSY-HNCOCACB, Best-TROSY-HNCOCANH and N-NOESY-HSQC experiments were recorded at 35°C on Bruker ADVANCE III HD spectrometer operating at ^1^H frequency of 950 MHz equipped with a triple resonance pulsed field gradient cryoprobe. All experiments can be found in the NMRlib library (Favier and Brutscher, 2019). [^15^N,^1^H]-TRACT was recorded to estimate the overall correlation time of the protein, which is expected to be a monomer. 1D-NMR was used to characterize PRIN2 as a well-folded protein suitable for crystallization. Data were deposited in the BMRB with the ID 52532.

Phasing by molecular replacement using Arcimboldo (Rodriguez et al., 2012) from CCP4 (CCP n°4, 1994) was performed with four α-helices as model. The model was after partially automatically built using Arp/Warp (Chojnowski et al., 2019). The refinements and rebuilding were next done using PHENIX (Liebschner et al., 2019) and COOT (Emsley & Cowtan, 2004), respectively. The water molecules were added using PHENIX in the last stages of the refinement. Refinement statistics are summarized in Table S4. Atomic coordinates and X-ray data were deposited in the PDB with the accession number 9EPT.

### Mass spectrometry-based proteomic characterization of PEP-enriched fraction

Proteins were solubilized in Laemmli buffer and stacked in the top of a 4-12% NuPAGE gel (Invitrogen). After staining with R-250 Coomassie Blue (Biorad), proteins were digested in-gel using trypsin (modified, sequencing purity, Promega) as previously described (Casabona et al. 2013), except that Tris(2-carboxyethyl)phosphine hydrochloride was used instead of dithiothreitol. The resulting peptides were analyzed by online nanoliquid chromatography coupled to MS/MS (Ultimate 3000 RSLCnano and Q-Exactive HF, Thermo Fisher Scientific). For this purpose, the peptides were sampled on a precolumn (300 μm x 5 mm PepMap C18, Thermo Scientific) and separated in a 75 μm x 250 mm C18 column (Aurora Generation 3, 1.7μm, IonOpticks) using a 60 min gradient. The MS and MS/MS data were acquired by Xcalibur version 2.9 (Thermo Fisher Scientific). The mass spectrometry proteomics data have been deposited to the ProteomeXchange Consortium via the PRIDE (PMID: 34723319) partner repository with the dataset identifier PXD0X.

Peptides and proteins were identified by Mascot (version 2.8.0, Matrix Science) through concomitant searches against the Uniprot database (*Arabidopsis thaliana* taxonomy, 20230922 download), and a homemade database containing the sequences of classical contaminant proteins found in proteomic analyses (keratins, trypsin, etc.). Trypsin/P was chosen as the enzyme and two missed cleavages were allowed. Precursor and fragment mass error tolerances were set at respectively 10 and 20 ppm. Peptide modifications allowed during the search were: Carbamidomethyl (C, fixed), Acetyl (Protein N-term, variable) and Oxidation (M, variable). The Proline software (Bouyssié et al., 2020) was used for the compilation, grouping and filtering of the results (conservation of rank 1 peptides, peptide length ≥ 6 amino acids, false discovery rate of peptide-spectrum-match identifications < 1% (Couté et. al., 2020), unique interpretation of spectrum, and minimum of one specific peptide per identified protein group). Proline was then used to perform a MS1-based label-free quantification of the identified protein groups. For each protein groups, iBAQ values (Schwanhäusser et al., 2011) were computed from MS1 intensities of specific and razor peptides. Proteins identified in the contaminant database, and proteins bearing a biotin attachment domain (PFAM accession PF00364) or associated to proteins bearing a biotin attachment domain were discarded.

### Mass spectrometry-based proteomic characterization of PAP8 interactome and proxisome

For each condition (WT-Ctrl, tagged stP8 short linker, tagged stλP8 long linker, all in dark or light), four biological replicates were prepared. For interactome analysis, proteins were solubilized in Laemmli buffer and stacked in the top of a 4-12% NuPAGE gel (Invitrogen). After staining with R-250 Coomassie Blue (Biorad), proteins were digested in-gel using trypsin (modified, sequencing purity, Promega) as previously described (Casabona et al., 2013). For proxisome analysis, proteins were digested on beads as previously described (Mair et al, 2019). The resulting peptides were analyzed by online nanoliquid chromatography coupled to MS/MS (Ultimate 3000 RSLCnano and Q-Exactive HF, Thermo Fisher Scientific). For this purpose, the peptides were sampled on a precolumn (300 μm x 5 mm PepMap C18, Thermo Scientific) and separated in a 75 μm x 250 mm C18 column (Reprosil-Pur 120 C18-AQ, 1.9 μm, Dr. Maisch) using a 120 min gradient. The MS and MS/MS data were acquired by Xcalibur version 2.8 or 2.9 (Thermo Fisher Scientific). The mass spectrometry proteomics data have been deposited to the ProteomeXchange Consortium via the PRIDE (PMID: 34723319) partner repository with the dataset identifier PXD0X.

Peptides and proteins were identified by Mascot (version 2.7.1, Matrix Science) through concomitant searches against the TAIR database (version 10.0), a homemade database containing the sequences of tagged PAP8, a homemade database containing the sequences of classical contaminant proteins found in proteomic analyses (keratins, trypsin, etc.), and their corresponding reversed databases. Trypsin/P was chosen as the enzyme and two missed cleavages were allowed. Precursor and fragment mass error tolerances were set at respectively at 10 and 20 ppm. Peptide modifications allowed during the search were: Carbamidomethyl (C, fixed), Acetyl (Protein N-term, variable) and Oxidation (M, variable). The Proline software (Bouyssié et al., 2020) was used for the compilation, grouping and filtering of the results (conservation of rank 1 peptides, peptide length ≥ 6 amino acids, peptide-spectrum-match score ≥ 25 allowing to reach a false discovery rate < 1%, unique interpretation of spectrum, and minimum of one specific peptide per identified protein group). Proline was then used to perform a MS1-based label-free quantification of the identified protein groups based on specific and razor peptides.

Statistical analysis was performed using the ProStaR software (Wieczorek et al., 2017) based on the quantitative data obtained with the four biological replicates analyzed per condition. For each pairwise comparison, proteins identified in the contaminant database, proteins identified by MS/MS in less than two replicates of one condition, and proteins quantified in less than four replicates of one condition were discarded. After log2 transformation, abundance values were normalized either using condition-wise the variance stabilizing normalization (vsn) method (for stP8 versus WT comparisons), or on the abundance value of tagged PAP8 (for dark versus light), before missing value imputation (SLSA algorithm for partially observed values in the condition and DetQuantile algorithm for totally absent values in the condition). Statistical testing was conducted with limma, whereby differentially expressed proteins were selected using a log2(Fold Change) cut-offs of 1.6 and 1, for respectively WT versus tagged PAP8 and dark versus light comparisons, and a p-value cut-off of 0.01, allowing to reach false discovery rates inferior to 2.5% according to the Benjamini-Hochberg estimator. Proteins found differentially abundant but identified by MS/MS in less than two replicates or detected in less than four replicates in the condition in which they were found to be more abundant were manually invalidated (p-value = 1).

### Biological materials

*Arabidopsis thaliana* seeds; *pap8-1*: SALK_024431 (N524431); and Col-0: SALK_6000, were obtained from The European Arabidopsis Stock Centre NASC. *E. coli*, DH5α strain (lacZ-ΔM15 Δ(lacZYA-argF) U169 recA1 endA1 hsdR17(rK-mK+) supE44 thi-1 gyrA96 relA1) was used for cloning. *Agrobacterium tumefaciens* strain C58C1 pMP90 was used for transgenesis.

### Cloning

Minipreps were performed using Qiagen kits and DNA in-gel purification using Monarch^R^ (New England Biolabs). For cloning, PCR fragments were amplified using Phusion^TM^ High-Fidelity DNA Polymerase (Thermo Scientific). Gotaq^TM^ polymerase (Promega) was used for A tailing. DNA ligations were done using T4 DNA ligase (New England Biolabs, (NEB)). Bacterial transformation was performed in competent heat shock *E. coli* strain DH5α (Invitrogen). sP8 (pFX002) was generated by Overlap Extension PCR (OEP); fragment XhoI-cTP-2xStrep-PAP8-XbaI (Table S5 for the list of primers) cloned into pGEM-T easy (Promega) producing pFX001 then inserted XhoI XbaI in pBB389 (Liebers et al., 2020). stPAP8 (pFX007): the fragment NotI-HA-TurboID-NotI amplified from 3xHA-TurboID-NLS_pCDNA3, cloned into pGEM-T easy yielded pFX005; then the NotI fragment was ligated in pFX002 giving pFX007. stλP8 (pFX024): NcoI-PAP8-XbaI fragment was cloned in pGemT easy giving pFX020, then transferred to pBB389 to give pFX023. The NotI fragment from pFX005 was ligated in pFX004 (pFX004 is cloned from pFX002 by addition of the fragment NotI-GFP-XbaI previously cloned in pGem-T giving pFX003) to generate pFX006; then the NcoI-HA-TurboID-NcoI fragment was ligated in pFX023 giving pFX024. All constructions were verified by restriction and sequencing. Coding sequences in Figure 5d and 6e, were amplified from cDNA prepared with germinating seedlings grown in the dark or light conditions as in the AP/PL experiment. PCR fragments cloned in the pCR^R^-bluntII-TOPO vector (Thermofisher). Vectors were sequenced then digested and corresponding inserts ligated in GFP and split YFP vectors (Liebers et al., 2020).

### Plant transformation

Thermo-competent *Agrobacterium tumefaciens* were transformed with binary plasmids containing transgene and positive colonies were screened under a mixture of antibiotics: rifampicin (bacterial strain selection), gentamycin (helper plasmid selection), and spectinomycin (binary vector selection). Strains were then used for floral dip infiltration of the significant genotypes using the infiltration Medium (2.2 g MS salts, 1 mL Gamborg’s 1000× B5 vitamins, 0.5% sucrose, 44 nM benzyl amino purine, 300 μL/L Silwet L-77) and *A. tumefaciens* concentration at around 2<OD600<3. Sporophytic lethal *pap8-1* was used as the progeny of a heterozygous plant; transgenic plants were then selected to carry the mutant allele *pap8-1* (yielding albino plants in the progeny) and the hygromycin antibiotic selection marker.

### Growth conditions

The parameters of the growth chamber were: day / night (16hrs / 8hrs); light intensity between 130-150 µmol.m^-2^.s^-1^; lamp reference (Neon 77 fluora (OSRAM) 18 W); temperature day / night (22°C / 18°C) and the relative humidity day / night (70% / 60%). The light intensity was assessed before each experiment using a PAR-FAR photometer (Apogee instruments, reference S2-141-SS). Seeds were sterilized by successive shaking treatments of bleach 3%, ethanol 90%, triton-X100 3 drops/ 50 mL (1 bath, 4 minutes) and absolute ethanol (4 baths, 15 seconds). Dried seeds were sown on a 1/2 MS media, MES (250 µM), Gamborg’s vitamin 1X, 0.8% agar and let imbibed 1 hour under light exposure. Seeds were stratified (5 days, dark, 4°C), light induced for germination (4-hour light, 22°C) and dark induced for skotomorphogenesis (4-day dark, 22°C). Afterward photomorphogenesis was triggered by transferring the seedling under light at 22°C. The dark condition corresponds to wrapping the plates in two layers of aluminum foil. When necessary, the plates were manipulated in a near-dark room with a 30 Watt Far-Red Led (pic at 730 nm +/-20 nm) using an intensity variator set at minimum. The use of spy plates guaranteed proper skotomorphogenic stage set at 10-12 mm etiolated hypocotyls.

### Protein sub-cellular localization

Transient expression and fluorescent microscopic analysis in onion cells and Arabidopsis seedlings were performed as previously described (Liebers et al., 2020) with few modifications; transient assay in onion cells (bulb sliced to ∼16 cm^2^) was conducted using the Biolistic PDS 1000/He Particle Delivery System (Biorad) (1100 psi, 10 cm traveling distance) with DNA onto 1 µm gold particles (Seashell Technology TM) following instructions. After 16 to 40 h in the dark at 24°C, the epidermis was peeled and observed by fluorescence microscopy with a Nikon AxioScope equipped with FITC filters and an AxioCam MRc camera. Pictures were acquired with the Nikon’s Zen software.

### Protein extraction and Western immuno-detection

Arabidopsis seedling (50 mg, approx. 100 seedlings) were collected and homogenized by tissue grinding in liquid nitrogen using a mortar in 100 µL of denaturing extraction buffer (DEB: Tris HCl 100 mM pH 6.8, Urea 8 M, EDTA/EGTA 10 mM, DTT 10 mM, protease inhibitor (Roche) 1 tablet/10 mL. The samples were centrifuged (10 min, 4°C, 9,300 g). The total soluble protein samples (TSP) were titrated by Bradford assay before mixing in Laemmli buffer (Tris HCl 100 mM pH6.8, Glycerol 10%, SDS 2%, DTT 50 mM, Bromophenol Blue 0.25%) and heated 10 min at 80°C. TSP were separated by SDS-PAGE and transferred on nylon membrane (Biorad). The membrane was blocked in TBS, Tween 0.1%, non-fat dry milk 5% w/v; then probed in TBS Tween 0.1%, with different primary antibodies against PAP8 (Liebers et al., 2020), or against 2xStrep (streptagII IBA, StrepMAB-classic ref# 2-1507-001). Membranes were washed (5 times, 5 min in a TBS-Tween 0.1%); secondary antibody, Goat anti-Rabbit conjugated with a HorseRadish Peroxidase was used at a dilution of 1/5,000. Signal was detected using a chemiluminescent substrate (Biorad, ECL kit).

### Proximity labeling

For the proximity labeling/biotrap and streptrap experiments, 180 µL of Arabidopsis seeds were sown on 121-cm^2^ square plates. The biotin powder was dissolved in DMSO at a 200X of the desired final concentration indicated in each experiment, then diluted in MilliQ water under shaking. The control without biotin corresponds to a mock treatment with DMSO alone. The seedlings were immersed with 70 mL of biotin/DMSO solution per plate during 8 hours. Seeds were initially sown on a nylon membrane laid down on the MS-agar medium. The nylon membrane was removed using forceps and washed twice with 500 mL of MilliQ water to release the seedlings and remove excess of biotin. The seedlings were collected and quickly dried on tissue paper and flash-frozen in liquid nitrogen. The seedlings were ground in liquid nitrogen for five minutes and the powder stored at -80°C.

### Affinity purifications

Approximately 5 mL of packed seedling were grinded in a mortar with liquid nitrogen for 4 min into a fine powder. This procedure yielded approximately 4.5 mg or 15.5 mg of total soluble proteins for seedling in skoto- or photomorphogenesis respectively. The whole process was performed at 4°C in a cold room. The biotrap affinity purification was performed as described by Mair et al. (2018). For the streptrap affinity purification, the powder was suspended in 4 mL of extraction buffer (TrisHCl 100 mM pH7.9, NaCl 50 mM, MgCl2 2 mM, KCl 2 mM, BSA 0.1%, 2 mM DTT, 1x Complete^TM^ protease inhibitor, PMSF 1 mM, NaF 10 mM, Triton 0,5%, 0.5% n-dodecyl-β-D-maltoside (DDM)). The extract was centrifuged (2000 *g*, 4°C, 10 min) and its supernatant centrifuged again (17,000 *g*, 4°C, 15 min). The clear supernatant was then transferred in a 5-mL low-binding tube containing 200 µL of Streptactin XT magnetic beads conjugated (IBA) previously washed with the extraction buffer. The protein extract was incubated 2 h on a rotor wheel. The beads were washed 5 times with the washing buffer I (TrisHCl 100 mM pH7.9, NaCl 50 mM, MgCl2 2 mM, KCl 2 mM, BSA 0.1%, 2 mM DTT, 1x Complete^TM^, PMSF 1 mM, NaF 10 mM, Triton 0,5%, DDM 0.1%) and 3 times with the washing buffer II (TrisHCl 100 mM pH7.9, NaCl 50 mM, MgCl2 2 mM, KCl 2 mM, 2 mM DTT, 1x Complete^TM^, PMSF 1 mM, NaF 10 mM, Triton 0,1%). Beads (20 µL) were boiled in 50-µL Laemmli buffer 2.5x final, 20 mM DTT for immunodetection of PAP8, GFP and silver nitrate staining. The rest of the beads was stored without the buffer at -80°C for MS analysis.

